# The Origin and Early Evolution of the Legumes are a Complex Paleopolyploid Phylogenomic Tangle closely associated with the Cretaceous-Paleogene (K-Pg) Boundary

**DOI:** 10.1101/577957

**Authors:** Erik J.M. Koenen, Dario I. Ojeda, Royce Steeves, Jérémy Migliore, Freek T. Bakker, Jan J. Wieringa, Catherine Kidner, Olivier Hardy, R. Toby Pennington, Patrick S. Herendeen, Anne Bruneau, Colin E. Hughes

## Abstract

The consequences of the Cretaceous-Paleogene (K-Pg) boundary (KPB) mass extinction for the evolution of plant diversity are poorly understood, even although evolutionary turnover of plant lineages at the KPB is central to understanding the assembly of the Cenozoic biota. One aspect that has received considerable attention is the apparent concentration of whole genome duplication (WGD) events around the KPB, which may have played a role in survival and subsequent diversification of plant lineages. In order to gain new insights into the origins of Cenozoic biodiversity, we examine the origin and early evolution of the legume family, one of the most important angiosperm clades that rose to prominence after the KPB and for which multiple WGD events are found to have occurred early in its evolution. The legume family (Leguminosae or Fabaceae), with c. 20.000 species, is the third largest family of Angiospermae, and is globally widespread and second only to the grasses (Poaceae) in economic importance. Accordingly, it has been intensively studied in botanical, systematic and agronomic research, but a robust phylogenetic framework and timescale for legume evolution based on large-scale genomic sequence data is lacking, and key questions about the origin and early evolution of the family remain unresolved. We extend previous phylogenetic knowledge to gain insights into the early evolution of the family, analysing an alignment of 72 protein-coding chloroplast genes and a large set of nuclear genomic sequence data, sampling thousands of genes. We use a concatenation approach with heterogeneous models of sequence evolution to minimize inference artefacts, and evaluate support and conflict among individual nuclear gene trees with internode certainty calculations, a multi-species coalescent method, and phylogenetic supernetwork reconstruction. Using a set of 20 fossil calibrations we estimate a revised timeline of legume evolution based on a selection of genes that are both informative and evolving in an approximately clock-like fashion. We find that the root of the family is particularly difficult to resolve, with strong conflict among gene trees suggesting incomplete lineage sorting and/or reticulation. Mapping of duplications in gene family trees suggest that a WGD event occurred along the stem of the family and is shared by all legumes, with additional nested WGDs subtending subfamilies Papilionoideae and Detarioideae. We propose that the difficulty of resolving the root of the family is caused by a combination of ancient polyploidy and an alternation of long and very short internodes, shaped respectively by extinction and rapid divergence. Our results show that the crown age of the legumes dates back to the Maastrichtian or Paleocene and suggests that it is most likely close to the KPB. We conclude that the origin and early evolution of the legumes followed a complex history, in which multiple nested polyploidy events coupled with rapid diversification are associated with the mass extinction event at the KPB, ultimately underpinning the evolutionary success of the Leguminosae in the Cenozoic.

---

The Cretaceous-Paleogene (K-Pg) boundary (KPB), 66 Million years ago (Ma), is defined by the mass extinction event that famously killed the non-avian dinosaurs and led to major turnover in the earth’s biota. The Chicxulub meteorite impact is generally thought to have been the cause of the mass extinction, but Deccan trap flood basalt volcanism likely contributed or may have been the primary cause, in line with previous global mass extinctions that are all related to volcanism (Keller, 2014). The KPB event determined in significant part the composition of the Earth’s modern biota, because many lineages that were successful in repopulating the planet and diversifying in the wake of the KPB have remained abundant and diverse throughout the Cenozoic until the present. Probably the best-known examples of successful post-KPB lineages are the mammals and birds, both inconspicuous elements of the Cretaceous fauna, while their core clades Placentalia and Neoaves became ubiquitous throughout Cenozoic fossil faunas. Plants were also severely affected by the KPB, with a clear shift in floristic composition and a drop in macrofossil species richness of up to 78% reported across boundary-spanning fossil sites in North-America (Wilf & Johnson, 2004; McElwain & Punyasena, 2007; Vajda & Bercovici, 2014). In addition, a global fungal spike followed by a global fern spike in the palynological record (Vajda et al., 2001; Barreda et al., 2012) are consistent with sudden ecosystem collapse and a recovery period characterized by low diversity vegetation dominated by ferns. Although the KPB is not considered a major extinction event for plants as no plant family appears to have been lost at the KPB (McElwain & Punyasena, 2007; Cascales-Miñana & Cleal, 2014), a sudden increase in net diversification rate in the Paleocene has been inferred from a large paleobotanical data set (Silvestro et al., 2015), suggesting increased origination following the KPB. Arguably, analyses of global plant fossil data suffer from the poor rock record in the Maastrichtian just prior to the KPB (Nicholls & Johnson, 2008) and are limited to inferences at family or genus level due to the nature of palaeontological data, thereby potentially underestimating global extinction rates at the species level.

For individual plant lineages, macro-evolutionary dynamics relative to the KPB extinction event have received less attention than prominent vertebrate clades. However, given that plants are the main primary producers and structural components of terrestrial ecosystems, the shaping and diversification of Cenozoic biota cannot be fully understood without understanding the consequences of the KPB for evolutionary turnover of plant diversity. The legume family (Leguminosae or Fabaceae), perhaps more than any other plant clade, appears to parallel Placentalia and Neoaves. No fossils are known that pre-date the KPB and are clearly identifiable to the legume family (Herendeen & Dilcher, 1992), but the family was already abundant and diverse in one of the earliest examples of modern type rainforests in the Paleocene (Wing et al., 2009; Herrera et al., submitted). The oldest known fossils that are already referable to (stem groups of) subfamilies are from close to the Paleocene-Eocene Thermal Maximum (PETM) (morphotype # CJ76 of c. 58 Ma (Wing et al., 2009) can be referred to Caesalpinioideae and *Barnebyanthus buchananensis* of c. 56 Ma to Papilionoideae (Crepet & Herendeen, 1992)) and legumes are a ubiquitous element of many Eocene, Oligocene and Neogene floras (Herendeen & Dilcher, 1992). Today, it is the third most species-rich angiosperm family, and arguably the most spectacular evolutionary and ecological radiation of any angiosperm family (McKey, 1994). Leguminosae is subdivided into six subfamilies (Fig. 1A-F; LPWG, 2017), which share the defining feature of the family, the fruit (referred to as the “legume” or “pod”) (Fig. 1G). It is the second most cultivated plant family after the Poaceae, and its species serve many purposes for humans, including timber, ornamentals, fodder crops and perhaps most notably, a large set of globally important pulse crops (Fig. 1I). A key trait of many legumes is the ability to fix atmospheric nitrogen via symbiosis with “rhizobia”-bacteria in root nodules (Fig. 1H), which leads to enriched soil, high nitrogen content in the leaves, and protein-rich seeds. The fact that legume species are diverse, omnipresent and often abundant in nearly all vegetation types across the planet, ranging in habit from large rainforest trees to small temperate herbs (Fig. 1J-L), means that legumes are an excellent study system to understand plant evolution in the Cenozoic.

**Figure 1.**
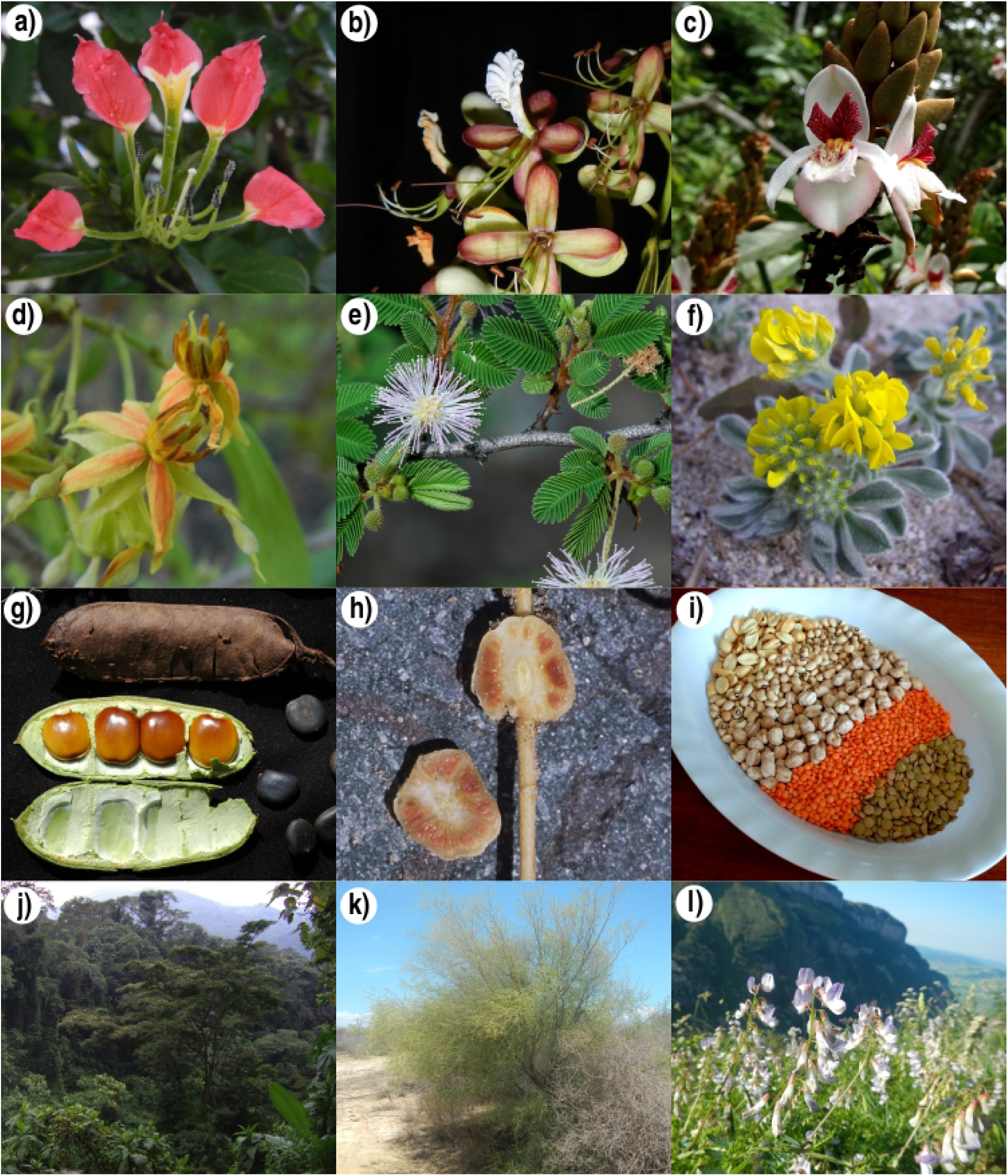
Diversity, ecology and economic importance of legumes. The family is subdivided into subfamilies (A) Cercidoideae (*Bauhinia madagascariensis*), (B) Detarioideae (*Macrolobium* sp.), (C) Duparquetioideae (*Duparquetia orchidacea*), (D) Dialioideae (*Baudouinia* sp.), (E) Caesalpinioideae (*Mimosa pectinatipinna*) and (F) Papilionoideae (*Medicago marina*). While the family has a very diverse floral morphology, the fruit (G), which comes in many shapes and is most often referred to as ‘pod’ or ‘legume’, is the defining feature of the family (fruit shown is of *Brodriguesia santosii*). A large fraction of legume species is known to fix atmospheric nitrogen symbiotically with ‘rhizobia’, bacteria that are incorporated in root nodules, for example in *Lupinus nubigenus* (H). Economically, the family is the second most important of flowering plants after the grasses, with a wide array of uses, including timber, ornamentals, fodder crops, and notably, pulse crops such as peanuts (Arachis), beans (Phaseolus), chickpeas (Cicer) and lentils (Lens) (I). Also ecologically, legumes are extremely diverse and important, occurring and often dominating globally across disparate ecosystems, including wet tropical forest, for example *Albizia grandibracteata* in the East African Albertine Rift (J), savannas, seasonally dry tropical forests, and semi-arid thorn-scrub, for example *Mimosa delicatula* in Madagascar (K) and temperate woodlands and grasslands, for example *Vicia sylvatica* in the European Alps (L). --Photos A, B, D, F, J, K, L by Erik Koenen, C by Jan Wieringa and E, G, H, I by Colin Hughes.

The rapid appearance of legume diversity shortly after the first occurrence in the fossil record has been likened to the ‘abominable mystery’ of the sudden appearance of the angiosperms (Sanderson, 2015). The legume phylogeny also suggests rapid early evolution of legume diversity with very short internodes subtending the six major lineages following the origin of the family (Lavin et al., 2005; LPWG, 2017) as well as at the base of subfamilies Detarioideae, Caesalpinioideae (Bruneau et al., 2008; LPWG, 2017) and Papilionoideae (Cardoso et al., 2012, 2013; LPWG, 2017). Just as for Placentalia (Teeling & Hedges, 2013) and Neoaves (Suh et al., 2015; Suh, 2016), this apparently rapid early diversification of legumes poses problems for phylogeny inference. In particular, the first few dichotomies in the phylogeny of the family have been difficult to resolve, as have deep divergences in Detarioideae, Caesalpinioideae and Papilionoideae (LPWG, 2013 & 2017). In this study, we attempt to resolve the deep-branching relationships in the legume family by using much larger molecular sequence data sets than those previously used in legume phylogenetics. Moreover, previous legume phylogenies have been mainly inferred from chloroplast markers (Wojciechowski et al., 2004; Lavin et al., 2005; Bruneau et al., 2008; Simon et al., 2009; Cardoso et al., 2012, 2013; LPWG, 2017). In addition to analysing nearly all protein-coding genes from the chloroplast genome, here we also analyse thousands of gene alignments from the nuclear genome.

Unlike birds and mammals, whole genome duplication (WGD) events are common in angiosperms, and such events have been suggested to be significantly concentrated around the KPB (Fawcett et al., 2009; Vanneste et al., 2014; Lohaus & Van de Peer, 2016). This is explained by the idea that polyploid lineages could have had enhanced survival and establishment across the KPB (Lohaus & Van de Peer, 2016) as well as greater potential to diversify rapidly thereafter relative to diploids (Levin & Soltis, 2018). WGDs have also been found to have occurred multiple times during the early evolution of the legumes (Cannon et al., 2015) and could have contributed to the initial rapid diversification of the family, as well to the difficulties of resolving relationships among the six subfamilies. There is considerable uncertainty about how many WGDs were involved in the early evolution of legumes and in the placements of possible WGDs on the legume phylogeny. From whole genome sequencing studies, it has been known for some time that several papilionoids share a WGD event (Cannon et al., 2006; Mudge et al., 2005), but recently it has been suggested that several other legume lineages have also undergone independent WGDs (Cannon et al., 2015). Indeed, Cannon et al. (2015) showed that the papilionoid WGD is shared by all members of that subfamily using phylogenetic methods, and used age estimates from *K*_*s*_ plots to infer additional independent WGDs early in the evolution of subfamilies Caesalpinioideae, Cercidoideae and Detarioideae. However, in the absence of data for several critical legume lineages, the phylogenetic positions of these additional putative WGDs remain uncertain. A more recent study (Wong et al., 2017) suggested instead that all legumes share the same WGD, based on rate-corrected *K*_*s*_ plots and a genetic linkage map of *Acacia* that suggested mimosoids (Caesalpinioideae) and Papilionoideae retained an orthologous duplicated chromosomal segment. From homolog gene family trees generated prior to separating orthologs from paralogs (Yang & Smith, 2014; Smith et al., 2015; Yang et al., 2015), we map the number of gene duplications over the legume phylogeny to evaluate how many early legume WGDs occurred and where they are located on the phylogeny.

While the legumes are not known with certainty from any Cretaceous fossil site, the family has a long stem lineage dating back to c. 80 – 100 Ma (Wang et al., 2009; Magallón et al., 2015). This long ghost lineage means that the timing of the initial radiation of the family, as well as of legume WGDs, and notably whether they pre- or post-date the KPB, are uncertain. In Placentalia and Neoaves, divergence time estimation has led to much debate, with some studies using molecular sequence data for divergence time estimation suggesting that both clades originated and diversified well before the KPB, implying that many lineages of both clades survived the end-Cretaceous event (Cooper & Penny, 1997; Jetz et al., 2012; Meredith et al., 2011). However, like the legumes, both groups first appear in the Paleocene fossil record. A phylogenetic study of mammals combining both molecular sequence data and morphological characters to enable inclusion of fossil taxa, found only a single placental ancestor crossing the KPB (O’Leary et al., 2013; but see Springer et al., 2013; dos Reis et al., 2014). Alternatively, it has been argued that diversification of Placentalia followed a “soft explosive” model, with a few lineages crossing the KPB followed by rapid ordinal level radiation during the Paleocene (Phillips, 2015; Phillips & Fruciano, 2018). Recent time-calibrated phylogenies for birds showed the age of Neoaves to also be close to the KPB (Jarvis et al., 2014; Claramunt & Cracraft, 2015; Prum et al., 2015), with initial rapid post-KPB divergence represented by a hard polytomy (Suh, 2016). For legumes, it is similarly unlikely that modern subfamilies of legumes have Cretaceous crown ages. These clades, in particular Papilionoideae, Caesalpinioideae and Detarioideae, appear to have rapidly diversified following their origins, which would imply mass survival of very large numbers of legume lineages across the KPB. Diversification into the six main lineages of legumes appears to have occurred rapidly (Lavin et al., 2005), with long stem branches leading to each of the modern subfamilies. Therefore, two hypotheses seem plausible: (1) the legumes have a Cretaceous crown age and diversified into the six subfamilies prior to the KPB, while crown radiations of the subfamilies occurred in the wake of the mass extinction event, corresponding to a “soft explosive” model, or (2) a single legume ancestor crossed the KPB and rapidly diversified into six main lineages in the wake of the mass extinction event, corresponding to a “hard-explosive” model, with the subsequent subfamily radiations related to the Paleocene-Eocene Thermal Maximum (PETM) and/or Eocene climatic optimum. Currently available molecular crown age estimates for the family range from c. 59 to 64 Ma (Lavin et al., 2005; Bruneau et al., 2008; Simon et al., 2009). These studies, however, lacked extensive sampling of outgroup taxa and relied instead on fixing the stem age of the legumes, thereby compromising the ability to estimate the crown age of the family. Furthermore, these earlier studies relied exclusively on chloroplast sequences, for which evolutionary rates are known to vary strongly across legumes (Lavin et al., 2005), such that nuclear gene data are likely to be better suited for estimating divergence times (Christin et al., 2014).

In this study, using large genomic-scale data sets, we aim to resolve the deep divergences in the legume family, find the phylogenetic locations of WGDs and estimate the timing of these. We analyse these new datasets with Maximum Likelihood (ML) analysis, Bayesian inference, a multi-species coalescent summary method and filtered supernetwork reconstruction to resolve the deep-branching relationships in the family. In particular, we focus on the relationships among the six major lineages recently recognized as subfamilies (LPWG, 2017). Sister-group relationships between subfamilies Papilionoideae and Caesalpinioideae (sensu LPWG, 2017), and of the clade combining the two with the newly recognized Dialioideae, were previously known (Lavin et al., 2005; Bruneau et al., 2008; LPWG, 2017). However, the relationships between the clade comprising those three subfamilies and the other three subfamilies Cercidoideae, Detarioideae and Duparquetioideae remained difficult to resolve (cf. Bruneau et al., 2008; LPWG, 2017). Having inferred the most likely species-tree topology, we evaluate numbers of supporting and conflicting bipartitions for critical nodes across gene trees. To infer likely locations of WGDs, we count the number of gene duplications present in nuclear homolog clusters and map these across the species tree. Finally, we perform molecular clock dating on a selection of informative and clock-like nuclear genes with 20 fossil calibration points, to infer whether the origin of the legumes and WGDs in the early evolution of the family are related to the K-Pg mass extinction event.

## Material & Methods

### DNA/RNA Extraction and Sequencing

For the newly generated chloroplast gene data, DNA was extracted from fresh leaves, leaf tissue preserved in silica-gel or herbarium specimens, using the Qiagen DNeasy Plant Mini Kit. Sequencing libraries were prepared using the NEBNext Ultra DNA Library Prep Kit for Illumina. They were then sequenced on the Illumina HiSeq 2000 sequencing platform, at low coverage (‘genome-skimming’) or as part of hybrid capture experiments for a separate study on mimosoid legumes (Koenen et al., unpublished data). RNA was extracted from fresh leaves using the Qiagen RNeasy Plant Mini Kit. RNA sequencing libraries were prepared using the Illumina TruSeq RNA Library Prep Kit and sequenced on the Illumina HiSeq 2000 sequencing platform. All lab procedures were performed according to the specifications and protocols provided by the manufacturers of the kits.

### Sequence Assembly

Raw reads for the chloroplast DNA data were cleaned and filtered using the following steps: (1) Illumina adapter sequence artifacts were trimmed using Trimmomatic v. 0.32 (Bolger et al., 2014), (2) overlapping read pairs were merged with PEAR v. 0.9.8 (Zhang et al., 2014) and (3) low quality reads were discarded and low quality bases at the end of the reads were trimmed with Trimmomatic v. 0.32 (using settings MAXINFO:40:0.1 LEADING:20 TRAILING:20). The quality-filtered reads were then assembled into contigs using the SPAdes assembler v. 3.6.2 (Bankevich et al., 2012). For RNA data, raw reads were quality-filtered using the FASTX-toolkit v. 0.0.13 (http://hannonlab.cshl.edu/fastx_toolkit/index.html) to remove low quality reads (less than 80% of bases with a quality score of 20 or higher), TagDust v. 1.12 (Lassmann et al., 2009) to remove adapter sequences and PRINSEQ-lite v. 0.20.4 (Schmieder & Edwards, 2011) to trim low quality bases off the ends of reads. Transcriptome assembly was then performed on the quality-filtered reads using Trinity (Grabherr et al., 2011; Release 2012-06-08), with default settings.

### Chloroplast Proteome Alignment

DNA sequences of protein-coding chloroplast genes were newly generated as described above, or extracted from several different data sources, as specified for each accession in Table S1. Sequence data were extracted directly from annotated plastomes in Genbank, by blast searches from *de novo* assembled contigs and from transcriptomes using custom Python scripts. Sequences for some outgroup taxa (data from Moore et al., 2010) were downloaded separately per gene from Genbank. For each gene, a codon alignment was inferred using MACSE v. 1.01b (Ranwez et al., 2011). Phylogenetic trees were then inferred for each gene separately to screen for erroneously aligned sequences with RAxML v. 8.2 (Stamatakis, 2014). For some species, individual gene sequences that led to anomalously long terminal branches were then removed. The genes *accD* and *clpP* were removed completely. The gene alignments were concatenated and the full alignment was visually checked and obvious misalignments were resolved. Furthermore, sequence errors (single A/T indels) that caused frameshift mutations were corrected and the accuracy of the alignment at codon level was assessed and corrected if necessary. For a few genes where the ends of coding sequences had varying lengths, all sites between the first and last stop codon in the alignment were excluded, since they were poorly aligned. Finally, using BMGE v. 1.12 (Block Mapping and Gathering with Entropy; Criscuolo & Gribaldo, 2010) the codon alignment was translated to amino acid sequences.

### Nuclear Gene Data and Matrix Assembly

Whole genome and transcriptome data were downloaded from various sources and augmented with newly generated transcriptome sequence data for six Caesalpinioideae and Detarioideae taxa (see Table S2). Peptide sequences were downloaded from annotated genomes, or were extracted from transcriptome assemblies using TransDecoder (http://transdecoder.github.io/). To assemble the nuclear peptide sequence data into aligned gene matrices, we used the pipeline of Yang & Smith (2014). We performed mcl clustering as described in Yang & Smith (2014), with a hit fraction cut-off of 0.75, inflation value of 2 and a minimum log-transformed e-value of 30. These settings lead to clusters with good overlap between sequences and good alignability (omitting genes that are too variable), although we may have lost a few short gene clusters. Next, the homolog gene clusters were subjected to two rounds of alignment with MAFFT v. 7.187 (Katoh & Standley, 2013), gene tree inference inference with RaxML v. 8.2 (Stamatakis, 2014), and pruning and masking of tips and cutting deep paralogs as described in Yang & Smith (2014). In the first round we used 0.3 and 1.0 as relative and absolute cut-offs for trimming tips, respectively, and 0.5 as the minimum cut-off for cutting deep paralogs, and keeping all clusters with a minimum of 25 taxa for the second round. In the second round we used more stringent cut-off values (0.2 and 0.5 for trimming tips and 0.4 for cutting deep paralogs). See Yang & Smith (2014) for more information on these parameter settings. One-to-one orthologs and rooted ingroup (RT) homologs were then extracted from the homolog cluster trees, with a minimum aligned length of 100 amino acids for each homolog. One-to-one orthologs are those homolog gene clusters in which each taxon is represented only by a single gene copy. RT homologs are extracted by orienting homolog cluster trees by rooting them on the outgroup (in our case *Aquilegia coerulea* and *Papaver somniferum*), and then detecting gene duplications and pruning the paralog copies with fewer taxa present until each taxon is represented by a single copy. The outgroup is pruned as well, and clusters without outgroup in which each taxon is only present once are also included, meaning that all 1-to-1 orthologs are also in the RT homolog set. See Yang & Smith (2014) for a more detailed description of how these homologs are extracted. Sequences with more than 50% gaps and all sites with more than 5% missing data were removed from the homolog alignments using BMGE. For the 1-to-1 orthologs that were used for species tree inference, alignments with fewer than 50 taxa were discarded, for the larger set of RT homologs that were used for counting of supporting and conflicting bipartitions, alignments with fewer than 25 taxa were discarded.

### Phylogenetic Inferences

Maximum likelihood (ML) and Bayesian analyses were run in RaxML v. 8.2 (Stamatakis, 2014) and Phylobayes-MPI 1.7 (Lartillot et al., 2013), respectively. For the ML analysis using nucleotide sequences of the chloroplast alignment, we used PartitionFinder 2 (Lanfear et al., 2017) to estimate partitions, with a minimum length per partition set to 500 nucleotides, and allowing different codon positions per gene to be in different partitions. The resulting 16 partitions were run with the GTR + GAMMA model, and 1000 rapid bootstrap replicates were carried out. For the amino acid sequences, the ML analyses of both the chloroplast alignment and the concatenated alignment of nuclear 1-to-1 orthologs were analyzed with the LG4X model, without partitioning, as the model accounts for substitution rate heterogeneity across the alignment by estimating 4 different LG substitution matrices (Le et al., 2012). For the chloroplast alignment, 1000 rapid bootstrap replicates were additionally carried out. Gene trees of 1-to-1 orthologs and RT homologs were estimated with RAxML using the WAG + G model, with 100 rapid bootstrap replicates. We then calculated 80% majority-rule consensus trees for each ortholog or homolog and used these to calculate Internode Certainty All (ICA) values using RAxML, to include only nodes that received 80% or greater bootstrap support in the individual gene trees. Bayesian analyses were performed with the CATGTR model, with invariant sites deleted and default settings for other options in Phylobayes. Analyses were run until the chain reached convergence (usually after 10-20k cycles), with at least two independent chains run for each data set. To perform Bayesian analyses on the complete nuclear gene data set in a computationally tractable manner, we ran 25 gene jack-knifing replicates without replacement, dividing the total number of genes over 5 subsets with 5 replicates. These subsampled replicates were run in Phylobayes-MPI, with a starting tree derived from the analysis sampling the 100 genes with the longest gene tree length, using the CATGTR model with constant sites deleted, for 1000 cycles each. We found that all 25 chains had converged after a few hundred cycles, and discarded the first 500 cycles of each as burn-in. A majority-rule consensus tree was constructed using sumtrees.py (from the Dendropy library (Sukumaran et al., 2010)) from 12500 total posterior trees, representing the MCMC cycles 501-1000 of each replicate. For both the ML and Bayesian analyses, concatenated alignments were not partitioned. Instead we rely on the LG4X and CATGTR models to take rate heterogeneity into account, since these models describe heterogeneity across alignments more accurately than partitioning by gene and/or codon since the substitution process also varies across gene sequences and codon positions. For the multi-species coalescent analysis, we used ASTRAL (Mirabab et al., 2014) on the 1,103 gene trees estimated with RAxML, using local posterior probability and quartet support to evaluate the inferred topology (Sayyari & Mirabab, 2016).

### D_n_/D_s_ Ratio Analyses for cpDNA

The codon alignments for each chloroplast gene were analyzed individually using the branch model test in PAML v. 4.9 (Yang, 2007), to test if higher substitution rates in the 50-kb inversion and vicioid clades of Papilionoideae were related to differing selective pressures. These clades were partitioned separately to allow for the estimation of independent rates of synonymous and non-synonymous substitution rates for each of these clades relative to the rest of the tree. Since the vicioid clade is nested in the 50-kb inversion clade, the rates reported for the latter clade are estimated without the vicioid clade taxa. While this test does not evaluate selective pressures for specific sites, it does give an indication whether genes evolve neutrally or are under purifying or positive selection.

### Counting Supporting Bipartitions for Key Nodes across Gene Trees

Using a custom python script, numbers of matching and alternative bipartitions across gene trees were counted for particular nodes labeled A-H in Figure 3A in the legume phylogeny. For this purpose, we assessed monophyly of each of the subfamilies and combinations (clades) of subfamilies, against the outgroup, across all gene trees. For each gene tree, we first assessed whether all 6 groups (5 subfamilies plus the outgroup) are present and gene trees with missing groups were not taken into account. Next, we evaluated whether the gene tree includes a matching bipartition for the family, each subfamily and for all possible combinations of subfamilies. A matching bipartition means that all taxa of a subfamily or combination of subfamilies are separated from all other taxa in the gene tree, thus constituting support for that clade to be monophyletic. For combinations of subfamilies, the subfamilies themselves do not necessarily need to be monophyletic, but all taxa within those subfamilies should be separated from all other taxa to constitute a matching bipartition, and thus to be a supported clade in the gene tree. For well supported clades, we expect matching bipartitions for a majority of gene trees. For poorly supported clades, we expect most gene trees to be uninformative due to low phylogenetic signal, hence a low number of matching bipartitions, and possibly relatively high numbers of conflicting bipartitions. All counts were done for ML gene trees of RT homologs, and with 50 and 80% bootstrap cutoffs. The recently published DiscoVista software package (Sayyari et al., 2018) allows similar evaluations of conflicting and supporting bipartitions to those described here to be made and visualized.

### Phylogenetic Supernetwork Analysis

We used SplitsTree4 to draw a filtered supernetwork (Whitfield et al., 2008) of the 1,103 1-to-1 orthologs, using the 80% majority-rule consensus trees to only include well-supported bipartitions to infer the network. All gene trees were pruned for simplified visualization, focusing on the deep divergences within the legume family. All taxa outside the nitrogen-fixing clade comprising Cucurbitales, Fabales, Fagales, Rosales, as well as a subset of taxa in the relatively densely sampled Papilionoideae and Caesalpinioideae were pruned, preferentially keeping taxa that were sampled in as many gene trees as possible. The mintrees parameter was set to 552 (at least 50% of the number of orthologs) and the maximum distortion parameter was set to 0.

### Gene Duplication Mapping

We used the homolog clusters generated from the Yang & Smith (2014) pipeline prior to extracting 1-to-1 and RT orthologs to map duplications onto the species tree. First, all sites with more than 5% missing data were removed with BMGE, to reduce the amount of missing data. Also all sequences with more than 75% gaps were removed, to avoid having fragmented paralog sequences present, which could inflate the number of gene duplications. These data removal steps also led to the elimination of some clusters with large amounts of missing data. Tree estimation was then repeated on these clusters, with RAxML using the WAG + G model and 100 rapid bootstrap replicates. Next, rooted ingroup clades were extracted from the resulting homolog trees with the extract_clades.py script that is included with the Yang & Smith (2014) pipeline. To extract the clades, we only considered *Aquilegia* and *Papaver* as outgroup taxa, because the outgroup is not included in the extracted clades, and this way we could maximize the number of taxa per extracted clade. However, because of uncertain relationships along the backbone of Pentapetalae, we observed that the clusters were often not correctly rooted. This does not have much effect for the number of duplications that are observed near the tips, but it does lead to erroneous mapping near the base of the tree. Therefore, we rooted the extracted clades with the Phyx package (Brown et al., 2017), using a list of the non-legume taxa ordered by their phylogenetic relationships, rooting the trees on the taxon that is most distantly related to legumes. Clusters that included only legume species, without any outgroup taxa present, were excluded. From the resulting multi-labeled trees (i.e. each taxon can be present multiple times, representing different paralogs), duplications were mapped onto the species tree, with and without a 50% bootstrap cut-off, using phyparts (Smith et al., 2015).

### Divergence time analyses

Fossils used to calibrate molecular clock analyses are listed in Table 1 and are discussed in Methods S1.

**Table 1.**
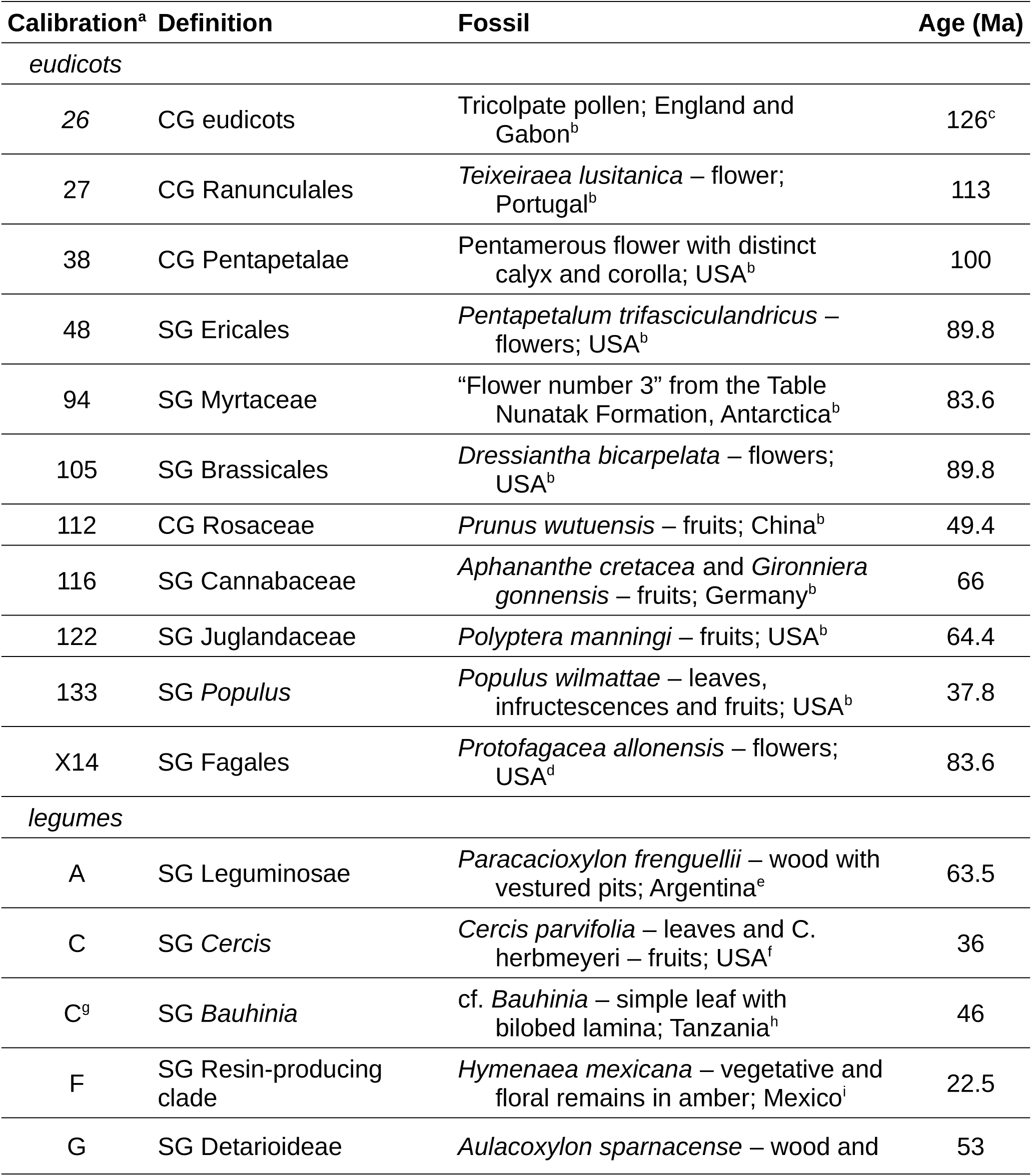
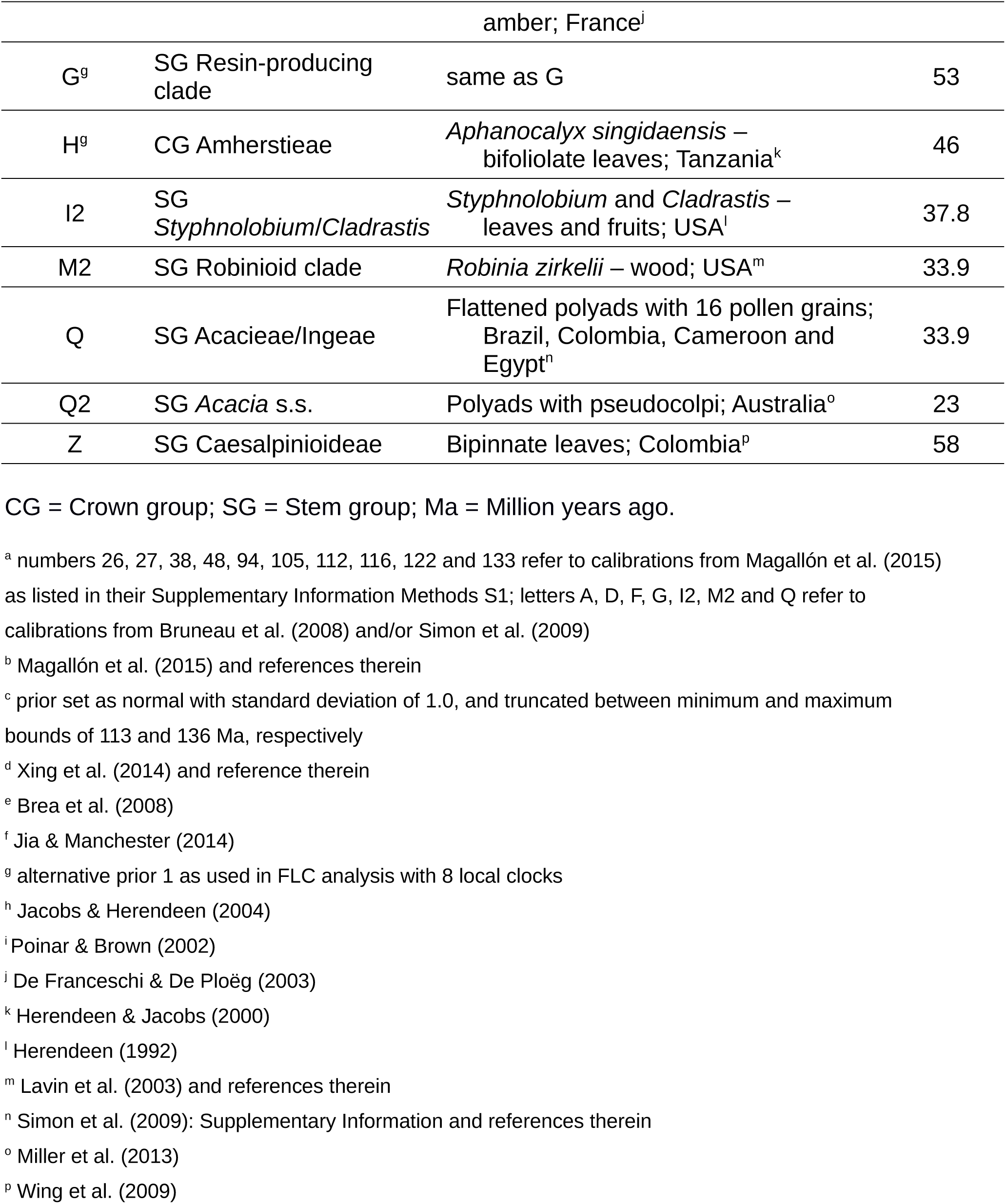
Fossil calibrations used in the divergence time analyses.

Using SortaDate (Smith et al., 2018b), we analyzed all gene trees to estimate the total tree length (a proxy for sequence variation or informativeness), root-to-tip variance (a proxy for clock-likeness) and compatibility of bipartitions with the ML tree that was inferred using the full data set (the RAxML tree inferred with the LG4X model, shown in Figure 3A). We then selected the best genes for dating based on cutoff values that were arbitrarily chosen from the estimated values across gene trees: (1) total tree length greater than 5, (2) root-to-tip variance less than 0.005 and (3) at least 10% of the bipartitions in common with the ML tree. This yielded 36 genes, which were concatenated to have a total aligned length of 14462 amino acid sites. We also used the ‘pxlstr’ program of the Phyx package (Brown et al., 2017) to calculate taxon-specific root-to-tip lengths from the ML tree, after pruning the Ranunculales, on which the tree was rooted. The values obtained were then used to define local clocks as described below. *Arabidopsis thaliana, Linum usitatissimum* and *Polygala lutea* were removed because of much higher root-to-tip lengths relative to their closest relatives. *Panax ginseng* was also removed because of a low root-to-tip length relative to the other sampled asterids, leaving a total of 72 taxa.

We used BEAST v.1.8.4 (Drummond et al., 2012) with various clock models to estimate divergence time estimates across the phylogeny based on the alignment of the selected 36 genes and the fossil calibrations described above. All analyses were run with the LG + G model of amino acid substitution and the birth-death tree prior, and using the ML tree to fix the topology. Fossil calibration priors were set as uniform priors between the minimum age as specified in Table 1 and a maximum age of 126 Ma (oldest fossil evidence of eudicots) as listed in Table S4, with the exception of the root node, for which we used a normal prior at 126 Ma with a standard deviation of 1.0 and truncated to minimum and maximum ages of 113 (the Aptian-Albian boundary) and 136 Ma (the oldest crown angiosperm fossil, see Magallón et al. (2015)). With these settings, we ran analyses under the uncorrelated lognormal (UCLN), strict (STRC), random (RLC) and 3 different fixed local (FLC) clock models. To specify the different FLC models, we looked at root-to-tip length variation across subclades to specify biologically meaningful *a priori* clock partitions (Fig. S19). The 50kb-inversion clade of papilionoid legumes and the asterids (without *Panax ginseng*) have uniformly longer root-to-tip lengths than the other taxa across the tree and were therefore assigned their own local clock, with a different clock for the remaining taxa in the tree (this model referred to as FLC3, partitioning of taxa is illustrated in Supplementary Figure S19A). A more complex model was specified where the rosid rate was decoupled from the background rate and more clock partitions within the legumes were created for the mimosoids together with the *Cassia* clade because of their longer root-to-tip lengths relative to other Caesalpinioideae and most of the rosid clade and for the combined clade of Cercidoideae and Detarioideae as well. This more complex model is referred to as FLC6 (Fig. S19B). The most complex model (FLC8; Fig. S19C) was generated by further partitioning the combined clade of Cercidoideae and Detarioideae with a separate local clock for each subfamily, and one on their combined stem lineages (this most complex partitioning is also indicated with colored branches in Figures 6 and S16-17 and those of the other FLC models in Figures S14-15). The Ranunculales that were pruned for the root-to-tip length calculations were included in the background clock for each FLC model.

The separate clock partitions assigned to Cercidoideae and Detarioideae in the FLC8 model are particularly useful for evaluating the controversial placement of Early and Middle Eocene fossils within their crown groups (see Methods S1). This was done by running two analyses under the FLC8 model, one with the same priors as the other analyses, and one where calibrations C and G were changed and another calibration (H^g^) was added to use similar placements of these calibrations as in Bruneau et al. (2008) and Simon et al. (2009) (Table 1 & Methods S1). We refer to this calibration scheme as “alternative prior 1” (Table S4). Since a separate local clock is assigned to the combined stem lineages of Cercidoideae and Detarioideae, substitution rate estimates for stem and crown groups can be compared under both calibration schemes.

Maximum ages of fossil calibrations were set conservatively, and perhaps overly so, which can lead to a poorly formed joint marginal prior on node ages across the tree (Phillips, 2015). Therefore, we also constructed an alternative prior with less conservative maxima as specified in Table S4 (“alternative prior 2”). These maxima represent boundary ages of older epochs from which the crown or stem group is not known, and in line with ages found by Magallón et al. (2015). These analyses serve to test the sensitivity of the UCLN model to the marginal prior.

Analyses sampling from the prior (without data) were run for 100 million generations, the strict clock and FLC3 and FLC6 analyses were run for 25 million generations and all other clock analyses were run for 50 million generations, and convergence was confirmed with Tracer v1.7.1 (Rambaut et al., 2018). For the non-prior analyses, the first 10% of the total number of generations was discarded as burn-in before summarizing median branch lengths and substitution rates with TreeAnnotator from the BEAST package.

## Results

The chloroplast alignment includes 72 protein-coding genes, for 157 taxa (including 111 legume species; Table S1), with a total aligned length of 75,282 bp or 25,094 amino acid residues. From transcriptomes and fully sequenced genomes, we gathered 9,282 homologous nuclear encoded gene clusters for 76 taxa including 42 legume species (Table S2). From these clusters, we extracted protein alignments of 1,103 1-to-1 orthologs for species tree inference with a total aligned length of 325,134 amino acids when concatenated, and 7,621 Rooted Ingroup (RT) homologs for additional gene tree inference. We also extracted 8,038 rooted clades from the homolog clusters to map the locations of gene duplications. The alignments, gene trees and species trees are available in TreeBASE (accession number XXXX) and on Dryad (doi: XXXX).

### Inferring the Species Tree

Our analyses reveal that both the chloroplast and nuclear data sets resolve all subfamilies as monophyletic with full support and most relationships among the subfamilies are also robustly resolved (Figs 2, 3A-C & S1-7), with the notable exception of the root node. The clade consisting of Papilionoideae, Caesalpinioideae and Dialioideae is recovered in all analyses, with *Duparquetia* as the sister-group to this clade as inferred from chloroplast data. *Duparquetia* is not sampled for nuclear data, therefore transcriptome or genome sequencing is necessary for this taxon to confirm the relationship found here. The root node of the legume family is more difficult to resolve, and the chloroplast and nuclear data sets lead to conflicting topologies. The chloroplast alignment supports Cercidoideae as sister to the rest of the family when analysing protein sequences with ML under the LG4X model (58% bootstrap support; Fig. S1) and Bayesian inference under the CATGTR model (0.98 posterior probability (pp); Figs 2 & S2). When analysing chloroplast nucleotide sequences, we recovered the same relationship in a partitioned ML analysis under the GTR model (recovered in 66% of the bootstrap replicates; Fig. S3), but a Bayesian analysis under the CATGTR model does not resolve the root, the majority-rule consensus tree showing a polytomy of Cercidoideae, Detarioideae and a robustly supported clade formed by the other four subfamilies (Fig. S4). To resolve deep divergences, amino acid sequences are more suitable because they are less saturated with substitutions (silent substitutions are absent), and less prone to long branch attraction (LBA). Additionally, the LG4X and CAT models better account for heterogeneous substitution rates across sites in the alignment (Lartillot & Philippe, 2004; Le et al., 2012). Taken together, this suggests that the sister-group relationship of Cercidoideae with the rest of the family is the most likely rooting as inferred from chloroplast data, but given the low bootstrap support values, phylogenetic signal with regards to the root node appears to be limited. A notable observation is that the chloroplast genome evolves markedly faster in the 50kb-inversion clade of Papilionoideae than in other legumes (with even higher rates apparent in the vicioid clade), as is evident from both the nucleotide and amino acid alignments (Figs 2C & S1-4), suggesting that this pattern is not driven solely by synonymous substitutions. However, branch model *D*_*n*_/*D*_*s*_ ratio tests do not find any evidence of differential selection acting on chloroplast genes across the different clades (Fig. 2D), and suggest that the majority of chloroplast genes across legumes are under purifying selection.

**Figure 2.**
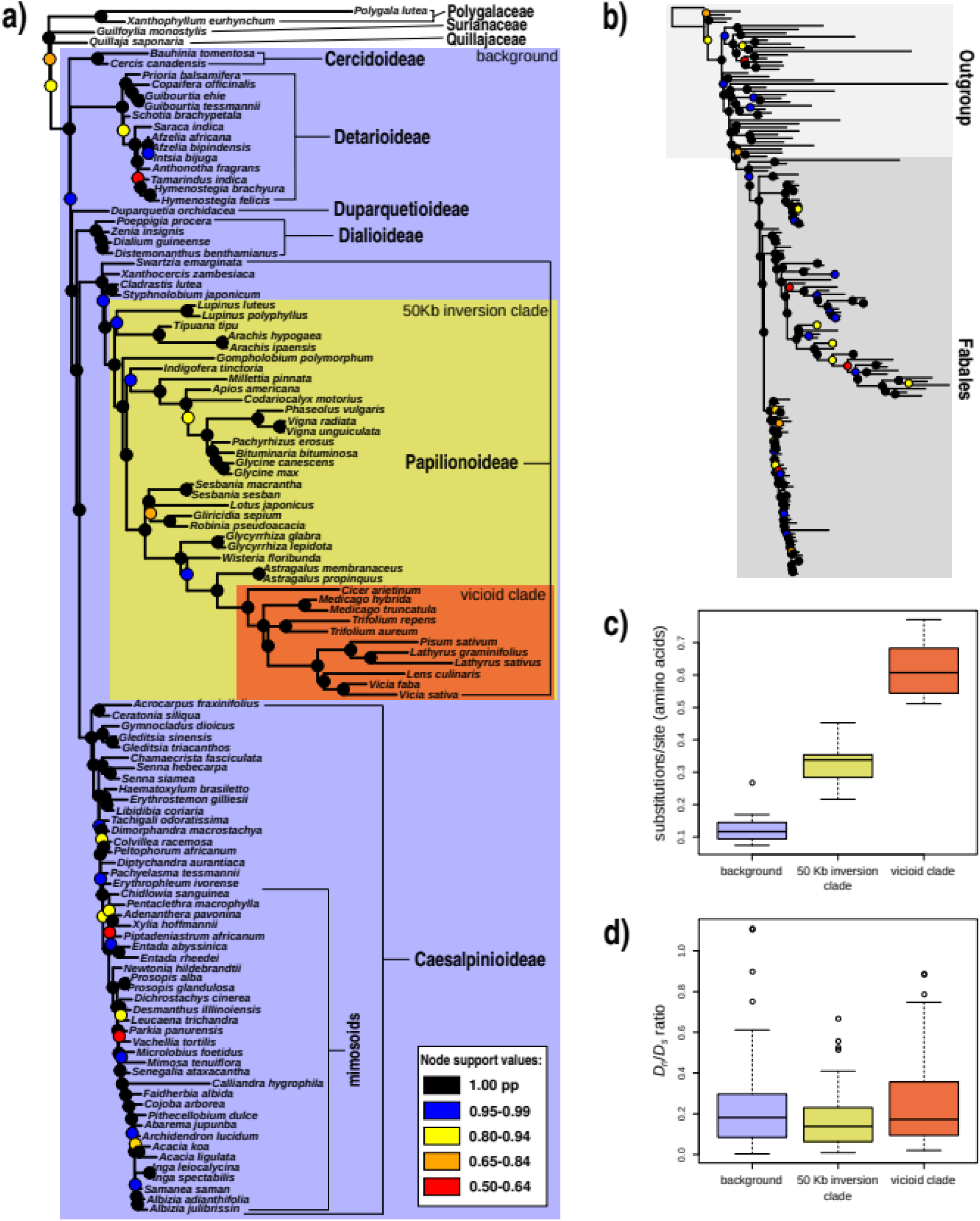
Phylogeny of legumes based on Bayesian analyses of 72 protein coding chloroplast genes under the CATGTR model in Phylobayes. (A) majority-rule consensus tree of the amino acid alignment, showing only the Fabales portion of the tree, outgroup taxa pruned, (B) complete tree including outgroup taxa, (C) Root-to-tip lengths measured from the legume crown node in amino acid substitutions per site and (D) *D_n_/ D_s_* ratios for background, the 50 Kb inversion clade (excluding the vicioid clade) and vicioid clade tree partitions. Majority-rule consensus trees for both the amino acid and nucleotide alignments with tip labels for all taxa and support values indicated are included in supporting information (Figs S1-2).

**Figure 3.**
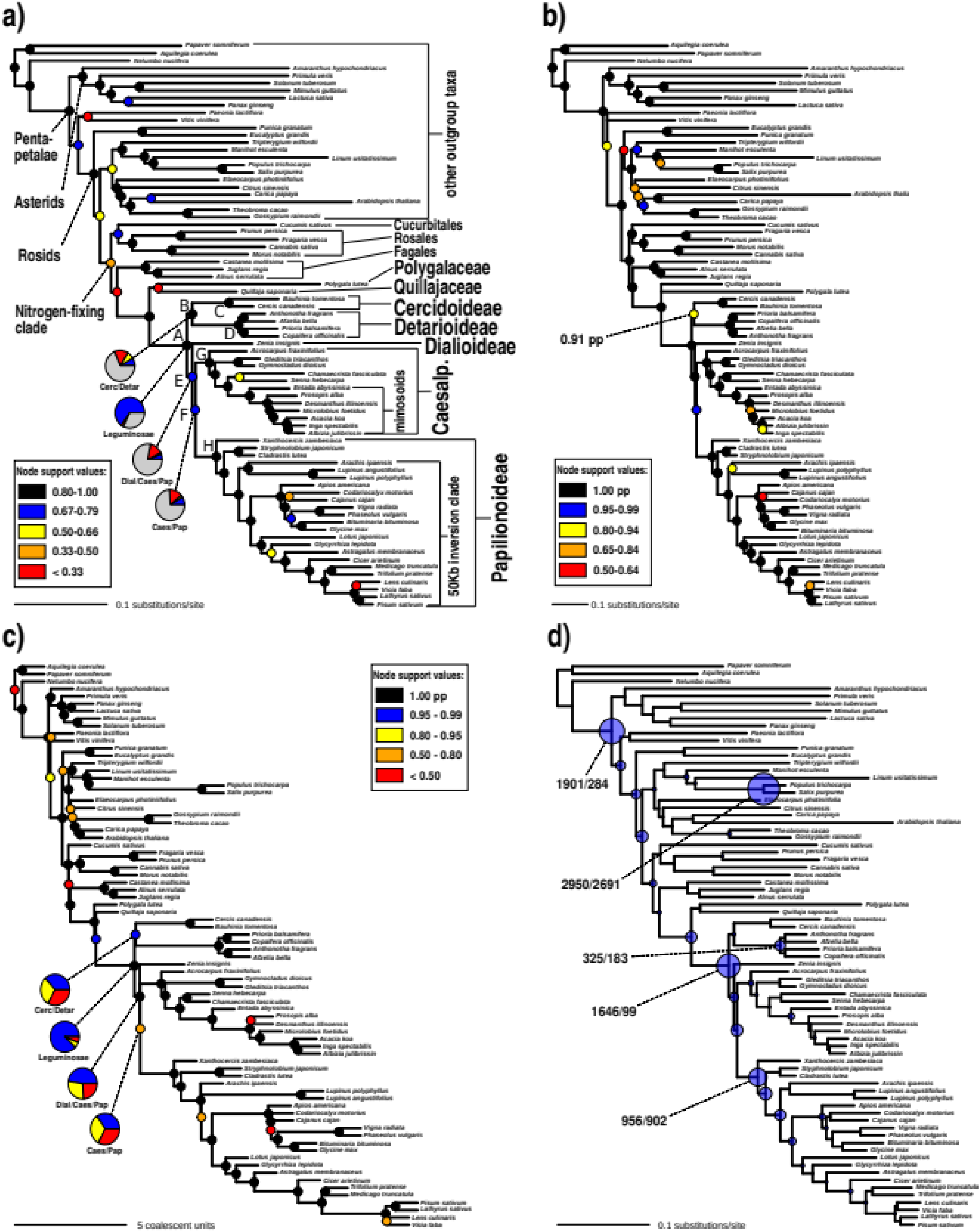
Congruent relationships among subfamilies when using different types of phylogenetic analysis, and phylogenetic locations of WGDs, as inferred from nuclear gene data. Support is indicated with coloured symbols on nodes for simplicity of presentation, as indicated in the legends; figures annotated with actual support values are included as Figures S5-7. (A) ML phylogeny estimated with RAxML under the LG4X model from a concatenated alignment of 1,103 nuclear orthologs. Support indicated represents Internode Certainty All (ICA) values, estimated with RAxML from 80% bootstrap threshold consensus gene trees of the same 1,103 orthologs. For the first four divergences in the legume family, pie charts indicate the proportions of gene trees supporting the relationship shown (blue), supporting the most prevalent conflicting bipartition (yellow), supporting other conflicting bipartitions (red) and genes without phylogenetic signal, i.e. no bootstrap support (gray). Numbers of bipartitions for the pie charts are derived from phyparts analyses with a 50% bootstrap support filter. Labelled nodes A-H are analysed in more detail in Figure 5. (B) Bayesian gene jackknifing majority-rule consensus tree of concatenated alignments of c. 220 genes per replicate, support indicated represents posterior probability averaged over 25 replicates for 500 posterior trees each (in total 12,500 posterior trees). (C) Phylogeny estimated under the multi-species coalescent with ASTRAL from gene trees, support indicated represents local posterior probability. Pie charts show relative quartet support for the first (blue) and the two (yellow and red) alternative quartets. (D) Gene duplications in 8,038 homolog clusters mapped onto the ML species tree topology. The size of the circles on nodes is proportional to the number of gene duplications inferred. For hypothesized WGD events, the number of gene duplications without/with bootstrap filter is indicated. See Figure S9 for the number of gene duplications for all nodes in the phylogeny.

In contrast to the results obtained with chloroplast data, in all analyses of the 1,103 nuclear 1-to-1 orthologs, we recover a sister-group relationship between Cercidoideae and Detarioideae, with this clade sister to the clade comprising of Dialioideae, Caesalpinioideae and Papilionoideae (note that Duparquetioideae is not sampled) (Figs 3A-C & S5-7). We inferred an ML tree of the concatenated alignment with the LG4X model, and calculated Internode Certainty All (ICA) values from bootstrapped gene trees on this topology (Fig. 3A & S5). Only bipartitions that received >80% bootstrap support were considered. The internode certainty metric was introduced to assess phylogenetic conflict among loci and identify internodes with high certainty, to be used in particular in phylogenomic studies where bootstrap values are often inflated (Salichos & Rokas, 2013). The sister-group relationship between Cercidoideae and Detarioideae is well-supported, receiving an ICA value of 0.85. A Bayesian jackknifing analysis with the CATGTR model infers a nearly identical topology to the ML topology (Fig. 3B & S6), with posterior probability of 0.91 in support of this same relationship. The multi-species coalescent species-tree inferred with ASTRAL (Mirabab et al., 2014), which accounts for incomplete lineage sorting (ILS), is also consistent with that relationship (Fig. 3C & S7), with the Cercidoideae/Detarioideae clade supported by a local posterior probability of 0.95 (Sayyari & Mirabab, 2016). In summary, all analyses of nuclear protein alignments lend strong support for a sister-group relationship between Cercidoideae and Detarioideae.

### Evaluation of Gene Tree Support and Conflict

While the chloroplast and nuclear phylogenies show a different topology with regards to the first two dichotomies within the legumes, the different types of analyses performed on the nuclear data set all yield the same topology at the base of the family (Figs 3A-C). Because the nuclear data set consists of 1,103 unlinked loci sampled from across the nuclear genome compared to the single locus that the chloroplast genome constitutes, this topology should be considered to be more likely. However, when evaluating gene tree conflict, it appears that a large number of conflicting bipartitions exist, with the most prevalent being nearly as frequent across gene trees as compatible bipartitions (pie charts in Figure 3A). The quartet support as calculated by ASTRAL is also low (37%, with alternative quartet supports 33% and 30%; pie charts in Figure 3C). The relationships among the remaining three sampled subfamilies are also supported by significantly fewer bipartitions and lower quartet support than for example the legume crown node (pie charts in Figures 3A & C). Furthermore, the filtered supernetwork shows a complex tangle of gene tree relationships at the base of the legumes (Fig. 4).

**Figure 4.**
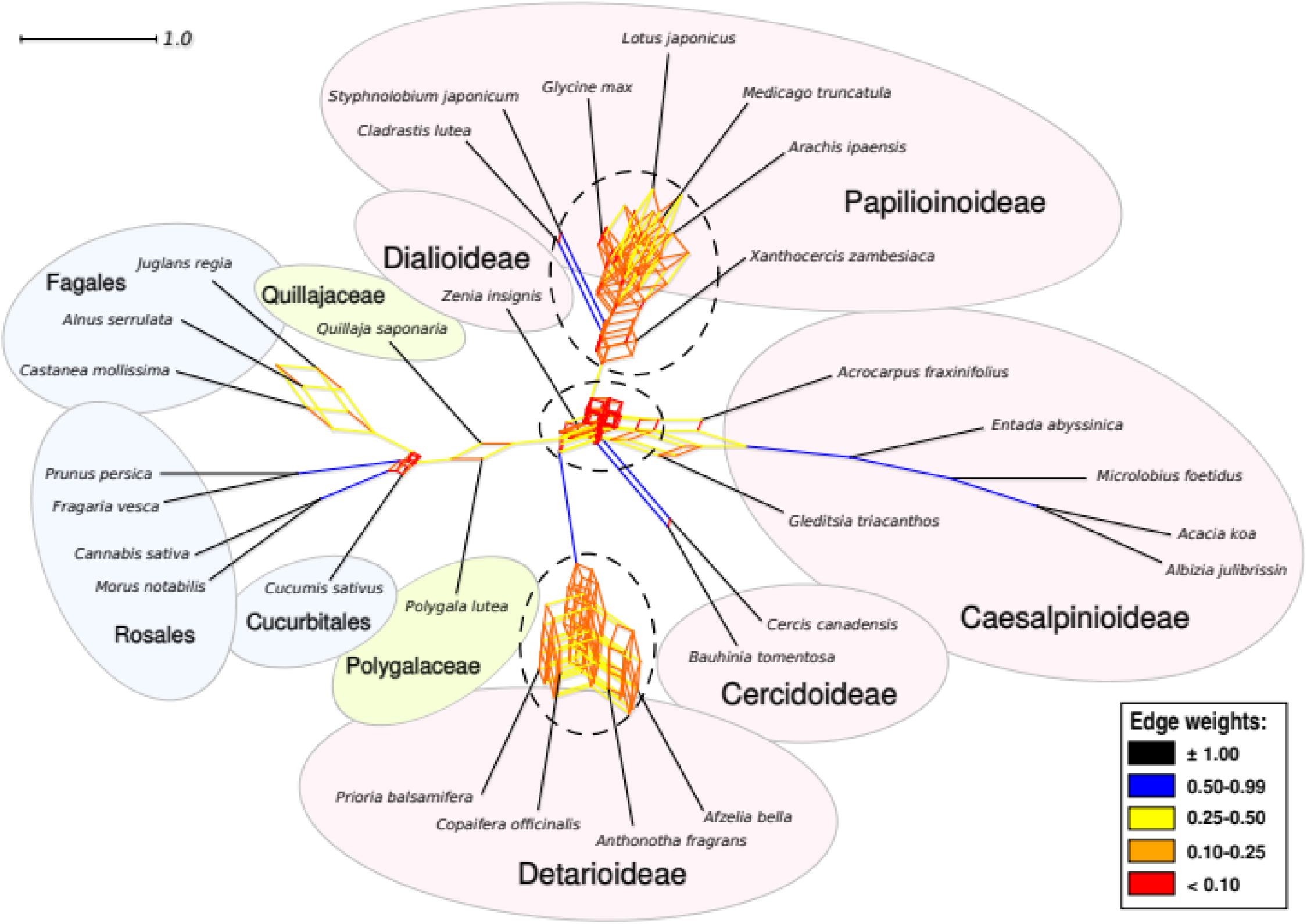
A filtered supernetwork shows tangles of gene tree relationships at the bases of the legumes, and subfamilies Detarioideae and Papilionoideae, that correspond to WGDs. The filtered supernetwork was inferred from the 1,103 1-to-1 ortholog gene tree set, only bipartitions that received more than 80% bootstrap support in gene tree analyses were included. Edge lengths and colours are by their weight, a measure of prevalence of the bipartition that the edge represents among the gene trees. Ellipses with dashed outline indicate increased complexity at putative locations of WGDs.

Rather than relying solely on ICA and quartet support values, we sought to evaluate in a more intuitive way how much support and conflict there is among gene trees for the deepest divergences in the legume family. For nodes labeled A-H in Figure 3A, we counted how often a bipartition that is equivalent to that node in the species tree is encountered across gene trees, and how often those bipartitions received at least 50 or 80% bootstrap support. We did this on all RT homologs (n=7,621) in which all subfamilies and the outgroup were represented by at least one taxon each, leading to 3,473 gene trees being considered. This shows that the legume family as a whole, and the four subfamilies for which more than one taxon was sampled (nodes C, D, G and H), are all found to be monophyletic across the majority of gene trees (Fig. 5A & Table 2), and those bipartitions mostly receive at least 50 or 80% bootstrap support. Nodes B, E and F, that is, the relationships among the subfamilies, are recovered in many fewer gene trees, especially when only considering bipartitions with at least 50 or 80% bootstrap support. For these nodes, we then checked how often the most important conflicting bipartitions were present (Figs 5B-D & Table 2). These conflicting bipartitions are each less prevalent than those found by the concatenated ML and Bayesian analyses as well as by ASTRAL, confirming that the recovered topology represents the relationships among legume subfamilies that is supported by the largest fraction of the genomic data used here. But it also shows that there is significant and well-supported gene tree conflict, in line with the complicated tangle and short edges observed in the filtered supernetwork at the base of the legumes (Fig. 4).

**Figure 5.**
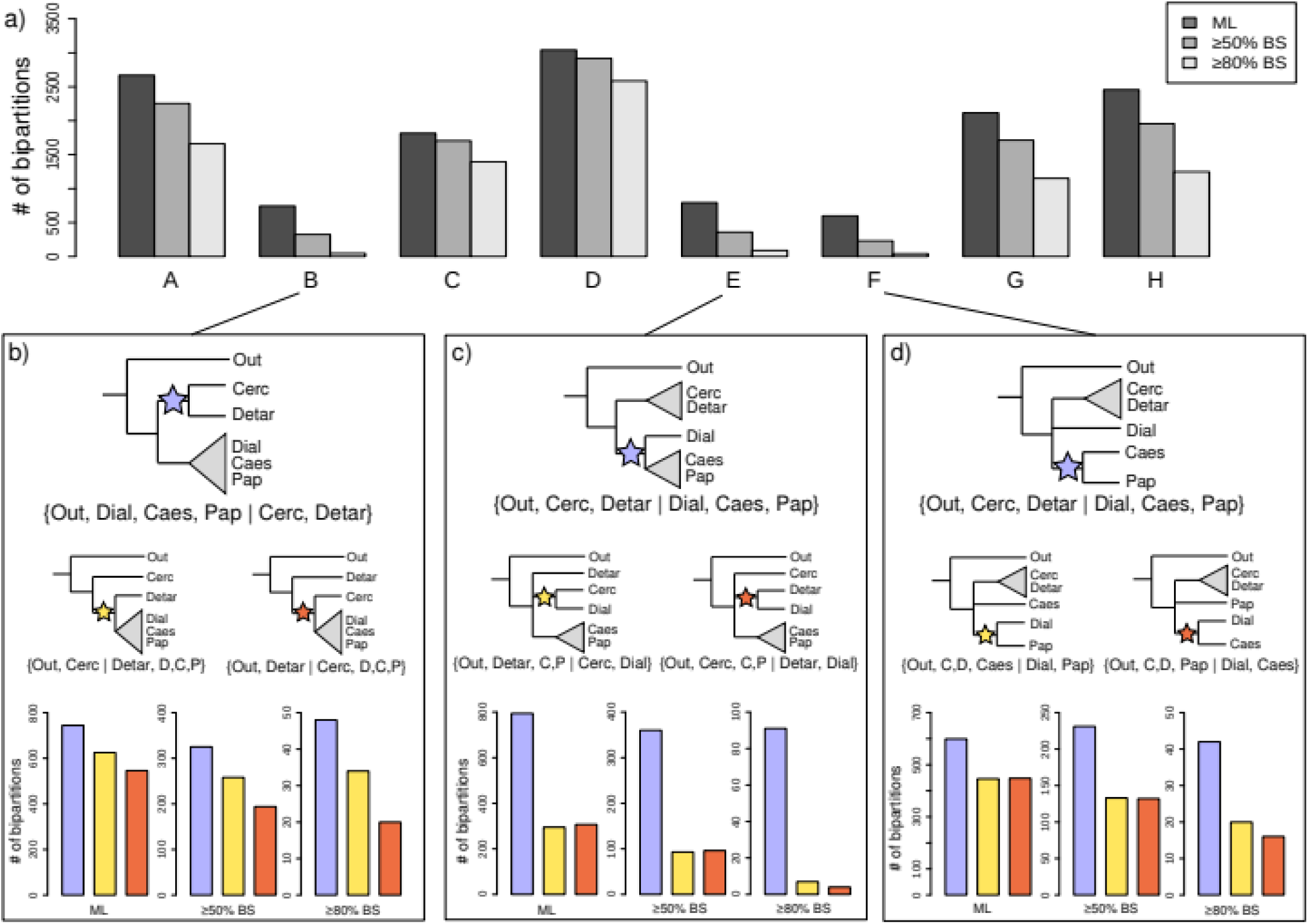
Leguminosae and its subfamilies are each supported by a large fraction of gene trees, in contrast to relationships among the subfamilies. (A) Prevalence of bipartitions that are equivalent to nodes A-H (see Fig. 3A), among the 3,473 gene trees inferred from the RT homolog clusters (including 1-to-1 orthologs) in which all five subfamilies and the outgroup were included. Numbers of bipartitions are shown as counted from the best-scoring ML gene trees as well as taking only bipartitions with more than 50 and 80% bootstrap support into account, as indicated in the legend. (B-D) Prevalence of bipartitions for nodes B, E and F plotted next to the most common alternative bipartitions. The locations of the stars in the illustrations indicate the internodes of the phylogeny that are equivalent to the bipartitions for which counts are plotted below, as counted from the ML estimates and for bipartitions with at least 50 or 80% bootstrap support. Colors of the stars correspond to the colors of the bars in the barplots.

### Inferring Phylogenetic Locations of WGDs

To map gene duplications over the species tree, we first removed fragmentary sequences and gappy sites from the 9,282 homolog clusters, after which 640 clusters with large amounts of missing data were eliminated. From trees that were inferred from the remaining 8,642 homologs, we extracted 8,038 rooted clades. Exemplar homolog trees with gene duplications are shown in Figure S8. We find significantly elevated numbers of gene duplications at several nodes where WGDs are hypothesized to have occurred, including the previously documented *Salix/Populus* clade (Tuskan et al., 2006) and one subtending Pentapetalae, consistent with the known *gamma* hexaploidization associated with that clade (Jiao et al., 2012) (Figs 3D & S9). For the Pentapetalae clade, many homologs show more than one gene duplication at that node, given that the number of duplications (1,901) is nearly twice as high than the number of homologs with duplications (1,105), as expected for two consecutive rounds of WGD. Part of these duplications may also stem from older events, since missing data for the three non-Pentapetalae taxa in our dataset could mean that we do not find duplicates of older events in these taxa. In the legumes, high numbers of gene duplications at particular nodes suggest that there were three early WGD events, one at the base of the family, and one each subtending subfamilies Papilionoideae and Detarioideae (Figs 3D & S9). When applying a bootstrap filter to the homolog trees (≥50% bootstrap support), numbers of gene duplications are considerably lower, but the pattern is the same (Figs 3D & S9). At the root of the family, the number of gene duplications drops from 1,646 to 99 when applying this bootstrap filter, in line with the difficulty of resolving the deepest dichotomies of the legume phylogeny. Notably, for the legume crown node we also find evidence for a significant part of homologs having had more than one gene duplication, because 1,646 duplications from only 1,229 homologs map on that node. This would suggest multiple rounds of WGD (e.g. Figs S8E & F), although some of these can be attributed to duplications in both paralog copies of genes duplicated at the *gamma* event, while for many others support values across the tree are low. For other hypothesized WGDs, the numbers of homologs with more than one duplication for those nodes are much lower, suggesting they involved a single round of WGD.

### Divergence Time Estimation

To establish whether the origin of legumes and the early WGD events are closely associated with the KPB, we performed clock dating in a Bayesian framework. Because the chloroplast phylogeny shows large root-to-tip length variation (Fig. 2), we refrained from using the chloroplast data to infer divergence time estimates, and instead rely on the better suited nuclear data for this purpose as suggested by Christin et al. (2014). We selected 36 relatively highly informative and clock-like nuclear genes and 20 fossil calibrations (Table 1 and Methods S1). The oldest definitive fossil evidence of crown group legumes is from the Late Paleocene, consisting of bipinnate leaves from c. 58 Ma (Wing et al., 2009; Herrera et al., submitted) and papilionoid-like flowers from c. 56 Ma (Crepet & Herendeen, 1992), representing Caesalpinioideae and Papilionoideae respectively. The older fossil woods with vestured pits, from the Early Paleocene of Patagonia (Brea et al., 2008) and the Middle Paleocene of Mali (Crawley, 1988), could represent stem relatives of the family (vestured pits are found in Papilionoideae, Caesalpinioideae and Detarioideae, so this is likely an ancestral legume trait). Based on this fossil evidence, c. 58 Ma can be considered the minimum age of the legume crown node. Molecular age estimates (95% HPD intervals) for the crown node range from 65.47-86.45 Ma and 73.46-81.18 Ma under the uncorrelated log-normal relaxed clock (UCLN) and the random local clock (RLC) models, respectively, to minima and maxima between 64.63 and 68.85 Ma under various fixed local clock (FLC) models (Table S3), the latter suggesting a close association of the origin of the legumes with the KPB (Fig. 6). Maximum clade credibility (MCC) trees for all clock analyses, with 95% HPD intervals indicated, are included in Supplementary Figures S10-17, and 95% HPD intervals for nodes A-H are listed in Table S3.

**Figure 6.**
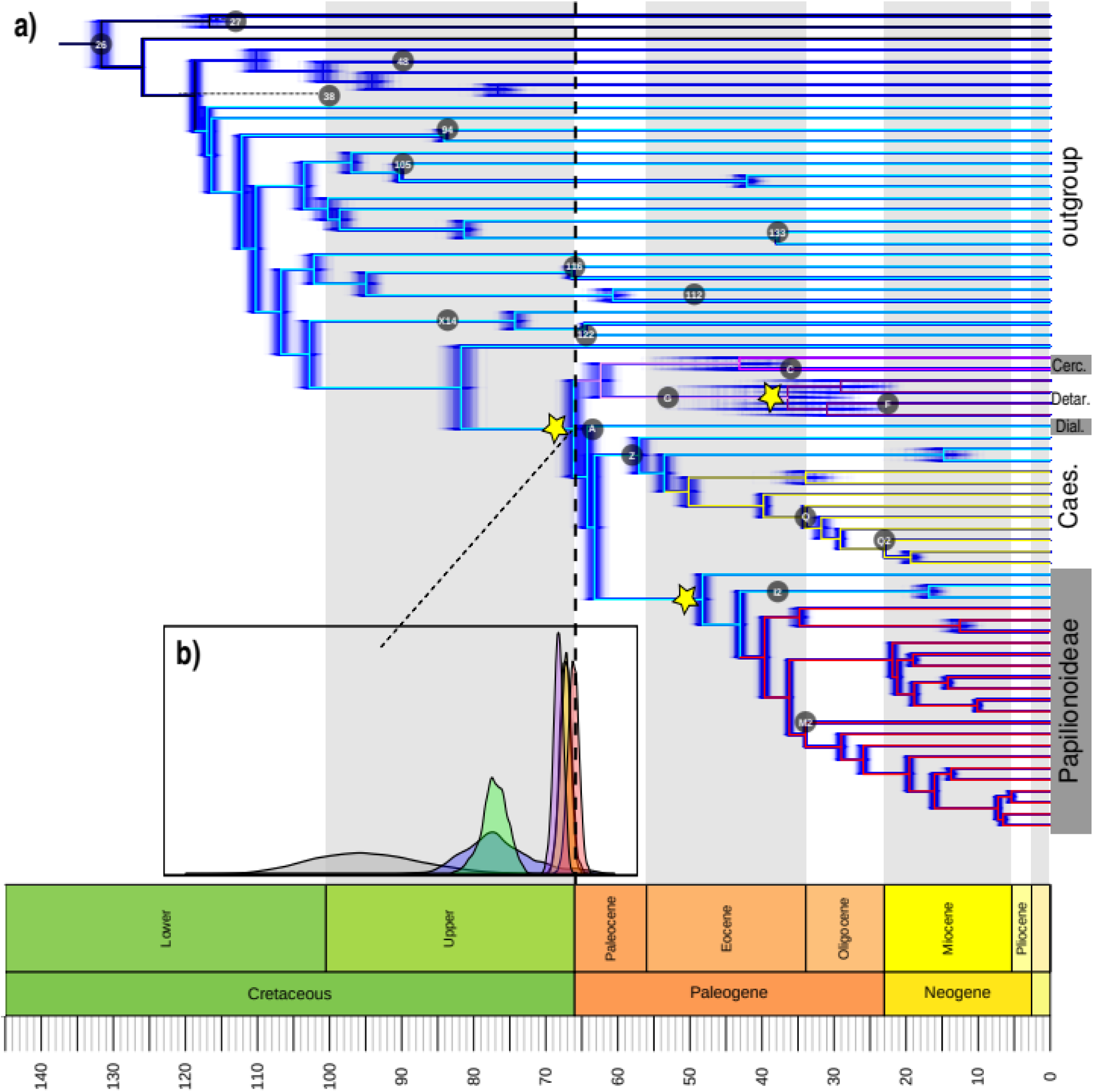
The origin of the legumes is closely associated with the KPB. (A) Chronogram estimated with 8 fixed local clocks (FLC8 model) in BEAST, with the clock partitions indicated by colored branches, from an alignment of 36 genes selected as both clock like and highly informative and hence well-suited for clock analyses. Blue shading represents 500 post-burnin trees (‘densitree’ plot) to indicate posterior distributions of node ages. Yellow stars indicate putative legume WGD events. Labelled circles plotted across the phylogeny indicate placement and age of fossil calibrations listed in Table 1. (B) Prior and posterior distributions for the age of legumes under different clock models. The marginal prior distribution is plotted in grey, UCLN in blue, RLC in green, STRC in purple and FLC3 in yellow, FLC6 in orange and FLC8 in red.

Placement of Eocene fossils of Detarioideae and Cercidoideae within the crown groups of those clades (Bruneau et al., 2008; Simon et al., 2009; de la Estrella et al., 2017), yields older crown group estimates for these clades. However, with these calibrations (alternative prior 1 in Table S4), a more than 10-fold higher substitution rate along the stem lineages of these two subfamilies relative to the rates within both crown clades is inferred (c. 8.82 × 10^−3^ vs 0.69 × 10^−3^ substitutions per site per million years, with identical rates estimated independently for Cercidoideae and Detarioideae; Fig. S18A). This rate is also nearly five times higher than the mean rate across the tree as a whole (1.54 × 10^−3^ substitutions per site per million years), while the crown clades are estimated to have rates about half as high as the mean. Analyses with the same clock partitioning but calibrated with Late Eocene *Cercis* fossils and Mexican amber (*Hymenaea*) as the oldest crown group evidence for Cercidoideae and Detarioideae, respectively, do not infer such strong substitution rate shifts, with all clock partitions across the phylogeny estimated to have a substitution rate ranging from 0.96 × 10^−3^ to 2.53 × 10^−3^ substitutions per site per million years (Fig. S18B). Either way, different placements of these fossils have little influence on the crown age estimates for the family in the FLC analyses (Figs S15 & S16, Table 3).

## Discussion

In this study, we present significant advances in our understanding of the origin and early evolution of the legume family. All the different species tree analyses of the nuclear genomic data yielded the same most likely topology with regards to relationships among subfamilies and the root of the legumes. Detailed evaluation of supporting and conflicting bipartitions across gene trees show that these relationships are the most prevalent, but we also found many conflicting bipartitions, and the chloroplast phylogeny also shows a different rooting of the family. Furthermore, we find evidence for three WGD events early in the evolution of the family, which further complicate the phylogenomic tangle at the base of the family. Time-calibration of the species tree suggests a close association of this complex origin of the legumes with the KPB. We discuss these findings and their relevance to understanding the evolution of the third largest angiosperm family, the likely complications caused by WGDs on phylogenetic inferences in deep time and the consequences of the KPB mass extinction event on plant evolution in the Cenozoic.

### Substitution Rate Variation in Legume Chloroplast Genomes

The chloroplast data set has the advantage of denser taxon sampling (including subfamily Duparquetioideae) compared to the nuclear genomic data. However, the chloroplast data are less useful for phylogenomic analysis, being effectively a single locus in the absence of recombination in plastid genomes. Furthermore, chloroplast genes have highly heterogeneous substitution rates across legumes, leading to a well-resolved topology in core Papilionoideae but poor resolution in other lineages, particularly Caesalpinioideae (Figs 2C & S1-4). It has long been known that there is significant variation in rates of chloroplast sequence evolution among plant lineages (Bousquet et al., 1992) and previous analyses of single chloroplast genes (Lavin et al., 2005) and legume chloroplast genomes (Dugas et al., 2015; Schwarz et al., 2017; Wang et al., 2018) have suggested substantial variation in rates of molecular evolution among legume lineages. Because branch model *D*_*n*_/*D*_*s*_ ratio tests do not provide evidence for different selective forces on photosynthesis genes across legumes, this pattern may rather be related to life-history strategies in the 50Kb-inversion clade, which includes many herbaceous plants of short stature (Lanfear et al., 2013) and shorter generation times (Smith & Donoghue, 2008), especially in the vicioid clade where the highest rates are found (Fig. 2C).

### Resolving the Deep-branching Relationships in the Leguminosae

The difficulty of obtaining resolution for the deep divergences in the legume family is in part caused by lack of phylogenetic signal in a large fraction of the sampled genes (pie charts in Figure 3A), with too few substitutions having accumulated along the deepest short internodes due to rapid early divergence of the six principal legume lineages. Lack of phylogenetic signal could potentially be explained by rapid diversification which, especially in combination with extinction of stem-relatives, causes alternations of long and short internodes, leading to “bushy” phylogenies that are extremely difficult to resolve (Rokas & Carrol, 2006). However, for a significant proportion of those genes that do have sufficient phylogenetic signal, we find strongly supported conflicting evolutionary histories. Putting aside methodological issues such as poor orthology inference for a number of genes, this conflict is likely to be caused by incomplete lineage sorting (ILS) (Pamilo & Nei, 1988; Maddison, 1997). Together with the complexity depicted in the supernetwork (Fig. 4), the strongly supported conflicting gene trees suggest that a fully bifurcating tree is an oversimplified representation of the initial radiation of the legumes. As we show here, genes have many different evolutionary histories across the early divergences of legumes (Table 2), while the species tree merely represents the dominant evolutionary history. In the case of complete lack of phylogenetic signal, or equally prevalent conflicting evolutionary histories without a single dominant one, this would constitute a hard polytomy, meaning (nearly) instantaneous speciation of three or more lineages, as demonstrated for Neoaves (Suh, 2016). Alternatively, a phylogenetic network can provide a better representation of evolutionary relationships when there is significant gene tree conflict. In the legumes, there does appear to be one dominant evolutionary history in the relationships among subfamilies, suggesting that the root of the family is strictly speaking not a hard polytomy. Nevertheless, the short internodes leading to lack of phylogenetic signal and significant conflict among gene trees at the base of the legumes suggest that the first few divergences in the family occurred within a short time span, leading to ILS. Indeed, strong gene tree conflict caused by ILS has been shown to be relatively common when internodes are short due to rapid speciation and this provides an explanation as to why many relationships are contentious (e.g. Pollard et al., 2006; Suh et al., 2015; Moore et al., 2017). In such cases, it is essential that phylogenomic studies explicitly evaluate conflicting phylogenetic signals across the genome. By taking into account alternative topologies that are supported by significant numbers of gene trees (Fig. 5) and inferring a phylogenetic network (Fig. 4), the phylogenomic complexity of the initial radiation of the legumes is revealed.

### Locating WGD Events on the Phylogeny

Numbers of gene duplications mapped onto the species tree provide evidence for three WGD events early in the evolution of the legume family, one shared by the whole family, plus independent nested WGDs subtending subfamilies Detarioideae and Papilionoideae. We note that several nodes that immediately follow the most likely locations of hypothesized WGD events also show elevated numbers of gene duplications (Figs 3D & S9). This is most likely caused by missing data for some taxa. For example, sequences for *Xanthocercis zambesiaca*, *Cladrastis lutea* and *Styphnolobium japonicum* are derived from transcriptomes, while in the core Papilionoideae, several accessions are represented by fully sequenced genomes and therefore have higher gene sampling. Alternatively, paralog copies for a subset of genes could have been lost in lineages outside the core Papilionoideae. These gene sampling issues mean that a considerable number of gene duplications are likely to be mapped onto the second and third divergences in the subfamily, even though they probably stem from the same WGD event shared by the subfamily as a whole. Similar patterns are apparent at the bases of the legumes and of Pentapetalae (Figs 3D & S9). At the bases of subfamily Caesalpinioideae and the Mimosoid clade, we also find modestly elevated numbers of gene duplications, but fewer than for the three main duplication events (Figs 3D & S9). This could indicate a partial genome duplication shared by all Caesalpinioideae and another one shared by all mimosoids. Alternatively, it could reflect higher gene coverage in the mimosoid transcriptomes relative to the other Caesalpinioideae, in which case many of the gene duplications currently depicted as subtending the Mimosoid clade should potentially map at the base of the Caesalpinioideae. It is therefore possible that another WGD has occurred at the base of the Caesalpinioideae, as suggested by Cannon et al. (2015), but the rather low numbers of gene duplications inferred from our data cannot be considered as strong evidence for that. Cannon et al. (2015) also hypothesized another WGD early in the evolution of subfamily Cercidoideae, shared by Cercis and Bauhinia. Our results do not support this, and furthermore, in Bauhinia s.l. (Cercidoideae) the most common haploid chromosome number is n=14, while Cercis has n = 7. This suggests that an early WGD in Cercidoideae was not shared by Cercis. This is further supported by a densely sampled phylogenetic analysis of the LegCyc gene in Cercidoideae, which is duplicated in all Cercidoideae except Cercis, the sister group to the rest of the subfamily (Carole Sinou, unpublished data). Cannon et al. (2015) further suggested that the ancestral legume most likely had a haploid chromosome number of n = 6 or 7 and had independently doubled in most lineages to arrive at n = 14, the haploid chromosome number that is most commonly found across legume subfamilies except Detarioideae (n = 12) and core Papilionoideae (Cannon et al., 2015: Fig. 1; chromosome counts for Duparquetia are not available). This would imply that Cercis, with n = 7, would have retained the ancestral haploid chromosome number. Indeed, given our results it is likely that the mrca of Cercidoideae and Detarioideae would have had a haploid chromosome number of 6 or 7, followed by independent WGDs in Bauhinia s.l. and Detarioideae to arrive at n = 14 and n = 12, respectively. However, the mrca of Dialioideae, Caesalpinioideae and Papilionoideae most likely had a haploid chromosome number of n = 14, followed by reductions in chromosome number in Chamaecrista and Papilionoideae (Cannon et al., 2015: Fig. 1). Even after an additional WGD in Papilionoideae, extant members of the subfamily still have chromosome numbers <14 (Cannon et al., 2015: Fig. 1), suggesting extensive genomic rearrangement. That leaves the chromosome number of the mrca of all legumes uncertain, being either n = 6 or 7, or n = 14, suggesting either chromosome number reduction in some lineages, or potentially inheritance of different ploidy levels in different lineages from an ancestral polyploid complex. In conclusion, we find evidence that supports many of the findings of Cannon et al. (2015), but our results suggest an additional WGD event that is shared by all legumes, in line with the findings of Wong et al. (2017). Our study expands the taxon sampling of Cannon et al. (2015), but has the same limitation in that a large number of accessions are based on transcriptome data and are thus not sampling complete exomes. Denser sampling of completely sequenced legume genomes will be needed to resolve the number and placement of WGD events with higher confidence, precision and accuracy.

### Estimating the Timeline of Legume Evolution

Our divergence time analyses update previous analyses of Lavin et al. (2005), Bruneau et al. (2008) and Simon et al. (2009), and provide, to our knowledge, the first divergence time estimates for legumes based on nuclear genomic data as well as the first molecular clock dating estimate for the crown age of the legumes. The age estimates under the FLC models and the strict clock model are mostly rather similar, but the RLC and UCLN models, that relax the clock assumption more, lead to older divergence time estimates. By allowing independent substitution rates on all branches, these models are potentially overfitting the data, to attempt to satisfy the marginal prior on node ages (Brown & Smith, 2017). As inferred from analyses run without data, the marginal prior that is constructed across all nodes of the tree, can be considered as “pseudo-data” (Brown & Smith, 2017), derived from the node calibration priors (based on fossil ages) and the branching process prior (constant birth-death model in our case), and should therefore not be overly informative on node ages. FLC and strict clock models lend greater weight to the molecular data and can overrule the marginal prior distributions on divergence times whilst still respecting hard maximum and minimum bounds of the fossil constraints on calibrated nodes, as suggested by our results. It is also clear from running analyses without data, that the marginal age prior on the (uncalibrated) crown node of the legumes is rather poorly informed, with the 95% HPD interval between 79.37-109.20 Ma (Fig. 6B), the minimum being much older than the oldest legume fossils, presumably caused by overly conservative maximum bounds on calibrated nodes (Phillips, 2015). UCLN and RLC analyses also inferred relatively high substitution rates for a few deep branches in the outgroup during the Lower Cretaceous, relative to the more derived and terminal branches of the tree (Figs S10 & S12), presumably to satisfy the poorly informed marginal priors. Phillips (2015) suggested that setting less conservative maxima on priors could remedy this problem, but our analysis with such prior settings shows little effect (Fig. S11), with some of the deepest branches still having much higher estimated substitution rates. Since there is no evidence, nor any reason to assume, that substitution rates along those branches should be elevated relative to terminal branches, we conclude that this is indeed caused by overfitting of rate heterogeneity across branches under the influence of the marginal prior. Furthermore, the RLC analyses fitted c. 45 local clocks across the phylogeny, a rather high number relative to the total of 142 branches in the tree (implying a separate clock for every 3 branches), which is also indicative of overfitting. At the same time, this could be seen as evidence that the data are not the product of clock-like evolution, but it becomes difficult to estimate how much the clock deviates if the marginal prior on node ages is too influential. A more pragmatic approach is to use FLC analyses, by defining local clocks based on root-to-tip length distributions across clades and pruning outlier taxa (see Methods and Fig. S19). This approach accounts in large part for the violation of the molecular clock but it does not relax the clock to the extent that the marginal prior on node ages is given excessive weight relative to the molecular signal. Furthermore, because the genes we selected for divergence time estimation are reasonably clock-like and highly informative, it is desirable that these data inform the node ages with sufficient weight. One drawback of using this approach is that the relatively large amount of sequence data in combination with the FLC model results in estimates that appear unrealistically precise, and the discovery of new fossils may well prove the legumes to be slightly older. Nevertheless, the evidence presented here suggests that the legume crown age dates back to the Maastrichtian or Early Paleocene, likely within one or two million years before or after the KPB, although such high precision is not warranted due to the idiosyncrasies of the molecular clock.

Polyploidy (Senchina, et al., 2003) as well as the KPB itself (Berv & Field, 2018), have been implicated as potentially causing transient substitution rate increases, raising the possibility that substitution rates during the early evolution of the legumes could have deviated temporarily but markedly from the "background" rate of Cretaceous rosids. This would render the ages inferred for the first few dichotomies as well as those of the subfamilies less certain. The age estimates inferred for these nodes rely in large part on the assumption that the substitution rate did not vary significantly within the different clock partitions, and most importantly within the rosid partition which includes most of the branches along the backbone of the family and the stem lineage subtending it. The WGD events along the stem lineages of the family, and subfamilies Papilionoideae and Detarioideae could have affected substitution rates along those branches. By selecting for smaller stature and shorter generation times and reducing population sizes (Berv & Field, 2018), the KPB could additionally have resulted in increased rates along some or all of the stem lineages of the subfamilies, and, in the case of "hard" explosive diversification after the KPB, perhaps also along the legume stem lineage. A third factor that could influence node age estimates along the backbone of the family, is the strong gene tree incongruence observed for nodes B, E and F (Fig. 5), which is also observed among the 36 genes that were used for time-scaling. The divergence time analyses need to accommodate this incongruence within a single topology, meaning that additional substitutions need to be inferred for conflicting gene trees, which can inflate the branch lengths between rapid speciation events (Mendes & Hahn, 2016). Taken together, these three factors could mean that the timeframe for the early evolution of the legumes appears inflated in our results, with (some of the) subfamily ages likely being slightly older than estimated here, as well as divergence of the subfamilies happening nearly instantaneously (hence the gene tree incongruence and lack of phylogenetic signal), rather than spanning the c. 3 - 5 million years inferred here (Figs 6 & S10-17). Potentially, even the legume crown age could be slightly older due to the effects of polyploidy, but not due to the KPB, because if the crown is older, the stem lineage would not have crossed the KPB.

Different interpretations of Eocene fossils of Cercidoideae and Detarioideae (see Methods S1) lead to very different crown age estimates for these clades. As expected, this also leads to very different substitution rates along the stem lineages of these subfamilies, whit rates increasing 10-fold when interpreting these fossils as crown group members. While it cannot be ruled out that the stem lineages of Cercidoideae and Detarioideae experienced such markedly elevated substitution rates, it is unlikely that rates were five times higher relative to the rest of the eudicots across all 36 nuclear genes analysed, especially as these genes were chosen because of their approximately clock-like evolution, and given that these two clades comprise long-lived woody perennials. The idea that molecular information from extant taxa could inform that particular fossils are too old to belong to a crown clade is controversial. However, the test we have performed here is similar to the cross-validation method proposed by Near et al. (2005), which also uses molecular data to discover fossil calibration points that do not fit well with a larger set of fossils. Favouring those calibrations that do not lead to extreme substitution rate shifts is more parsimonious, and we believe that additional evidence is necessary to justify the inference of such a strong shift in substitution rates as that observed in the FLC8 analysis with alternative prior 1 (Fig. S16). While there seems little doubt that the Early Eocene fossils from the Mahenge in Tanzania and the Paris Basin in France do represent Cercidoideae and Detarioideae, the extreme substitution rate heterogeneity implied by their treatment as crown group members suggest that they may better be reinterpreted as stem-relatives of these subfamilies (see additional discussion about the affinities of these fossils in Supplementary Methods S1).

### The Impact of the KPB on Plant Diversification

The impacts of the KPB mass extinction event on plant diversity are the focus of debate, with several studies claiming that extinction was less severe for plants than across marine and terrestrial faunas (Nicholls & Johnson, 2008; Cascales-Miñana & Cleal, 2014; Silvestro et al., 2015). However, our results suggest that the massive KPB turnover event likely played a critical role in the evolution of plant taxa. Our analyses indicate that the origin of crown group legumes is closely associated with the KPB. The analyses employing FLCs even suggest that potentially only a single legume ancestor crossed the KPB to give rise to the six main lineages during the early Paleocene, conforming to a “hard explosive” model. However, across the different analyses, part of the posterior density of the crown age estimate falls in the late Maastrichtian, suggesting a “soft explosive” model, with the six main lineages diverging in the Late Cretaceous and crossing the KPB, giving rise to the crown groups of the modern subfamilies in the Cenozoic. These different explosive models have been used to describe the origin and early diversification of the placental mammals, although other studies have lent support to “short fuse” or “long fuse” models (summarized in Phillips, 2015: Fig. 1). For birds, the timing of diversification relative to the KPB has also been controversial (Ksepka & Phillips, 2015), but it now appears likely that the Neoaves underwent explosive radiation from a single ancestor that crossed the KPB (Suh, 2016). Apart from Placentalia and Neoaves, recent studies on frogs (Feng et al., 2017) and fishes (Alfaro et al., 2018) have also demonstrated rapid diversification following the KPB, suggesting this is a common pattern across many terrestrial and marine animal groups. We present here, to our knowledge, the first example of a major plant family whose origin and initial diversification appears to be closely linked to the KPB. This is notable because a recent family-level paleobotanical study suggested that the KPB did not constitute a mass extinction event for plants (Cascales-Miñana & Cleal, 2014). Phylogenetic studies in some plant families originating in the Cretaceous also lack any evidence of a significant effect of the KPB on diversification (e.g. Annonaceae (Couvreur et al., 2011a) and Arecaceae (Couvreur et al., 2011b)), except for the smaller plant family Menispermaceae (Wang et al., 2012), which shows increased diversification following the KPB. In contrast, fern diversification appears to have been strongly affected, with some groups of ferns showing much reduced diversity in the Cenozoic compared to earlier times (Lehtonen et al., 2017), and especially epiphytic groups of ferns showing increased diversification rates since the KPB (Schuettpelz & Pryer, 2009). Furthermore, the generic-level study of Silvestro et al. (2015) showed high extinction rates for non-flowering plant groups during the late Cretaceous, and elevated origination rates for angiosperms during the Paleocene, in line with the pattern we observe for the origins of legume diversity. Thus, even if extinction was less severe for plants than for animals at the KPB, the Paleocene was nevertheless a time of major origination of lineages across biota, and we expect further examples of KPB-related accelerated plant diversification to be discovered when inferring larger angiosperm timetrees.

### Implications for our Understanding of the Evolution of Legume Diversity and Traits

Rapid divergence of the six main lineages of legumes is clearly relevant to our understanding of the evolution of legume diversity and the appearance of key traits. Over the last few decades, the prevailing characterization of legume evolution has been that of mimosoids and papilionoids as “derived” clades that evolved from a paraphyletic “grade” of caesalpinioid legumes (e.g. LPWG, 2013). However, we show that all six subfamilies diverged across a short time span after the origin of the legume crown group, with long stem lineages subtending each subfamily, suggesting that none of the modern subfamilies should be seen as diverging earlier or later than any other. The complex phylogenomic paleopolyploid tangle documented here means that it will be extremely difficult to reconstruct trajectories of trait evolution across the first few divergences within the family. For example, it is not clear how to understand the evolution of floral diversity across the family and what the ancestral legume flower would look like. That makes it questionable, for example, to what extent the specialized and strongly canalized zygomorphic papilionoid flowers are derived within the family. Fossil papilionoid flowers from the Paleocene (Crepet & Herendeen, 1992) are among the oldest evidence of the family in the fossil record. The higher morphological diversity of flowers in other subfamilies may well have evolved in parallel or even later than the papilionoid flower, given the crown age estimates that we find in Bayesian clock analyses (Figs 6, S10-17 and Table S3).

While we are not able at this point to confidently distinguish between a “hard” or “soft explosive” model of early diversification of the family, it is clear that the early radiations of the legume subfamilies all occurred in the Cenozoic. While stem age estimates of each subfamily are remarkably close to each other, crown age estimates are strikingly different (but see the discussion above on potential effects of polyploidy and the KPB on substitution rates and ages of subfamilies). Caesalpinioideae are found to have the oldest crown age (late Paleocene), followed by Papilionoideae with a crown age in the Early Eocene. Both of these subfamilies therefore likely diversified considerably during the PETM and Eocene climatic optimum, when tropical forests extended far into the Northern Hemisphere. This is in line with the numerous legume fossil taxa known from the Eocene of North America, often of uncertain affinities, but with a majority ascribed to Caesalpinioideae and Papilionoideae (Herendeen, 1992). There is also fossil evidence of Early and Middle Eocene stem-relatives of Cercidoideae and Detarioideae (as discussed above and in Methods), but their crown group divergences are most likely placed in the Late Eocene or Oligocene. Our results suggest extinction of stem-relatives of these two subfamilies, most likely related to Late Eocene and Oligocene cooling, and subsequent diversification of the crown groups during the Oligocene and Miocene, when both groups become diverse at several fossil sites (e.g. Wang et al., 2014; Lin et al., 2015; Poinar, 1991; Poinar and Brown, 2002). Although it remains uncertain whether the crown group divergence of Detarioideae occurred in the (Late) Eocene or the Oligocene, the younger age of the subfamily inferred here contrasts with previous views of the evolutionary trajectories of this subfamily dating back into the Paleocene, comprising relatively slowly evolving lineages (de la Estrella et al., 2017), and with Amazonian subclades within Detarioideae conforming to the museum model of tropical rainforest diversification (Schley et al., 2018). This has important implications for our understanding of the origins of tropical African plant diversity, since Detarioideae dominate the canopy of many equatorial African rainforests, as well as being an important group in African savannas (de la Estrella et al., 2017). Our results for Detarioideae suggest that the extant diversity in tropical Africa, in particular the large diversity in tribe Amherstieae, is of relatively recent origin following a major turnover event at the Eocene-Oligocene boundary, which also affected other plant groups such as palms (Pan et al., 2006). This more recent diversification of detarioids is also more in line with the widely proposed recent assembly of the savanna biome (Cerling et al., 1997; Bouchenak-Khelladi et al., 2009; Maurin et al., 2014).

### The Added Complications of Paleopolyploidy on Evolutionary Inferences in Deep Time

The recent proliferation of genomic data is revealing just how prevalent repeated WGDs have been in the history of the angiosperms (e.g. Wendel, 2015; Soltis et al., 2016; Yang et al., 2018) and how many large angiosperm clades are characterized by genome triplications (e.g. Pentapetalae, Brassicaceae, Asteraceae, Solanaceae). Here we show that there were also multiple WGDs during the early history of the legumes, including a WGD subtending the family as a whole. It has been suggested that angiosperm WGDs are non-randomly distributed through time and significantly clustered around the KPB (Fawcett et al., 2009; Vanneste et al., 2014; Lohaus & Van de Peer, 2016). The WGD that we identify that is shared by all legumes is also temporally close to the KPB (Fig. 6), lending further support to the idea that polyploid survival and establishment were enhanced at or soon after the KPB with its associated rapid turnover of lineages (Lohaus & Van de Peer, 2016; Levin & Soltis, 2018). WGDs have also been hypothesized to trigger accelerated rates of lineage diversification at least in some lineages, albeit potentially after a time lag (Schranz et al., 2012; Tank et al., 2015; Landis et al., 2018; Smith et al., 2018a). The three legume WGDs we detected are each followed by rapid divergence of lineages as indicated by short internodes (Figs 2, 3 & 6). Polyploidy could have helped ancestral legumes and other plant lineages to both survive the mass extinction event and rapidly diversify owing to differential gene loss and other processes of diploidization (Adams & Wendel, 2005; Dodsworth et al., 2016). Increased polyploid speciation and reduced diploid speciation in the wake of the KPB (Levin & Soltis, 2018) would then lead to over-representation of these WGD-derived lineages in the extant flora and clustering of WGDs around the KPB. On the other hand, many paleopolyploidy events that significantly pre- and post-date the KPB are known (e.g. Angiospermae (Jiao et al., 2011), Pentapetalae (Jiao et al., 2012), Salicaceae (Tuskan et al., 2006), Caryophyllales (Yang et al., 2018), *Gossypium* (Wendel, 2015)), including in legumes (e.g. *Glycine*, Genisteae, the *Leucaena* group, *Vachellia*), and more extensive sampling of recently diversified groups may well reveal a weaker pattern of clustering around the KPB.

The WGD events subtending all legumes and subfamilies Detarioideae and Papilionoideae are likely to have contributed to the difficulties of obtaining phylogenetic resolution for the deep nodes in these clades (Cardoso et al., 2012 & 2013; de la Estrella 2018). WGDs may have promoted increased lineage diversification rates resulting in short internodes and ILS. If the polyploidy event happened some time before the first divergences in the legume family, or in the case of allopolyploidy, this could have led to divergent gene copies prior to lineage splitting which should make orthology detection easier. However, if the polyploidy event happened shortly before rapid cladogenesis, potentially a large fraction of paralogous gene copies would not have diverged at this point, making orthology detection challenging. In both cases, paralogous or homoeologous gene copies will have subsequently been differentially lost, pseudogenized or sub- or neo-functionalized, further complicating correct orthology detection. Together with ILS, this could explain the large fraction of gene trees supporting alternative topologies at the root of the legumes. It is notable that several other large plant clades, such as Pentapetalae (Zeng et al., 2017), Asteraceae (Barker et al., 2016; Huang et al., 2016) and Brassicaceae (Couvreur et al., 2010; Huang et al., 2015), also appear to show similar lack of resolution in clades subtended by WGDs to that revealed here for the legume family and subfamilies Papilionoideae and Detarioideae. This suggests that the association of polyploidy with rapid divergence, which leads to a lack of phylogenetic signal and gene tree conflict, is potentially a common feature in the evolution of angiosperms and the origination of major plant clades.

A large number of homolog clusters do not show gene duplications at the base of the legumes or any of the subfamilies, suggesting that loss of paralog copies is widespread, as observed for ancient WGDs more generally (Adams & Wendel, 2005; Dehal & Boore, 2005; Brunet et al., 2006; Scannel et al., 2007). If many of those losses occurred along the stem lineages of the six subfamilies after their divergence, this could lead to different paralog copies being retained in different lineages, adding to conflict among gene trees. Loss of paralog copies along stem lineages of subfamilies will also make it difficult to distinguish whether a gene duplication corresponds to the WGD shared by all legumes, or whether it represents a nested WGD such as those subtending Detarioideae and Papilionoideae. Lack of support in those homolog trees showing gene duplications further complicates this issue, making it potentially extremely challenging to accurately reconstruct the history of WGDs. Given these difficulties, sampling a wider range of complete genomes will be important, since with transcriptome data it is unknown whether duplicate gene copies are lost or simply not expressed in the tissue from which the RNA was extracted. Furthermore, increased taxon sampling will help to counteract negative impacts of missing data, since particular duplicate gene copies may have been lost in all species sampled here, but not necessarily across the whole clade or subfamily which those species represent. Despite all these complications, a clear pattern of either high or low numbers of gene duplications is observed when mapping duplications from 8,038 extracted clades from homolog trees across the species tree (Figs 3D & S8). This suggests that when summarizing gene duplications over a sufficiently large data set, it is still possible to make sense of the confusing topological differences that are observed and hence to accurately map WGD events. This leads us to propose the hypothesis presented in Figure 7 to reconcile the complicated topological patterns observed across gene trees. In this hypothesis, the six major legume lineages (i.e. subfamilies) diverged rapidly one after another from a polyploid ancestor. The different gene copies would still be nearly identical at the moment of cladogenesis and would diverge into paralog copies in each lineage independently, making it impossible to infer relationships between paralog copies from different subfamilies, consistent with lack of phylogenetic signal in most clusters. Coupled with differential loss of paralog copies, the diversity of topologies and the lack of support that we observe in homolog trees is exactly what would be expected from the sort of evolutionary history depicted in Figure 7.

**Figure 7.**
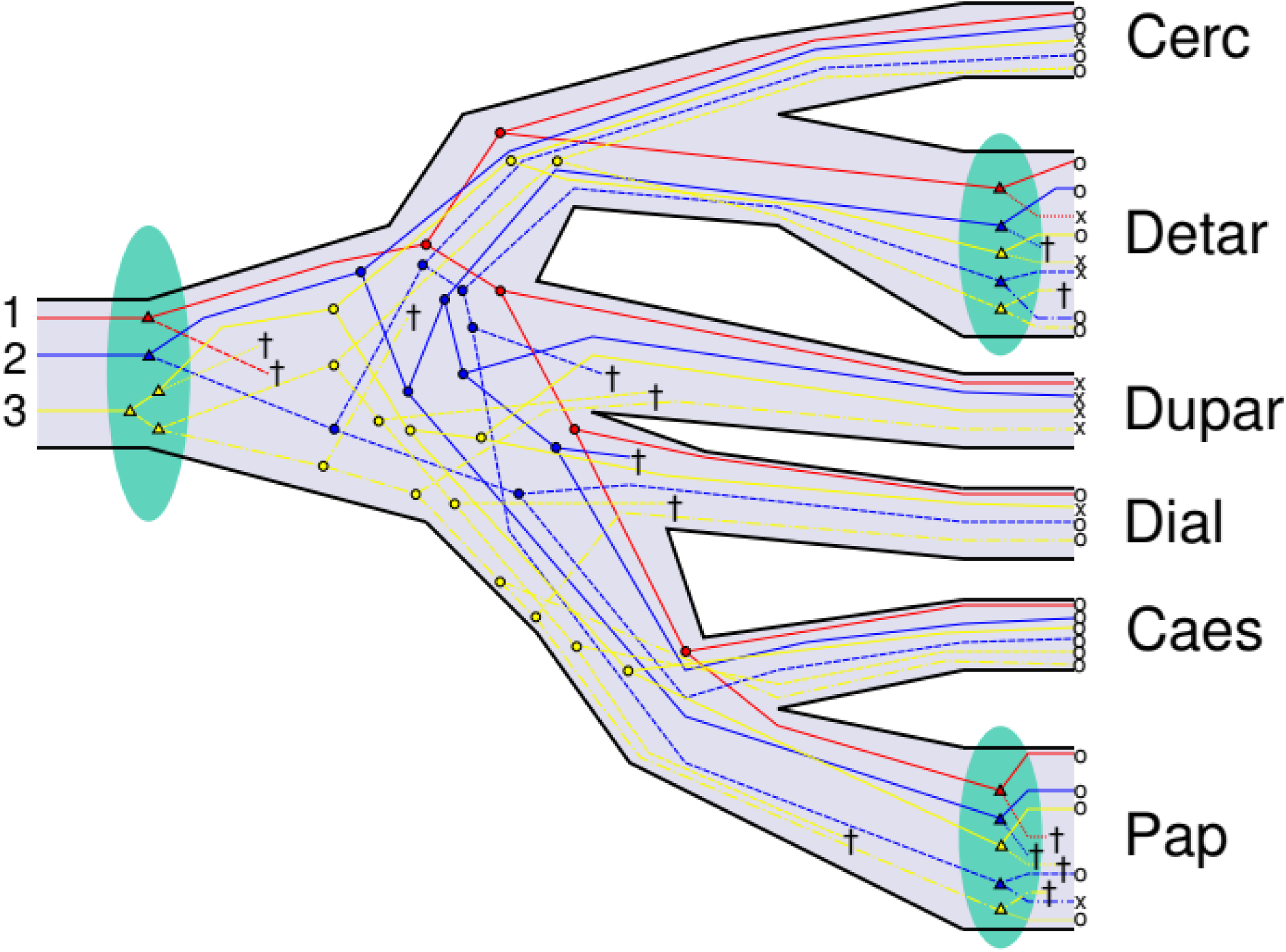
Differential loss of paralog copies combined with ILS leads to complex patterns of gene tree evolution. Gene trees 1, 2 and 3 (red, blue and yellow, respectively) are examples of increasingly complex hypothetical evolutionary gene histories, as reconciled with the species tree. Gene 1 loses one paralog copy prior to speciation, and the remaining copy yields the species tree topology in the absence of ILS. Gene 2 is modelled on the homolog cluster2941_1rr_1rr (Fig. S8D), where both duplicated copies are lost or not sampled in a few lineages and there is also ILS. Gene 3 is modelled on the homolog cluster544_1rr_1rr (Fig. S8F) and shows a hypothetical evolutionary history where two rounds of pan-legume WGD occurred in quick succession, with different paralog copies lost either early or late in some lineages and there is also ILS. Blue ovals indicate WGD events, triangles indicate gene duplications and circles indicate coalescences. † = gene loss, o = sampled, x = not sampled, Cerc = Cercidoideae, Detar = Detarioideae, Dupar = Duparquetioideae, Dial = Dialioideae, Caes = Caesalpinioideae and Pap = Papilionoideae.

A polyploid ancestor reconciles the complex patterns of gene duplications observed in the homolog clusters, suggesting we have six legume lineages derived from a recently polyploidized ancestor. This raises a number of important questions: Did the polyploidization event involve hybridization, leading to allopolyploidy? Was the ancestor tetraploid or did it have a higher ploidy level? Did all six lineages inherit the same ploidy level? Alternatively, given that polyploidization results in immediate reproductive barriers, perhaps divergence of these six lineages was even facilitated by differing ploidy levels, with all modern legume taxa derived from an ancestral polyploid complex?

These questions are difficult to answer for an event that occurred 66 Ma and for which much of the evidence has been obscured by subsequent genome reorganization and loss of the large majority of duplicate gene copies. Over such timescales, it appears nearly impossible to distinguish between autopolyploidization or allopolyploidization between two species that had only recently diverged, or multiple recurrent WGDs in a polyploid complex, or even to disentangle the impacts of possible reticulation from the effects of ILS. On the one hand, a hybridization event involving WGD could explain the strong gene tree conflict that we observe. However, equally this conflict could be explained by ILS alone. The first few divergences within the family occurred within less than 5 Myr (Fig. 6 & S10-17), and this is probably an overestimate due to gene tree incongruence (Mendes & Hahn, 2016). With a sufficiently large effective population size, the majority of loci would not yet have reached reciprocal monophyly over such a short time. Furthermore, the lack of resolution among the different gene copies in the majority of homolog trees suggests that genes did not diverge significantly prior to the WGD, therefore ruling out the possibility of allopolyploidization of two divergent lineages.

We hypothesize a polyploid ancestor of all legumes, but the ploidy level of this ancestor remains uncertain. Some of the gene trees suggest that multiple rounds of WGD occurred at the base of the legumes, prior to further WGDs that occurred independently in subfamilies Detarioideae and Papilionoideae (Fig. S8 E&F). Indeed, of the 794 homolog trees in which pan-legume duplications occurred, for 166 trees more than one duplication was mapped to the legume crown node. Some of these homolog clusters have low support values, so not all of them lend strong support to multiple rounds of WGD. Nevertheless, many of them clearly show more than two well supported duplicated clusters per subfamily, including for subfamilies other than Detarioideae and Papilionoideae. Therefore, the possibility of a hexaploid or octoploid legume ancestor, akin to events in Angiospermae (Jiao et al., 2011), Pentapetalae (Jiao et al., 2012), Asteraceae (Huang et al., 2016) and cotton (*Gossypium*) (Paterson et al., 2012), should also be considered given the evidence presented here. To further enhance knowledge on legume molecular biology and genome evolution, an obvious next step will be to sequence multiple complete genomes for all six legume subfamilies and the other Fabales families, something that will be forthcoming as part of the 10KP initiative (Cheng et al., 2018). This would potentially make it possible to disentangle the early genome evolution of legumes by comparing conserved synteny blocks, detecting genomic rearrangements and reconstructing chromosome evolution and the ancestral legume karyotype, as has recently been done for vertebrates (Sacerdot et al., 2018) and birds (Damas et al., 2018), as well as providing ample other opportunities to further enhance our understanding of legume evolution and diversification.

Ancient polyploidy not only provides a possible explanation for the difficulties in resolving the root of the legumes, it could also explain the sudden appearance of diverse legume fossil taxa in the Paleogene. A polyploid ancestor of all legumes would have provided a much expanded genomic substrate for rapid evolution and diversification of legume traits, with further rounds of genome duplication leading to an even further expanded genomic evolutionary substrate independently in Papilionoideae and Detarioideae and potentially several other legume lineages. In this sense the parallels to the sudden rise of the angiosperms (Sanderson, 2015) are even more compelling given that angiosperms are also subtended by one or two ancient WGD events (Jiao et al., 2011; Ruprecht et al., 2017).

### Concluding Remarks

It is becoming increasingly clear that the origin and early evolution of the legumes followed a complex scenario with multiple nested polyploidy events, and rapid divergence of the six main lineages against the background of a mass extinction event that led to major turnover in the Earth’s biota and biomes. WGD likely contributed to the survival and evolutionary diversification of the legumes in the wake of the KPB mass extinction event, and contributed to the rise to ecological dominance of legumes in early Cenozoic tropical forests. At the same time, these events make it more difficult to reconstruct aspects of the early evolutionary history of the clade, including evolutionary relationships, divergence time estimates and the phylogenetic location of the WGD events themselves. The similarities between legumes and other major Cenozoic clades such as mammals and birds are striking. All three of these prominent Cenozoic clades show recalcitrant basal polytomies and parallel trajectories of rapid early divergence closely associated with the KPB, further emphasizing the importance of the KPB mass extinction event and the earth system succession that followed in its aftermath (Hull, 2015) in shaping the modern biota.

## Acknowledgements

We thank the S3IT of the University of Zurich for the use of the ScienceCloud computational infrastructure and the Functional Genomics Center Zurich (FGCZ) for library preparation and sequencing.

## Online Appendices

**Methods S1.** Discussion on fossils used for calibrating divergence time analyses.

**Table S1.** Accession information for the taxa included in the chloroplast alignment.

**Table S2.** Accession information for the taxa included in the nuclear genomic and transcriptomic data set.

**Table S3.** Counts of bipartitions representing nodes A-H and conflicting bipartitions representing other subfamily relationships among 3,473 gene trees.

**Table S4.** Age intervals specified for the fossil calibration priors under different alternative priors.

**Table S5.** Node age estimates and priors (95% HPD intervals) of nodes A-H in the different analyses.

**Figure S1.** ML topology as inferred by RAxML from amino acid alignment of chloroplast genes under the LG4X model. Numbers on nodes indicate bootstrap percentages estimated from 1000 replicates.

**Figure S2.** Bayesian majority-rule consensus tree inferred with Phylobayes from amino acid alignment of chloroplast genes under the CATGTR model. Numbers on nodes indicate posterior probabilities (pp) from 9000 post-burn-in MCMC cycles.

**Figure S3.** ML topology as inferred by RAxML from nucleotide alignment of chloroplast genes under the GTR + G model. Numbers on nodes indicate bootstrap percentages estimated from 1000 replicates.

**Figure S4.** Bayesian majority-rule consensus tree inferred with Phylobayes from nucleotide alignment of chloroplast genes under the CATGTR model. Numbers on nodes indicate the posterior probabilities (pp) from 9000 post-burn-in MCMC cycles.

**Figure S5.** ML topology as inferred by RAxML from a concatenated alignment of 1,103 nuclear genes, under the LG4X model. Numbers on nodes indicate Internode Certainty All (ICA) values, as estimated from gene trees of the same 1,103 genes.

**Figure S6.** Bayesian gene jackknifing majority-rule consensus tree inferred with Phylobayes from a concatenated alignment of 1,103 nuclear genes. Numbers on nodes indicate posterior probabilities (pp), averaged over 500 posterior trees each, for 25 replicates (12,500 posterior trees in total).

**Figure S7.** Phylogeny estimated under the multi-species coalescent with ASTRAL. Support values indicated represent local posterior probability (blue rectangles) and quartet support (yellow rectangles).

**Figure S8.** Examples of homolog clusters with gene duplications in legumes that passed the bootstrap filter. Yellow stars behind nodes indicate locations of gene duplications, numbers on nodes indicate bootstrap support. The plotted gene trees are extracted from (A) cluster3675_1rr_1rr, showing a duplication subtending Detarioideae, (B) cluster1032_1rr_1rr, showing a duplication subtending Papilionoideae, (C) cluster1248_1rr_1rr and (D) cluster2941_1rr_1rr, both with a duplication subtending the legume family. Trees for (E) cluster51_7rr_1rr and (F) cluster544_1rr_1rr show evidence of more than one duplication, including one specific to Papilionoideae in the former.

**Figure S9.** Numbers of gene duplications mapped across the phylogeny. The topology used is the ML topology of the nuclear concatenated alignment of 1,103 genes, duplications were counted from 8,038 homolog clusters. Numbers above branches (with blue background) and below branches (with yellow background) represent numbers of duplications and numbers of homolog trees with duplications without or with a bootstrap filter of 50%, respectively.

**Figure S10.** Chronogram estimated under the UCLN clock model. Numbers behind nodes indicate 95% HPD intervals. Substitution rate is indicated by colored branches, as indicated by the color legend, in substitutions per site per million years. Fossil calibrations as listed in Table 1 are indicated by blue labeled circles.

**Figure S11.** Chronogram estimated under the UCLN clock model, with alternative prior 2. Numbers behind nodes indicate 95% HPD intervals. Substitution rate is indicated by colored branches, as indicated by the color legend, in substitutions per site per million years. Fossil calibrations as listed in Table 1 are indicated by blue labeled circles.

**Figure S12.** Chronogram estimated under the RLC model. Numbers behind nodes indicate 95% HPD intervals. Substitution rate is indicated by colored branches, as indicated by the color legend, in substitutions per site per million years. Fossil calibrations as listed in Table 1 are indicated by blue labeled circles.

**Figure S13.** Chronogram estimated under the FLC3 model. Numbers behind nodes indicate 95% HPD intervals. Clock partitions are indicated by colored branches. Fossil calibrations as listed in Table 1 are indicated by blue labeled circles.

**Figure S14.** Chronogram estimated under the FLC6 model. Numbers behind nodes indicate 95% HPD intervals. Cock partitions are indicated by colored branches. Fossil calibrations as listed in Table 1 are indicated by blue labeled circles.

**Figure S15.** Chronogram estimated under the FLC8 model. Numbers behind nodes indicate 95% HPD intervals. Clock partitions are indicated by colored branches. Fossil calibrations as listed in Table 1 are indicated by blue labeled circles.

**Figure S16.** Chronogram estimated under the FLC8 model, with alternative prior 1. Numbers behind nodes indicate 95% HPD intervals. Clock partitions are indicated by colored branches. Fossil calibrations as listed in Table 1 are indicated by blue labeled circles, with alternative calibrations as red circles.

**Figure S17.** Chronogram estimated under the STRC model. Numbers behind nodes indicate 95% HPD intervals. Fossil calibrations as listed in Table 1 are indicated by blue labeled circles.

**Figure S18.** Substitution rates as estimated in FLC8 analyses for the different clock partitions. Boxplots for each partition for (A) alternative prior 1 and (B) the “normal” prior setting. Colors correspond to the partitions as shown in Figures 5, S14, S15 and S18.

**Figure S19.** Root-to-tip lengths per taxon with partitions of fixed local clocks indicated. Pruned taxa with outlier root-to-tip lengths are indicated with an X, partitions are indicated with colors. (A) FLC3, (B) FLC6, (C) FLC8.

