## Supplementary Methods, Tables and Figures for "The Origin and Early Evolution of the Legumes are a Complex Paleopolyploid Phylogenomic Tangle closely associated with the Cretaceous-Paleogene (K-Pg) Boundary"

**Methods S1. Discussion on fossils used for calibrating divergence time analyses.**

*Non-legume Eudicot Fossils*

These were taken from Magallon et al. (2015) and are thoroughly discussed in the supplementary information of that article. The numbers listed in Table 1 are the same numbers as used in the Supplementary Information Methods 1 of Magallon et al. (2015). We have followed their fossil placements although our more limited taxon sampling means that some minimum ages are placed on deeper nodes. The only exception is the stem node of Fagales (calibration X14), which was here calibrated using the oldest fossil prior used by Xing et al. (2014). All minimum ages were updated to the latest version of the Geologic Time Scale (v. 4.0; Gradstein et al., 2012).

*Legume Fossils*

The selection of legume fossils used here for calibrating the divergence time estimation analyses differs from previous legume time tree studies (Lavin et al., 2005; Bruneau et al., 2008; Simon et al., 2009), both in the placement of fossils as well as in the minimum ages that some of these fossils represent. Calibrations Q2 and Z are used for the first time here. Calibrations A, D, F, G, I2, M2 and Q are labelled according to the schemes of Bruneau et al. (2008) and/or Simon et al. (2009), and differences from previous studies are discussed here. Other fossils used by Bruneau et al. (2008) and/or Simon et al. (2009) are not used here because of our sparser taxon sampling.

First, we did not fix the crown age of the family, which is critical as it is the most important node for which we want to estimate the age. The oldest definitive legume fossil, a fossil wood from the Early Paleocene of Patagonia (Brea et al., 2008), is used to set a minimum age on the stem node of the family at 63.5 Ma (calibration A, same node as in Bruneau et al. (2008) and Simon et al. (2009), but a new fossil and minimum age). This calibration is probably not very informative because of the long stem of the family, but it is included for completeness. The oldest crown group fossil, bipinnate leaves from the Late Paleocene of Colombia (Wing et al., 2009; Herrera et al., submitted), is placed on the stem node of Caesalpinioideae with a minimum age of 58 Ma (calibration Z), a new calibration that has not been used in previous studies. This calibration renders the calibration of the stem of Papilionoideae (which is sister to Caesalpinioideae), with fossil flowers of *Barnebyanthus buchananensis* from the Paleocene-Eocene boundary at 56 Ma (Crepet & Herendeen, 1992), redundant.

We find the interpretation of some Early and Middle Eocene fossils, that were used in previous studies to calibrate lineages within crown group Cercidoideae and Detarioideae (Bruneau et al., 2008; Simon et al., 2009) to be problematic. Bruneau et al. (2008: Table 3) already pointed out the large discrepancy in age estimates of Detarioideae between calibrated and non-calibrated analyses. Given that this subfamily has a very long stem lineage (Figs 2 & 3), placing Early to Middle Eocene fossils within the crown group would require very high inferred substitution rates along the stem lineage, while at the same time implying a relatively low substitution rate for the Detarioideae crown group lineages (see Results). Cercidoideae are also subtended by a long stem lineage, leading to similar, although less severe substitution rate discrepancies than in Detarioideae. We investigate and test this with molecular clock analyses with fixed local clocks, as described below. Here, we discuss the interpretation of these fossils as either stem or crown relatives and how we have calibrated lineages from subfamilies Cercidoideae and Detarioideae.

*Bauhinia*-like bilobed leaves from the Eocene of Tanzania (c. 46 Ma) (Jacobs & Herendeen, 2004) were used by Bruneau et al. (2008) and Simon et al. (2009) to calibrate the stem lineage of *Bauhinia* s.l.. This leaf type is highly characteristic for Cercidoideae and therefore the fossil is certainly representative of the subfamily. However, even though this type of leaf is not found in *Cercis*, which has been found to be sister to the rest of the genera in the subfamily (Bruneau et al., 2008; Wang et al., 2018), it may not provide a strong apomorphy for crown group Cercidoideae. Leaves in *Bauhinia* s.l. are variously bifoliolate, bilobed or entire, implying that entire leaves like those of *Cercis* have evolved multiple times independently, leading to homoplasy. This means that the bilobed leaves may have been present in the most recent common ancestor (MRCA) or stem relatives of Cercidoideae, and evolved to having an entire lamina in *Cercis.* If the Tanzanian fossils are a possible stem-relative of Cercidoideae, we consider the oldest definitive crown group fossil evidence to be the recently described *Cercis* fossil leaves and fruits from the Late Eocene of Oregon (Jia & Manchester, 2014), at c. 36 Ma (calibration C, a slightly older minimum age than used by Bruneau et al. (2008) and Simon et al. (2009)).

Bifoliolate leaves from the same fossil site in Tanzania as the *Bauhinia* fossil were ascribed to *Aphanocalyx* (Detarioideae) (Herendeen & Jacobs, 2000) based on distinctive venation patterns, after comparing the leaves to all extant legume genera with bifoliolate leaves. The fossil was used to calibrate the stem lineage of that genus by Bruneau et al. (2008) and Simon et al. (2009). The genus is deeply nested within Detarioideae, also meaning that the difference between age estimates from calibrated and uncalibrated analyses is large (46.0 vs 4.4 Ma; Bruneau et al., 2008: Table 3). While venation patterns can be diagnostic in many cases, they are often variable even within modern genera and likely to be homoplasious. Therefore, these fossils might also represent an extinct lineage, possibly a stem relative of Detarioideae, that had evolved similar leaf morphology to extant *Aphanocalyx.* Moreover, the author of the most recent taxonomic account of *Aphanocalyx* (Wieringa, 1999), Jan Wieringa, does not accept this fossil as belonging to the genus. It also does not fit with the morphology-based phylogeny of *Aphanocalyx* which shows that bifoliolate leaves have evolved recently and are derived within *Aphanocalyx* (Wieringa, 1999). In general, leaflet numbers are highly variable across Detarioideae, so relatives of fossils should not be sought only among other bifoliolate taxa.

Further evidence of Detarioideae from the Eocene is found at two localities within the Claiborne Formation in western Tennessee, USA. Fruits and leaflets from those sites are ascribed to the genus *Crudia* (Herendeen & Dilcher, 1990). As for the *Aphanocalyx* fossil, the affinities of the fossils were carefully evaluated before concluding that they are related to *Crudia.* Bruneau et al. (2008) and Simon et al. (2009) used this fossil to calibrate the stem of *Crudia* at 45 Ma, but as for the *Aphanocalyx* fossil age, an uncalibrated analysis finds a far younger age (6.9 Ma; Bruneau et al., 2008: Table 3). It is possible that in this case, an extinct detarioid lineage may have evolved morphological features similar to extant *Crudia* species independently. The raised venation on the fruit valves and twisted petiolules that most strongly resemble *Crudia*, for example, are both homoplasious across Detarioideae.

Fossil wood, flowers and amber of *Aulacoxylon sparnacense*, which has previously been interpreted as related to the extant genus *Daniellia* (Detarioideae), from the Early Eocene of the Paris basin (De Franceschi & De Ploëg, 2003), provide the most convincing evidence of fossils representing Early to Middle Eocene crown group members of Detarioideae. The fossil wood has vestured pits and resin canals, like modern resin-producing

Detarioideae and the amber deposits are chemically similar to the Dominican ambers. Bruneau et al. (2008) considered the wood and flowers similar to *Daniellia*, but suggested they could also belong to a different genus of resin-producing Detarieae. However, it is also possible that resin-production was already present in stem-relatives of Detarioideae. This is quite likely given that this trait is homoplasious across the resin-producing clade, having apparently been independently gained or lost several times, with only about half of the extant genera in the clade producing resin (Fougère-Danezan et al., 2007). If the production of resin evolved in the ancestral lineage of Detarioideae it would not require many more losses to account for the absence of the trait in the other lineages of the subfamily, because the resin-producing clade branches deeply within Detarioideae and the basal relationships of the subfamily are poorly resolved and understood (Bruneau et al., 2008; de la Estrella et al., 2018). Furthermore, the large majority of genera in the subfamily are confined to the large clade of Amherstieae, so perhaps only a single additional loss of the trait in the lineage leading to this clade could have produced this homoplasious pattern. This makes it possible that the Paris basin fossils belong to an extinct genus belonging to the stem group of Detarioideae. Therefore, the *Aulacoxylon* fossils can be used either to calibrate the stem node of the resin producing clade (calibration $G^g$, as was done by Bruneau et al, (2008) and Simon et al., (2009) or the stem node of Detarioideae (calibration G), with a minimum age of 53 Ma .

For the disputed age of Dominican amber (Iturralde-Vinent & MacPhee, 1996), an intermediate age of 24 Ma was chosen by Bruneau et al. (2008), which was followed by Simon et al. (2009), but it is preferable to not consider an intermediate age as a valid minimum, but rather to use the minimum age that was estimated for Mexican amber that includes flowers of *Hymenaea mexicana*, the extinct species that presumably produced the amber (Poinar & Brown, 2002), and we calibrate the Detarieae s.s. stem node with a minimum age of 22.5 Ma (calibration F, a more inclusive node than in Bruneau et al, (2008) and Simon et al., (2009), and a different fossil age).

The calibration of the stem group of Styphnolobium and Cladrastis (calibration I2) is the same as used in Bruneau et al. (2008) and Simon et al. (2009), but the minimum age was updated to 37.8 Ma according to the latest version of the Geologic Time Scale (v. 4.0; Gradstein et al., 2012), representing the end of the Middle Eocene (end of the Bartonian). Calibration M2 is the same as used in Simon et al., (2009) but since *Robinia* itself is not sampled here, we place it on the stem node of the robinioid clade (represented here by *Lotus japonicus*) and update the minimum age to the Eocene-Oligocene boundary at 33.9 Ma.

Bruneau et al. (2008) and Simon et al. (2009) also set the ages of several fossil calibrations at the midpoint of the Eocene, at 45 Ma. This led to a bias that was observable in an LTT plot of legumes (Koenen et al., 2013), and here we prefer to use the minimum boundary ages for these fossils. Although most of these calibrations are not used in our analyses due to sparser taxon sampling, we use one of these fossils, *Acacia*-like polyads, to calibrate the minimum stem age of the clade including all *Acacia* s.l. segregates at 33.9 Ma, the Eocene-Oligocene boundary (calibration Q, same node but younger age than Bruneau et al, (2008) and Simon et al., (2009)). Finally, we add calibration Q2, based on Australian Oligocene polyads with pseudocolpi (Miller et al., 2013), which suggest affinity with *Acacia* s.s., and we calibrate the stem node of that genus with a minimum age of 23 Ma, the Oligocene-Miocene boundary.

*Alternative Calibrations for Detarioideae and Cercidoideae*

In the analysis with “alternative prior 1”, calibration C was replaced with a minimum age of 46 Ma on the stem of *Bauhinia*, based on fossil leaves from Tanzania (Herendeen & Jacobs, 2000; discussed above). Calibration G[g] was applied on the stem of the resin-producing clade (i.e. the crown node of Detarioideae) instead of on the stem node of Detarioideae. Calibration H[g] is taken from Bruneau et al. (2008) and Simon et al. (2009), and is added in the alternative analysis to specify a minimum age of 46 Ma on the stem of *Anthonotha*, based on fossil leaves assigned to the closely related genus *Aphanocalyx* from Tanzania (Herendeen & Jacobs, 2000; but see above).

**Table S1. Accession information for the taxa included in the chloroplast alignment.**

| Taxon | Herbarium voucher | Genbank accession number | Comments |
|---|---|---|---|
| *Abarema jupunba* | M.F. Simon 1600 (CEN) | XXXXXXXXXX | **Newly sequenced** |
| *Acacia koa* | | see Table S1. | **Transcriptome** |
| *Acacia ligulata* | | LN555649.2 | |
| *Acrocarpus fraxinifolius* | | | **Transcriptome, available at https://ics.hutton.ac.uk/tropiTree/** |
| *Adenanthera pavonina* | Ambriansyah & Arifin AA295 (K) | XXXXXXXXXX | **Newly sequenced** |
| *Afzelia africana* | S.L.A. Donkpegan 27 (BRLU) | KX673213 | |
| *Afzelia bipindensis* | S.L.A. Donkpegan 626 (BRLU) | XXXXXXXXXX | **Newly sequenced** |
| *Ajuga reptans* | | KF709391 | |
| *Albizia adianthifolia* | J.J. Wieringa 6278 (WAG) | XXXXXXXXXX | **Newly sequenced** |
| *Albizia julibrissin* | E. Koenen 601 (Z) | XXXXXXXXXX | **Newly sequenced; Transcriptome** |
| *Anthonotha fragrans* | | see Table S1. | **Newly sequenced; Transcriptome** |
| *Apios americana* | | KF856618 | |
| *Arabidopsis thaliana* | | AP000423 | |
| *Arachis hypogaea* | | KJ468094 | |
| *Arachis ipaensis* | | GBIW00000000 | **Transcriptome** |
| *Aralia undulata* | R. Li 551 (KUN) | KC456163 | |
| *Archidendron lucidum* | Wang & Lin 2534 (L) | XXXXXXXXXX | **Newly sequenced** |
| *Astragalus membranaceus* | | | **Transcriptome, available at http://dx.doi.org/10.5061/dryad.ff1tq** |
| *Astragalus propinquus* | | | **Transcriptome, OneKP: MYMP, available at** |

| | | | |
|---|---|---|---|
| | | | **http://www.onekp.com/public_data.html** |
| *Azadirachta indica* | | KF986530 | |
| *Bauhinia tomentosa* | | see Table S1. | **Transcriptome, available at http://dx.doi.org/10.5061/dryad.ff1tq** |
| *Bituminaria bituminosa* | | see Table S1. | **Transcriptome, OneKP: TVSH, available at http://www.onekp.com/public_data.html** |
| *Bulnesia arborea* | M.J. Moore 334 (FLAS) | EU002159, EU002172, EU002205, EU002275, EU002299, EU002388, EU002478, GQ998005-GQ998073, HQ664597 | |
| *Buxus microphylla* | | EF380351 | |
| *Calliandra hygrophila* | L.P. Queiroz 15542 (HUEFS) | XXXXXXXXXX | **Newly sequenced** |
| *Carica papaya* | | EU431223 | |
| *Ceratonia siliqua* | | KJ468096 | |
| *Cercis canadensis* | | KF856619 | |
| *Chamaecrista fasciculata* | | see Table S1. | **Transcriptome, available at http://dx.doi.org/10.5061/dryad.ff1tq** |
| *Chidlowia sanguinea* | J.J. Wieringa 4338 (WAG) | XXXXXXXXXX | **Newly sequenced** |
| *Chrysobalanus icaco* | | KJ414480 | |
| *Cicer arietinum* | | EU835853 | |
| *Citrus sinensis* | | DQ864733 | |
| *Cladrastis lutea* | | | **Transcriptome, available at http://dx.doi.org/10.5061/dryad.ff1tq** |
| *Codariocalyx motorius* | | | **Transcriptome, available at http://dx.doi.org/10.5061/dryad.ff1tq** |
| *Coffea arabica* | | EF044213 | |

| | | | |
|---|---|---|---|
| *Cojoba arborea* | M.F. Simon 1545 (CEN) | XXXXXXXXXX | **Newly sequenced** |
| *Colvillea racemosa* | Kew living collection 1993-224 (K) | XXXXXXXXXX | **Newly sequenced** |
| *Copaifera officinalis* | | | **Transcriptome, available at http://dx.doi.org/10.5061/dryad.ff1tq** |
| *Cucumis sativus* | | AJ970307 | |
| *Daucus carota* | | DQ898156 | |
| *Desmanthus illlinoiensis* | | | **Transcriptome, available at http://dx.doi.org/10.5061/dryad.ff1tq** |
| *Dialium guineense* | T. van Andel 4184 (WAG) | XXXXXXXXXX | **Newly sequenced** |
| *Dichrostachys cinerea* | O. Maurin 256 (JRAU) | XXXXXXXXXX | **Newly sequenced** |
| *Dimorphandra macrostachya* | J.R. Igançi 877 (RB) | XXXXXXXXXX | **Newly sequenced** |
| *Diptychandra aurantiaca* | J.R.I. Wood 26513 (K) | XXXXXXXXXX | **Newly sequenced** |
| *Distemonanthus benthamianus* | G. Dauby 728 (BRLU) | XXXXXXXXXX | **Newly sequenced** |
| *Duparquetia orchidacea* | J.J. Wieringa 7805 (L) | XXXXXXXXXX | **Newly sequenced** |
| *Entada abyssinica* | MSB 0133199 (K) | XXXXXXXXXX | **Newly sequenced; Transcriptome** |
| *Entada rheedei* | E. Koenen 496 (Z) | XXXXXXXXXX | **Newly sequenced** |
| *Erythrophleum ivorense* | J.J. Wieringa 5487 (WAG) | XXXXXXXXXX | **Newly sequenced** |
| *Erythrostemon gilliesii* | R. Steeves 852 (MT) | XXXXXXXXXX | **Newly sequenced** |
| *Eucalyptus grandis* | | HM347959 | |
| *Euonymus americanus* | W. Judd 8071 (FLAS) | EU002160, EU002170, EU002193, EU002277, EU002321, EU002409, EU002500, GQ998147-GQ998219, HQ664608 | |

| | | | |
|---|---|---|---|
| *Fagopyrum esculentum* | | EU254477 | |
| *Faidherbia albida* | O. Maurin 3495 (JRAU) | XXXXXXXXXX | **Newly sequenced** |
| *Garcinia mangostana* | | HQ331601, HQ331906, HQ332057, HQ848709, JX661816, JX661859, JX661902, JX661944, JX661980, JX662020, JX662065, JX662109, JX662151, JX662196, JX662237, JX662279, JX662320, JX662359, JX662399, JX662434, JX662467, JX662502, JX662543, JX662580, JX662622, JX662666, JX662710, JX662752, JX662799, JX662841, JX662880, JX662914, JX662955, JX662996, JX663032, JX663071, JX663104, JX663149, JX663196, JX663237, JX663280, JX663322, JX663365, JX663410, JX663583, JX663630, JX663677, JX663721, JX663763, JX663804, JX663841, JX663874, JX663915, JX663962, JX664006, JX664049, JX664091, JX664127, JX664165, JX664209, JX664252, JX664297, JX664341, JX664385, | |

| | | | |
|---|---|---|---|
| | | JX664458, JX664495, JX664535, JX664580, JX664623, JX664659, JX664694, JX664726, JX664771, JX664812, JX664852, JX664895, JX664939, JX665004, KF783277, U92876, U92877, U92878 | |
| *Gleditsia sinensis* | | | **Transcriptome, OneKP: VHZV, available at http://www.onekp.com/public_data.html** |
| *Gleditsia triacanthos* | | | **Transcriptome, available at http://dx.doi.org/10.5061/dryad.ff1tq** |
| *Gliricidia sepium* | | | **Transcriptome, available at https://ics.hutton.ac.uk/tropiTree/** |
| *Glycine canescens* | | KC893635 | |
| *Glycine max* | | DQ317523 | |
| *Glycyrrhiza glabra* | | KF201590 | |
| *Glycyrrhiza lepidota* | | | **Transcriptome, available at http://dx.doi.org/10.5061/dryad.ff1tq** |
| *Gompholobium polymorphum* | | | **Transcriptome, OneKP: VLNB, available at http://www.onekp.com/public_data.html** |
| *Gossypium hirsutum* | | DQ345959 | |
| *Guibourtia ehie* | F. Tosso 272 (BRLU) | XXXXXXXXXX | **Newly sequenced** |
| *Guibourtia tessmannii* | | XXXXXXXXXX | **Newly sequenced** |
| *Guilfoylia monostylis* | P.I. Forster 28103 (Z) | XXXXXXXXXX | **Newly sequenced** |
| *Gymnocladus dioicus* | | | **Transcriptome, available at http://dx.doi.org/10.5061/dryad.ff1tq** |
| *Haematoxylum brasiletto* | | KJ468097 | |
| *Helianthus annuus* | | DQ383815 | |

| | | | |
|---|---|---|---|
| *Hymenostegia brachyura* | Zenker 4481 (WAG) | XXXXXXXXXX | **Newly sequenced** |
| *Hymenostegia felicis* | Jacques-Félix 5129 (WAG) | XXXXXXXXXX | **Newly sequenced** |
| *Indigofera tinctoria* | | KJ468098 | |
| *Inga leiocalycina* | T.D. Pennington 13822 (K) | KT428296 | |
| *Inga spectabilis* | T.D. Pennington 15061 (K) | XXXXXXXXXX | **Newly sequenced** |
| *Intsia bijuga* | | KX673214 | |
| *Lathyrus graminifolius* | | KJ806193 | |
| *Lathyrus sativus* | | HM029371 | |
| *Lens culinaris* | | KF186232 | |
| *Leucaena trichandra* | | KT428297 | |
| *Libidibia coriaria* | | KJ468095 | |
| *Lotus japonicus* | | AP002983 | |
| *Lupinus luteus* | | KC695666 | |
| *Lupinus polyphyllus* | | | **Transcriptome, OneKP: CMFF, available at http://www.onekp.com/public_data.html** |
| *Macadamia integrifolia* | | KF862711 | |
| *Manihot esculenta* | | EU117376 | |
| *Medicago hybrida* | | KJ850240 | |
| *Medicago truncatula* | | AC093544 | |
| *Microlobius foetidus* | C.E. Hughes 1219 (FHO) | XXXXXXXXXX | **Newly sequenced; Transcriptome** |
| *Millettia pinnata* | | JN673818 | |
| *Mimosa tenuiflora* | L.P. Queiroz 15498 (HUEFS) | XXXXXXXXXX | **Newly sequenced** |
| *Morus indica* | | DQ226511 | |
| *Nelumbo nucifera* | | JQ336993 | |
| *Nerium oleander* | W. Judd 8076 (FLAS) | KJ953907 | |

| | | | |
|---|---|---|---|
| *Newtonia hildebrandtii* | O. Maurin 2457 (JRAU) | XXXXXXXXXX | **Newly sequenced** |
| *Oenothera biennis* | | EU262889 | |
| *Olea europaea* | | GU228899 | |
| *Oxalis latifolia* | M.J. Moore 316 (FLAS) | EU002165, EU002186, EU002248, EU002282, EU002350, EU002438, EU002528, GQ998511-GQ998580, HQ664602, KF783277, U92876, U92877, U92878 | |
| *Pachyelasma tessmannii* | J.J. Wieringa 5229 (WAG) | XXXXXXXXXX | **Newly sequenced** |
| *Pachyrhizus erosus* | | KJ468100 | |
| *Paeonia obovata* | | KJ206533 | |
| *Parkia panurensis* | J.R. Igançi 842 (RB) | XXXXXXXXXX | **Newly sequenced** |
| *Pelargonium alternans* | | KF240617 | |
| *Peltophorum africanum* | Koenen 601 (Z) | XXXXXXXXXX | **Newly sequenced** |
| *Pentaclethra macrophylla* | Galeuchet & Balthazar 10 (Z) | XXXXXXXXXX | **Newly sequenced** |
| *Phaseolus vulgaris* | | DQ886273 | |
| *Piptadeniastrum africanum* | E. Koenen 152 (WAG) | XXXXXXXXXX | **Newly sequenced** |
| *Pisum sativum* | | HM029370 | |
| *Pithecellobium dulce* | B. Marazzi 309 (?) | XXXXXXXXXX | **Newly sequenced** |
| *Poeppigia procera* | Hernández 558 (Z) | XXXXXXXXXX | **Newly sequenced** |
| *Polygala lutea* | | EF489041 | |
| *Populus trichocarpa* | | KF753634 | |
| *Primula poissonii* | | KF753634 | |

| | | | |
|---|---|---|---|
| *Prioria balsamifera* | | XXXXXXXXXX | **Newly sequenced; Transcriptome** |
| *Prosopis alba* | | see Table S1. | **Transcriptome** |
| *Prosopis glandulosa* | | KJ468101 | |
| *Prunus persica* | | HQ336405 | |
| *Quercus rubra* | | JX970937 | |
| *Quillaja saponaria* | | | **Transcriptome, available at http://dx.doi.org/10.5061/dryad.ff1tq** |
| *Ranunculus macranthus* | | DQ359689 | |
| *Robinia pseudoacacia* | | KJ468102 | |
| *Samanea saman* | C.E. Hughes 421 (FHO) | XXXXXXXXXX | **Newly sequenced** |
| *Sapindus mukorossi* | | KM454982 | |
| *Saraca indica* | Kew living collection 2011-1421 (K) | XXXXXXXXXX | **Newly sequenced** |
| *Schotia brachypetala* | R. Steeves 846 (MT) | XXXXXXXXXX | **Newly sequenced** |
| *Sedum sarmentosum* | | JX427551 | |
| *Senegalia ataxacantha* | C. Jongkind 10603 (WAG) | XXXXXXXXXX | **Newly sequenced** |
| *Senna hebecarpa* | | | **Transcriptome, available at http://dx.doi.org/10.5061/dryad.ff1tq** |
| *Senna siamea* | | | **Transcriptome, available at https://ics.hutton.ac.uk/tropiTree/** |
| *Sesbania macrantha* | | | **Transcriptome, available at https://ics.hutton.ac.uk/tropiTree/** |
| *Sesbania sesban* | | | **Transcriptome, available at https://ics.hutton.ac.uk/tropiTree/** |
| *Silene latifolia* | | JF715055 | |
| *Solanum lycopersicum* | | KP331414 | |
| *Styphnolobium japonicum* | | see Table S1. | **Transcriptome, available at https://www.hindawi.com/journals/bmri/2014/75** |

| | | | |
|---|---|---|---|
| | | | **0961/sup/** |
| *Swartzia emarginata* | M.P. Morim 576 (RB) | XXXXXXXXXX | **Newly sequenced** |
| *Tachigali odoratissima* | M.P. Morim 562 (RB) | XXXXXXXXXX | **Newly sequenced** |
| *Tamarindus indica* | | KJ468103 | |
| *Theobroma cacao* | | HQ244500 | |
| *Tipuana tipu* | | | **Transcriptome, available at https://ics.hutton.ac.uk/tropiTree/** |
| *Trachelium caeruleum* | | EU090187 | |
| *Trifolium aureum* | | KC894708 | |
| *Trifolium repens* | | KC894706 | |
| *Trochodendron aralioides* | | KC608753 | |
| *Vaccinium macrocarpon* | | JQ757046 | |
| *Vachellia tortilis* | E. Koenen 603 (Z) | XXXXXXXXXX | **Newly sequenced** |
| *Vicia faba* | | KF042344 | |
| *Vicia sativa* | | KJ850242 | |
| *Vigna radiata* | | GQ893027 | |
| *Vigna unguiculata* | | JQ755301 | |
| *Vitis vinifera* | | DQ424856 | |
| *Wisteria floribunda* | | | **Transcriptome, OneKP: RMWJ, available at http://www.onekp.com/public_data.html** |
| *Xanthocercis zambesiaca* | | | **Transcriptome, available at http://dx.doi.org/10.5061/dryad.ff1tq** |
| *Xanthophyllum eurhynchum* | P. Herendeen H.416 (?) | XXXXXXXXXX | **Newly sequenced** |
| *Xylia hoffmannii* | E. Koenen 402 (Z) | XXXXXXXXXX | **Newly sequenced** |
| *Zenia insignis* | Averyanov et al. 5748 (?) | XXXXXXXXXX | **Newly sequenced** |

**Table S2. Accession information for the taxa included in the nuclear genomic and transcriptomic data set.**

| Taxon | Source | Citation |
|---|---|---|
| *Acacia koa* | Genbank BioProject: PRJNA268386 | Ishihara et al. |
| *Acrocarpus fraxinifolius* | TropiTree: https://ics.hutton.ac.uk/tropiTree/ | Russel et al. 2014 |
| *Afzelia bella* | Genbank BioProject: XXXXXXXXXX | **Newly sequenced** |
| *Albizia julibrissin* | Genbank BioProject: XXXXXXXXXX | **Newly sequenced** |
| *Alnus serrulata* | Dryad: http://dx.doi.org/10.5061/dryad.ff1tq | Cannon et al. 2015 |
| *Amaranthus hypochondriacus* | Phytozome v11: v1.0 | Clouse et al. 2016 |
| *Anthonotha fragrans* | Genbank BioProject: XXXXXXXXXX | **Newly sequenced** |
| *Apios americana* | Dryad: http://dx.doi.org/10.5061/dryad.ff1tq | Cannon et al. 2015 |
| *Aquilegia coerulea* | Phytozome v11: v3.1 | Filiault et al. 2018 |
| *Arabidopsis thaliana* | Phytozome v11: TAIR10 | Lamesch et al. 2012 |
| *Arachis ipaensis* | Peanutbase.org: K30076.a1.M1 | Bertioli et al. 2016 |
| *Astragalus membranaceus* | Dryad: http://dx.doi.org/10.5061/dryad.ff1tq | Cannon et al. 2015 |
| *Bauhinia tomentosa* | Dryad: http://dx.doi.org/10.5061/dryad.ff1tq | Cannon et al. 2015 |
| *Bituminaria bituminosa* | OneKP: TVSH | Wicket et al. 2014 |
| *Cajanus cajan* | http://gigadb.org/dataset/100028 | Varshney et al. 2012 |
| *Cannabis sativa* | Genbank BioProject: PRJNA74271 | van Bakel et al. 2011 |
| *Carica papaya* | Phytozome v11: ASGPBv4.0 | Ming et al. 2008 |
| *Castanea mollissima* | https://www.hardwoodgenomics.org/Genome-assembly/1962958 | *Not available* |
| *Cercis canadensis* | Dryad: http://dx.doi.org/10.5061/dryad.ff1tq | Cannon et al. 2015 |
| *Chamaecrista fasciculata* | Dryad: http://dx.doi.org/10.5061/dryad.ff1tq | Cannon et al. 2015 |
| *Cicer arietinum* | http://gigadb.org/dataset/100076 | Varshney et al. 2013 |
| *Citrus sinensis* | Phytozome v11: v1.1 | Wu et al. 2014 |

| | | |
|---|---|---|
| *Cladrastis lutea* | Dryad: http://dx.doi.org/10.5061/dryad.ff1tq | Cannon et al. 2015 |
| *Codariocalyx motorius* | Dryad: http://dx.doi.org/10.5061/dryad.ff1tq | Cannon et al. 2015 |
| *Copaifera officinalis* | Dryad: http://dx.doi.org/10.5061/dryad.ff1tq | Cannon et al. 2015 |
| *Cucumis sativus* | Phytozome v11: v1.0 | *Not available* |
| *Desmanthus illinoensis* | Dryad: http://dx.doi.org/10.5061/dryad.ff1tq | Cannon et al. 2015 |
| *Elaeocarpus photiniifolia* | Genbank BioProject: PRJDA67329 | Sugai et al. 2012 |
| *Entada abyssinica* | Genbank BioProject: XXXXXXXXXX | **Newly sequenced** |
| *Eucalyptus grandis* | Phytozome v11: v2.0 | Bartholomé et al. 2015 |
| *Fragaria vesca* | Phytozome v11: v1.1 | Shulaev et al. 2011 |
| *Gleditsia triacanthos* | Dryad: http://dx.doi.org/10.5061/dryad.ff1tq | Cannon et al. 2015 |
| *Glycine max* | Phytozome v11: Wm82.a2.v1 | Schmutz et al. 2010 |
| *Glycyrrhiza lepidota* | Dryad: http://dx.doi.org/10.5061/dryad.ff1tq | Cannon et al. 2015 |
| *Gossypium raimondii* | Phytozome v11: v2.1 | Paterson et al. 2012 |
| *Gymnocladus dioicus* | Dryad: http://dx.doi.org/10.5061/dryad.ff1tq | Cannon et al. 2015 |
| *Inga spectabilis* | https://doi.org/10.5061/dryad.r9c12 | Nicholls et al. 2015 |
| *Juglans regia* | https://www.hardwoodgenomics.org/Genome-assembly/2209485 | Martínez-García et al. 2016 |
| *Lactuca sativa* | Genbank BioProject: PRJNA65477 | *Not available* |
| *Lathyrus sativus* | Dryad: http://dx.doi.org/10.5061/dryad.ff1tq | Cannon et al. 2015 |
| *Lens culinaris* | Genbank BioProject: PRJNA65667 | Kaur et al. 2011 |
| *Linum usitatissimum* | Phytozome v11: v1.0 | Wang et al. 2012 |
| *Lotus japonicus* | http://www.plantgdb.org/LjGDB/ | Sato et al. 2008 |
| *Lupinus angustifolius* | Dryad: http://dx.doi.org/10.5061/dryad.ff1tq | Cannon et al. 2015 |
| *Lupinus polyphyllus* | Dryad: http://dx.doi.org/10.5061/dryad.ff1tq | Cannon et al. 2015 |
| *Manihot esculenta* | Phytozome v11: v6.1 | Bredeson et al. 2016 |
| *Medicago truncatula* | Phytozome v11: Mt4.0v1 | Young et al. 2011 |

| | | |
|---|---|---|
| *Microlobius foetidus* | Genbank BioProject: XXXXXXXXXX | **Newly sequenced** |
| *Mimulus guttatus* | Phytozome v11: v2.0 | Hellsten et al. 2013 |
| *Morus notabilis* | Genbank BioProject: PRJNA202089 (assembly version ASM41409v2) | He et al. 2013 |
| *Nelumbo nucifera* | Genbank BioProject: PRJNA264089 (assembly version 1.1) | Ming et al. 2013 |
| *Paeonia lactiflora* | Genbank BioProject: PRJNA245064 | Zhang et al. 2015 |
| *Panax ginseng* | Genbank BioProject: PRJNA173906 | Li et al. 2013 |
| *Papaver somniferum* | Dryad: http://dx.doi.org/10.5061/dryad.ff1tq | Cannon et al. 2015 |
| *Phaseolus vulgaris* | Phytozome v11: v1.0 | Schmutz et al. 2014 |
| *Pisum sativum* | Genbank BioProject: PRJNA211622 | Duarte et al. 2014 |
| *Polygala lutea* | Dryad: http://dx.doi.org/10.5061/dryad.ff1tq | Cannon et al. 2015 |
| *Populus trichocarpa* | Phytozome v11: v3.0 | Tuskan et al. 2006 |
| *Primula veris* | https://doi.org/10.5061/dryad.2s200 | Nowak et al. 2015 |
| *Prioria balsamifera* | Genbank BioProject: XXXXXXXXXX | **Newly sequenced** |
| *Prosopis alba* | Genbank BioProject: PRJNA218545 | Torales et al. 2013 |
| *Prunus persica* | Phytozome v11: v2.1 | International Peach Genome Initiative et al., 2013 |
| *Punica granatum* | Genbank BioProject: PRJNA231033 | Ophir et al. 2014 |
| *Quillaja saponaria* | Dryad: http://dx.doi.org/10.5061/dryad.ff1tq | Cannon et al. 2015 |
| *Salix purpurea* | Phytozome v11: v1.0 | Zhou et al. 2018 |
| *Senna hebecarpa* | Dryad: http://dx.doi.org/10.5061/dryad.ff1tq | Cannon et al. 2015 |
| *Solanum tuberosum* | Phytozome v11: v3.4 | Sharma et al. 2013 |
| *Styphnolobium japonicum* | https://www.hindawi.com/journals/bmri/2014/750961/sup/ | Zhu et al. 2014 |
| *Theobroma cacao* | Phytozome v11: v1.1 | Motamayor et al. 2013 |
| *Trifolium pratense* | Genbank BioProject: PRJNA219226 | Yates et al. 2014 |
| *Tripterygium wilfordii* | Genbank BioProject: PRJNA218574 | *Not available* |

| | | |
|---|---|---|
| *Vicia faba* | Genbank BioProject: PRJNA81211 | Kaur et al. 2012 |
| *Vigna radiata* | ftp://plantgenomics.snu.ac.kr/mungbean_data/ | Kang et al. 2014 |
| *Vitis vinifera* | Phytozome v11: Genoscope.12X | Jaillon et al. 2007 |
| *Xanthocercis zambesiaca* | Dryad: http://dx.doi.org/10.5061/dryad.ff1tq | Cannon et al. 2015 |
| *Zenia insignis* | Genbank BioProject: PRJNA285444 | *Not available* |

Phytozome is available at https://phytozome.jgi.doe.gov
OneKP data is available at http://www.onekp.com/public_data.html

**Table S3. Counts of bipartitions representing nodes A-H and conflicting bipartitions representing other subfamily relationships among 3,473 gene trees.**

| Clade | ML | >50% bootstrap support | >80% bootstrap support |
|---|---|---|---|
| *bipartitions of best supported topology* | | | |
| **Leguminosae (node A)** | 2669 | 2254 | 1660 |
| **Cerc + Detar (node B)** | 744 | 325 | 48 |
| **Cercidoideae (node C)** | 1815 | 1705 | 1394 |
| **Detarioideae (node D)** | 3041 | 2918 | 2585 |
| **Pap + Caes + Dial (node E)** | 794 | 360 | 91 |
| **Pap + Caes (node F)** | 599 | 231 | 42 |
| **Caesalpinioideae (node G)** | 2114 | 1712 | 1151 |
| **Papilionoideae (node H)** | 2456 | 1957 | 1248 |
| *conflicting bipartitions* | | | |
| **Pap + Caes + Dial + Detar** | 625 | 258 | 34 |
| **Pap + Caes + Dial + Cerc** | 546 | 194 | 20 |
| **Caes + Dial** | 448 | 132 | 16 |
| **Pap + Dial** | 446 | 133 | 20 |
| **Dial + Detar** | 307 | 96 | 4 |
| **Dial + Cerc** | 295 | 93 | 7 |
| **Caes + Dial + Cerc + Detar** | 247 | 47 | 4 |
| **Pap + Caes + Cerc + Detar** | 234 | 46 | 2 |
| **Dial + Cerc + Detar** | 202 | 21 | 1 |
| **Caes + Detar** | 200 | 44 | 3 |
| **Pap + Dial + Cerc + Detar** | 196 | 29 | 4 |
| **Pap + Detar** | 189 | 41 | 2 |
| **Pap + Cerc** | 173 | 30 | 2 |
| **Caes + Cerc** | 163 | 37 | 2 |
| **Pap + Caes + Detar** | 153 | 27 | 1 |
| **Caes + Dial + Detar** | 134 | 11 | 0 |
| **Pap + Caes + Cerc** | 132 | 15 | 0 |
| **Caes + Cerc + Detar** | 127 | 14 | 0 |
| **Pap + Dial + Detar** | 122 | 12 | 1 |
| **Caes + Dial + Cerc** | 121 | 16 | 0 |
| **Pap + Cerc + Detar** | 115 | 9 | 0 |
| **Pap + Dial + Cerc** | 110 | 12 | 0 |

**Table S4. Age intervals specified for the fossil calibration priors under different alternative priors.**

| Calibration | Definition | MRCA taxon 1 | MRCA taxon 2 | Prior | Alternative prior 1 | Alternative prior 2 |
|---|---|---|---|---|---|---|
| *eudicots* | | | | | | |
| *26* | CG eudicots | *Aquilegia coerulea* | *Medicago truncatula* | normal (mean 126.0, stdev 1.0) | normal (mean 126.0, stdev 1.0) | normal (mean 126.0, stdev 1.0) |
| 27 | CG Ranunculales | *Aquilegia coerulea* | *Papaver somniferum* | uniform (min 113.0, max 126.0) | uniform (min 113.0, max 126.0) | uniform (min 113.0, max 126.0) |
| 38 | CG Pentapetalae | *Nelumbo nucifera* | *Medicago truncatula* | uniform (min 100.0, max 126.0) | uniform (min 100.0, max 126.0) | uniform (min 100.0, max 126.0) |
| 48 | SG Ericales | *Primula veris* | *Solanum tuberosum* | uniform (min 89.8, max 126.0) | uniform (min 89.8, max 126.0) | uniform (min 89.8, max 113.0) |
| 94 | SG Myrtaceae | *Eucalyptus grandis* | *Punica granatum* | uniform (min 83.6, max 126.0) | uniform (min 83.6, max 126.0) | uniform (min 83.6, max 100.0) |
| 105 | SG Brassicales | *Carica papaya* | *Theobroma cacao* | uniform (min 89.8, max 126.0) | uniform (min 89.8, max 126.0) | uniform (min 89.8, max 100.0) |
| 112 | CG Rosaceae | *Fragaria vesca* | *Prunus persica* | uniform (min 49.4, max 126.0) | uniform (min 49.4, max 126.0) | uniform (min 49.4, max 66.0) |
| 116 | SG Cannabaceae | *Cannabis sativa* | *Morus notabilis* | uniform (min 66.0, max 126.0) | uniform (min 66.0, max 126.0) | uniform (min 66.0, max 83.6) |
| 122 | SG Juglandaceae | *Alnus serrulata* | *Juglans regia* | uniform (min 64.4, max 126.0) | uniform (min 64.4, max 126.0) | uniform (min 64.4, max 83.6) |
| 133 | SG Populus | *Populus trichocarpa* | *Salix purpurea* | uniform (min 37.8, max 126.0) | uniform (min 37.8, max 126.0) | uniform (min 37.8, max 56.0) |
| X14 | SG Fagales | *Alnus serrulata* | *Medicago truncatula* | uniform (min 83.6, max 126.0) | uniform (min 83.6, max 126.0) | uniform (min 83.6, max 126.0) |
| *legumes* | | | | | | |
| A | SG Leguminosae | *Medicago truncatula* | *Quillaja saponaria* | uniform | uniform | uniform |

| | | | | | | |
|---|---|---|---|---|---|---|
| | | | | (min 63.5, max 126.0) | (min 63.5, max 126.0) | (min 63.5, max 100.0) |
| C | SG Cercis | *Cercis canadensis* | *Bauhinia tomentosa* | uniform (min 36.0, max 126.0) | | uniform (min 36.0, max 83.6) |
| C& | SG Bauhinia | *Bauhinia tomentosa* | *Cercis canadensis* | | uniform (min 46.0, max 126.0) | |
| F | CG Resin-producing clade | *Copaifera officinalis* | *Prioria balsamifera* | uniform (min 22.5, max 126.0) | uniform (min 22.5, max 126.0) | uniform (min 22.5, max 66.0) |
| G | SG Detarioideae | *Copaifera officinalis* | *Bauhinia tomentosa* | uniform (min 53.0, max 126.0) | | uniform (min 53.0, max 83.6) |
| G& | SG Resin-producing clade | *Copaifera officinalis* | *Anthonotha fragrans* | | uniform (min 53.0, max 126.0) | |
| H& | CG Amherstieae | *Afzelia bella* | *Anthonotha fragrans* | | uniform (min 46.0, max 126.0) | |
| I2 | SG Styphnolobium/Cladrastis | *Styphnolobium japonicum* | *Medicago truncatula* | uniform (min 37.8, max 126.0) | uniform (min 37.8, max 126.0) | uniform (min 37.8, max 66.0) |
| M2 | SG Robinioid clade | *Lotus japonicus* | *Medicago truncatula* | uniform (min 33.9, max 126.0) | uniform (min 33.9, max 126.0) | uniform (min 33.9, max 66.0) |
| Q | SG Acacieae/Ingeae | *Albizia julibrissin* | *Prosopis alba* | uniform (min 33.9, max 126.0) | uniform (min 33.9, max 126.0) | uniform (min 33.9, max 66.0) |
| Q2 | SG Acacia s.s. | *Acacia koa* | *Albizia julibrissin* | uniform (min 23.0, max 126.0) | uniform (min 23.0, max 126.0) | uniform (min 23.0, max 56.0) |
| Z | SG Caesalpinioideae | *Albizia julibrissin* | *Medicago truncatula* | uniform (min 58.0, max 126.0) | uniform (min 58.0, max 126.0) | uniform (min 58.0, max 83.6) |

### Table S5. Node age estimates and priors (95% HPD intervals) of nodes A-H in the different analyses.

| **Node** | **A** | **B** | **C** | **D** | **E** | **F** | **G** | **H** |
|---|---|---|---|---|---|---|---|---|
| **Clade** | **Leguminosae** | **Cerc+Detar** | **Cercidoideae** | **Detarioideae** | **Dial+Caes+Pap** | **Caes+Pap** | **Caesalpinioideae** | **Papilionoideae** |
| **Standard prior** | | | | | | | | |
| Marginal prior | 79.37 - 109.20 | 54.56 - 99.48 | 36.00 - 80.55 | 28.91 - 87.21 | 73.77 - 106.04 | 68.16 - 101.69 | 56.31 - 95.76 | 58.85 - 96.39 |
| UCLN | 65.47 - 86.45 | 57.50 - 80.75 | 36.00 - 53.97 | 25.47 - 42.98 | 63.51 - 84.73 | 60.64 - 81.67 | 54.11 - 74.49 | 55.19 - 73.58 |
| RLC | 73.46 - 81.18 | 68.06 - 75.69 | 39.34 - 46.74 | 31.52 - 36.43 | 69.77 - 77.35 | 68.05 - 75.45 | 55.76 - 63.75 | 49.05 - 54.38 |
| strict clock | 66.94 - 69.55 | 60.45 - 63.87 | 36.00 - 36.66 | 26.25 - 28.71 | 65.60 - 68.22 | 64.90 - 67.48 | 56.01 - 59.12 | 56.89 - 59.47 |
| FLC 3 clocks | 65.99 - 68.85 | 60.79 - 64.20 | 36.00 - 36.85 | 27.69 - 30.59 | 63.77 - 66.52 | 62.78 - 65.43 | 56.09 - 59.04 | 47.39 - 50.03 |
| FLC 6 clocks | 65.74 - 68.81 | 60.70 - 64.40 | 36.00 - 36.86 | 27.53 - 30.94 | 63.53 - 66.47 | 62.57 - 65.43 | 56.10 - 59.20 | 47.24 - 49.86 |
| FLC 8 clocks | 64.63 - 67.64 | 60.24 - 64.79 | 36.00 - 52.41 | 27.00 - 49.18 | 62.72 - 65.61 | 61.86 - 64.65 | 55.53 - 58.51 | 46.98 - 49.60 |
| **Alternative prior 1 (Bruneau et al. 2008 for Cercidoideae and Detarioideae)** | | | | | | | | |
| Marginal prior | 81.60 - 110.20 | 63.87 - 103.63 | 46.00 - 85.81 | 53.00 - 90.89 | 75.00 - 106.55 | 69.90 - 103.35 | 57.41 - 97.38 | 60.69 - 98.39 |
| FLC 8 clocks | 64.81 - 67.96 | 64.06 - 67.35 | 57.35 - 63.89 | 55.38 - 63.49 | 63.08 - 65.94 | 62.16 - 64.90 | 55.73 - 58.76 | 46.96 - 49.59 |
| **Alternative prior 2 (tighter maxima)** | | | | | | | | |
| Marginal prior | 73.30 - 96.56 | 56.01 - 83.60 | 36.00 - 71.11 | 28.75 - 75.30 | 66.75 - 90.79 | 64.04 - 83.60 | 53.37 - 81.05 | 54.12 - 79.22 |
| UCLN | 66.92 - 76.45 | 60.43 - 72.34 | 36.01 - 50.55 | 39.85 - 52.84 | 63.48 - 71.28 | 61.45 - 69.38 | 54.20 - 63.50 | 47.59 - 55.25 |
| **Alternative prior 3 (reduced taxon sampling)** | | | | | | | | |
| Marginal prior | 72.85 - 106.32 | 53.45 - 95.52 | 36.00 - 78.70 | 29.28 - 85.30 | 64.01 - 100.04 | 58.03 - 91.93 | 38.95 - 83.49 | 46.50 - 86.28 |
| UCLN | 64.72 - 79.69 | 57.35 - 75.43 | 36.00 - 53.18 | 25.40 - 39.20 | 62.29 - 76.62 | 60.08 - 74.12 | 49.28 - 66.70 | 49.68 - 64.63 |

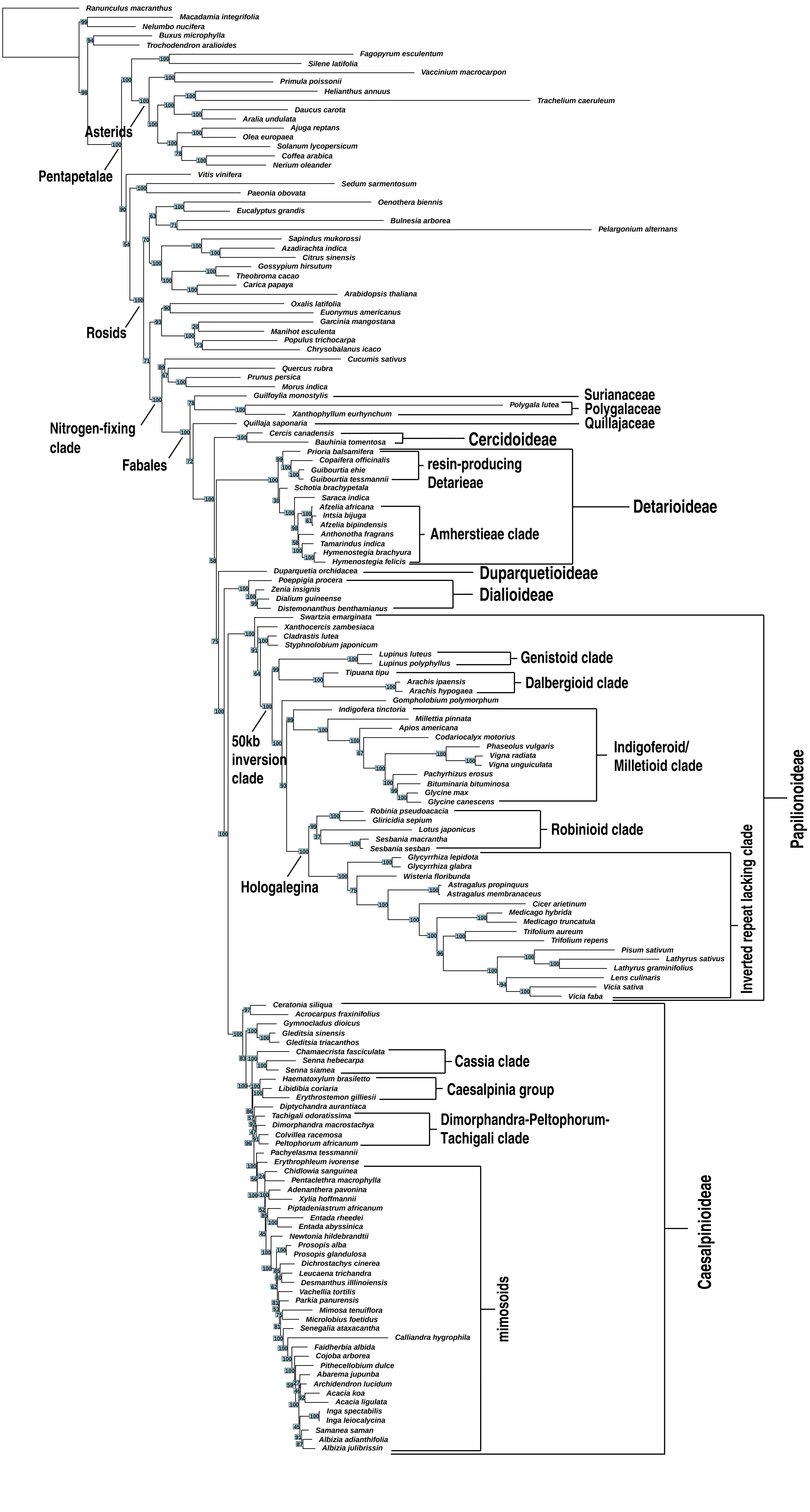


**Figure S1. ML topology as inferred by RAxML from amino acid alignment of chloroplast genes under the LG4X model.** Numbers on nodes indicate bootstrap percentages estimated from 1000 replicates.

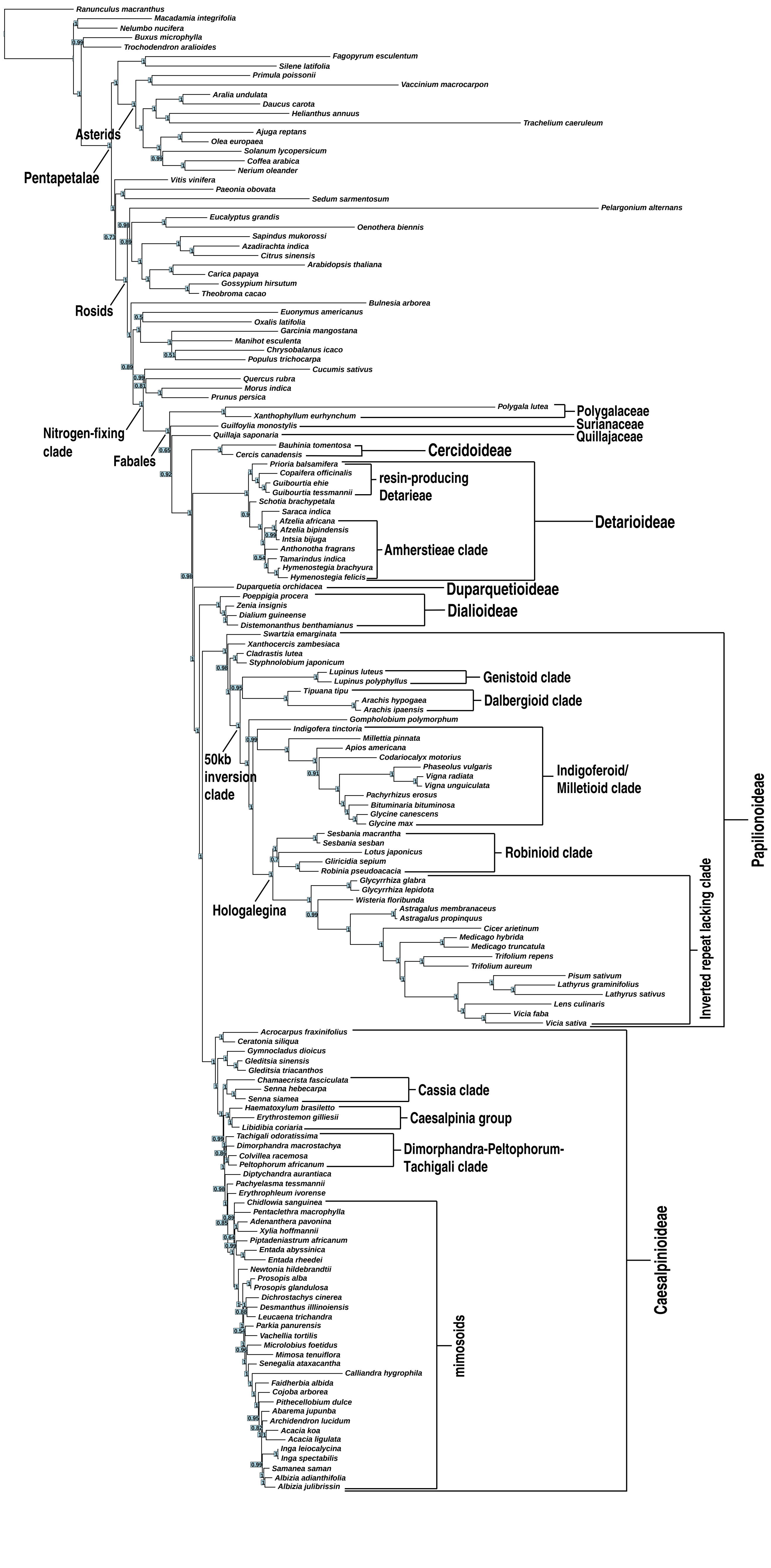


**Figure S2. Bayesian majority-rule consensus tree inferred with Phylobayes from amino acid alignment of chloroplast genes under the CATGTR model.** Numbers on nodes indicate posterior probabilities (pp) from 9000 post-burn-in MCMC cycles.

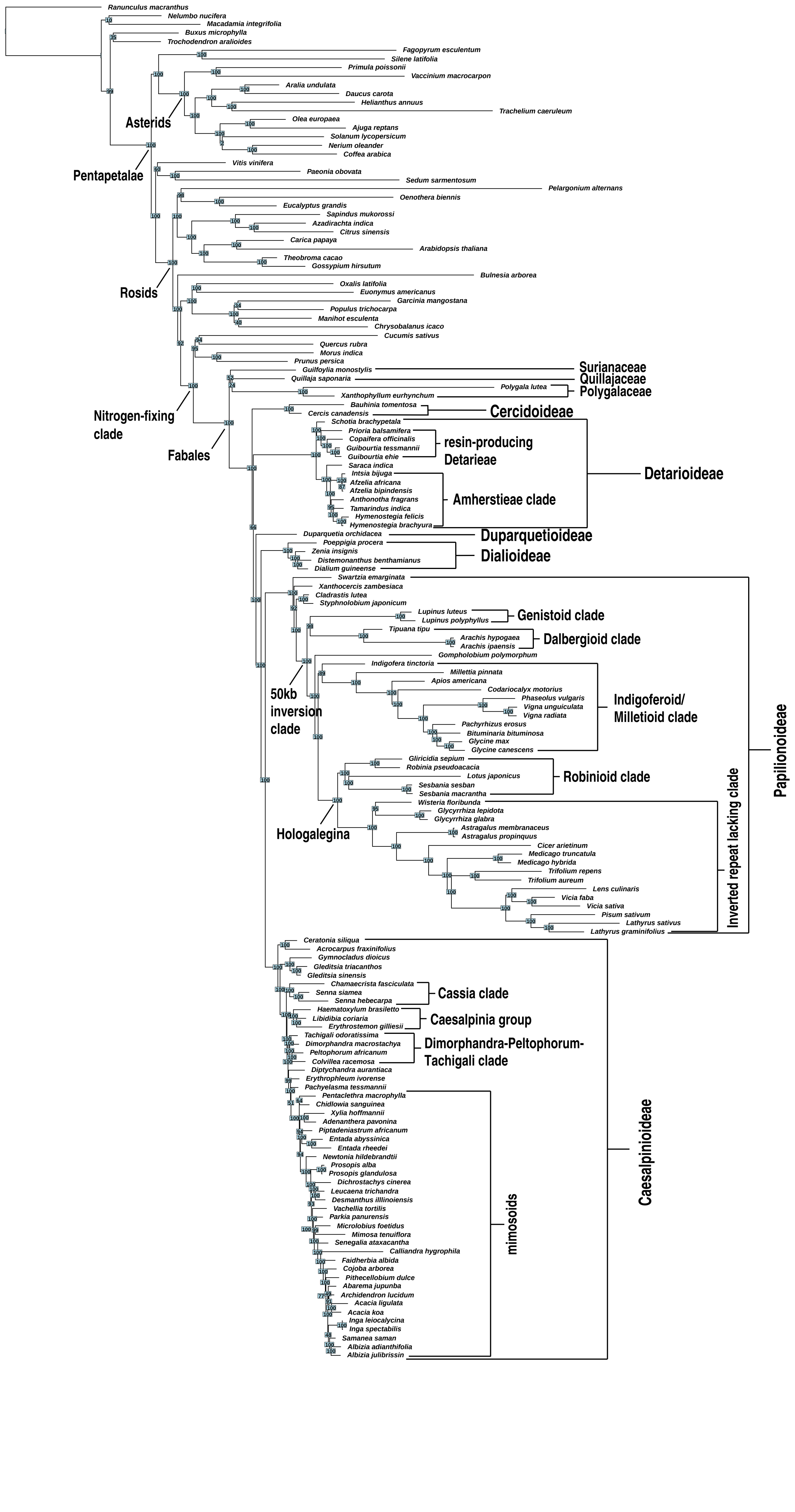


**Figure S3. ML topology as inferred by RAxML from nucleotide alignment of chloroplast genes under the GTR + G model.** Numbers on nodes indicate bootstrap percentages estimated from 1000 replicates.

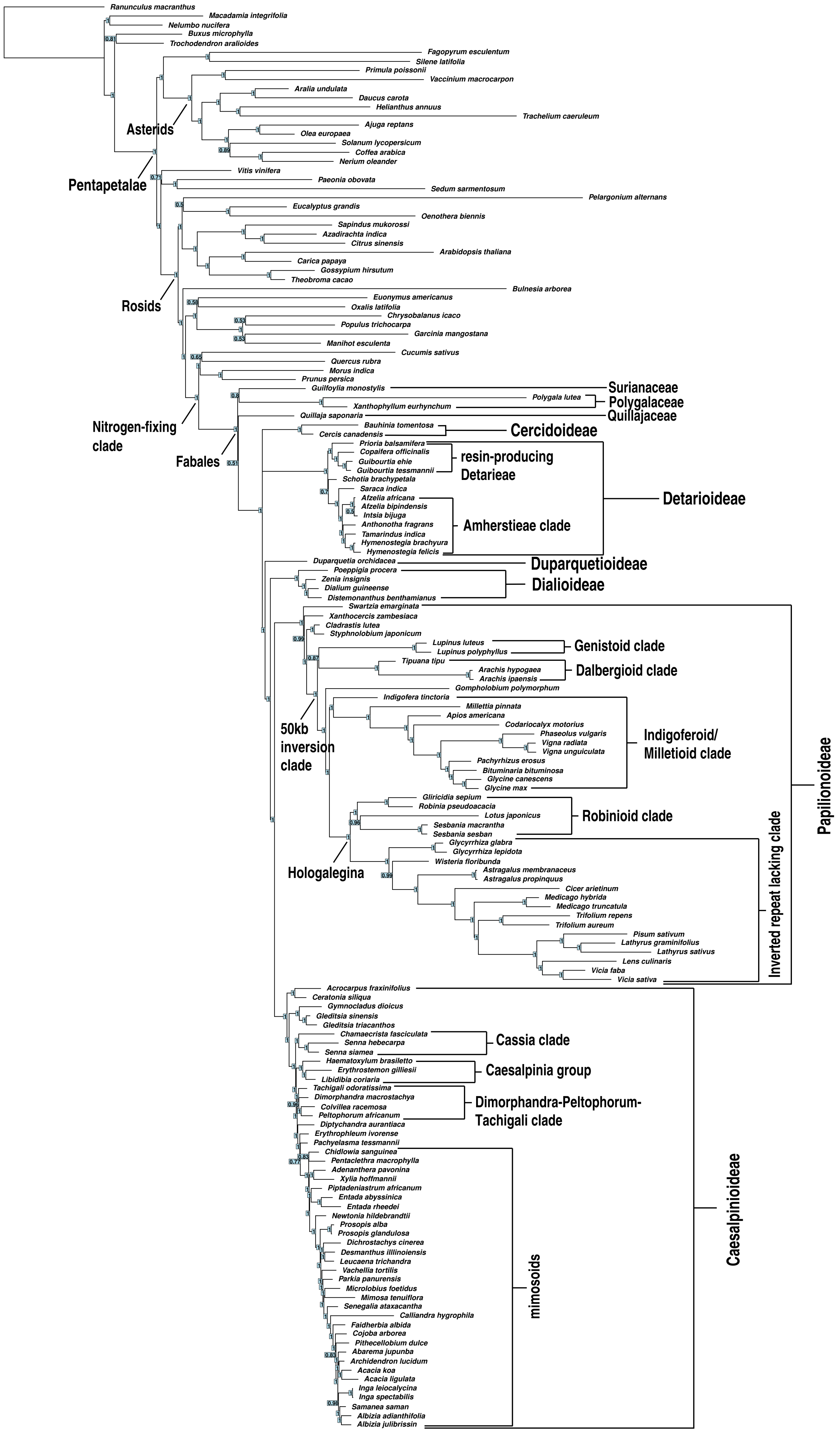


**Figure S4. Bayesian majority-rule consensus tree inferred with Phylobayes from nucleotide alignment of chloroplast genes under the CATGTR model.** Numbers on nodes indicate the posterior probabilities (pp) from 9000 post-burn-in MCMC cycles.

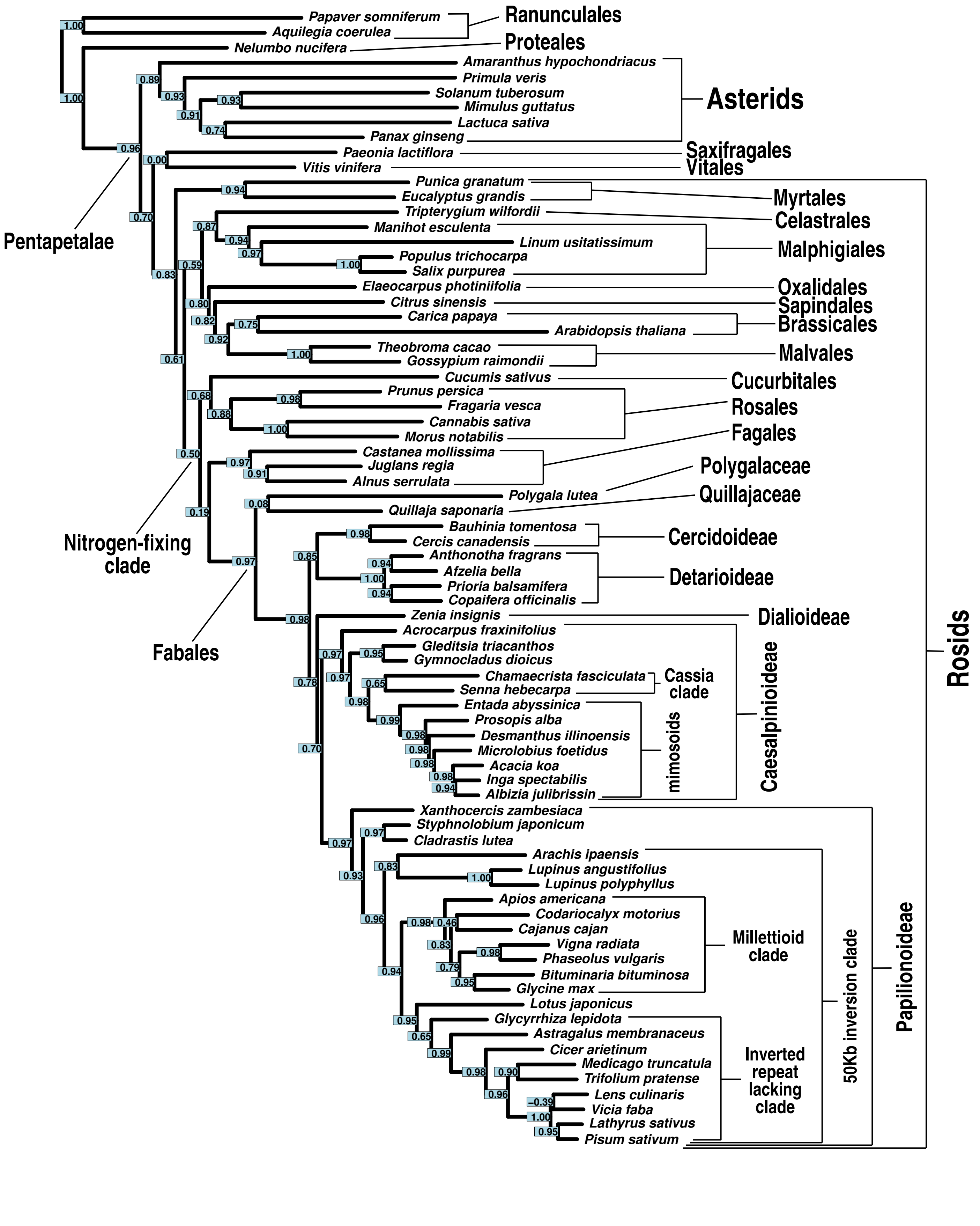


**Figure S5. ML topology as inferred by RAxML from a concatenated alignment of 1,103 nuclear genes, under the LG4X model.** Numbers on nodes indicate Internode Certainty All (ICA) values, as estimated from gene trees of the same 1,103 genes.

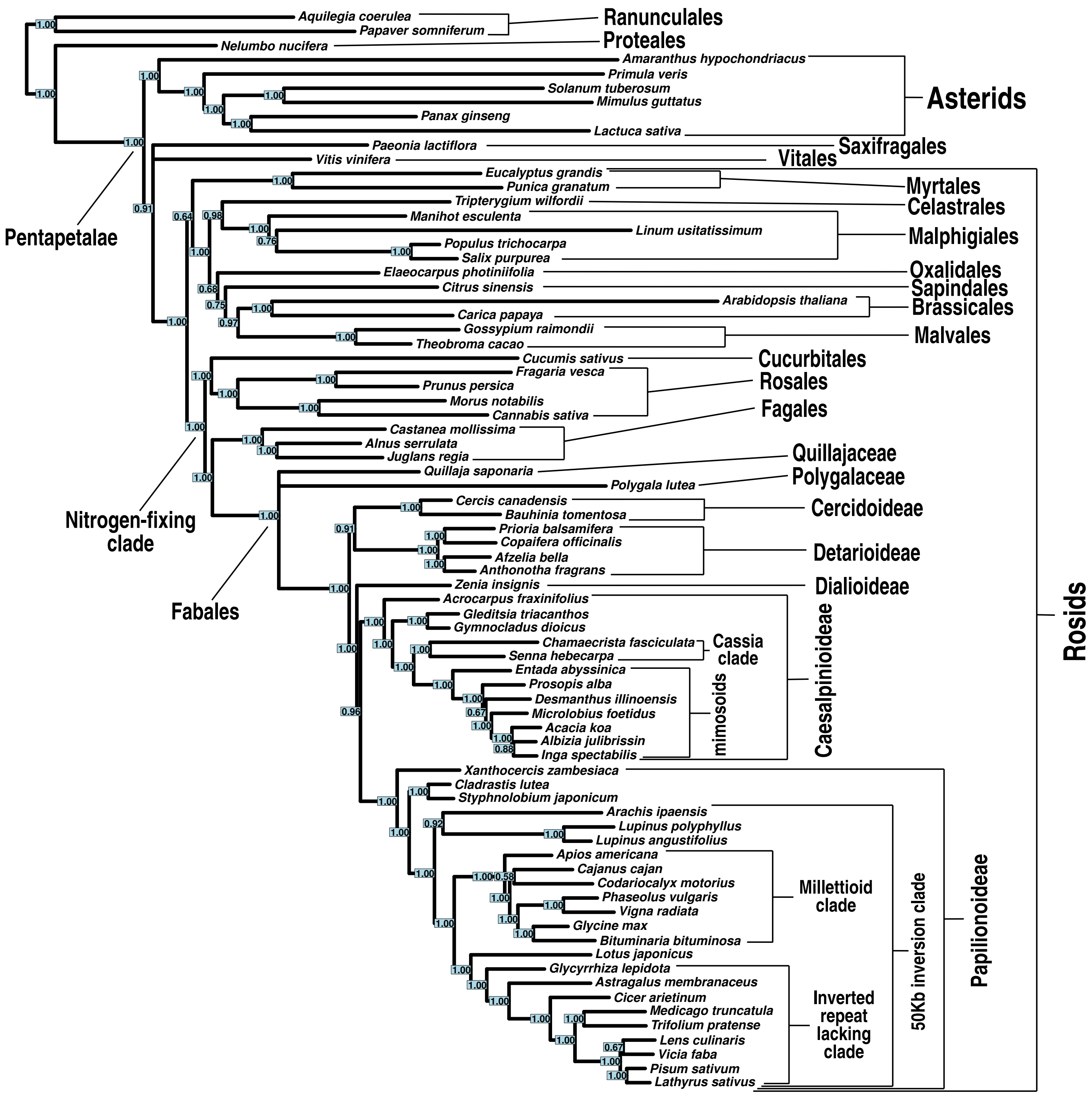


**Figure S6. Bayesian gene jackknifing majority-rule consensus tree inferred with Phylobayes from a concatenated alignment of 1,103 nuclear genes.** Numbers on nodes indicate posterior probabilities (pp), averaged over 500 posterior trees each, for 25 replicates (12,500 posterior trees in total).

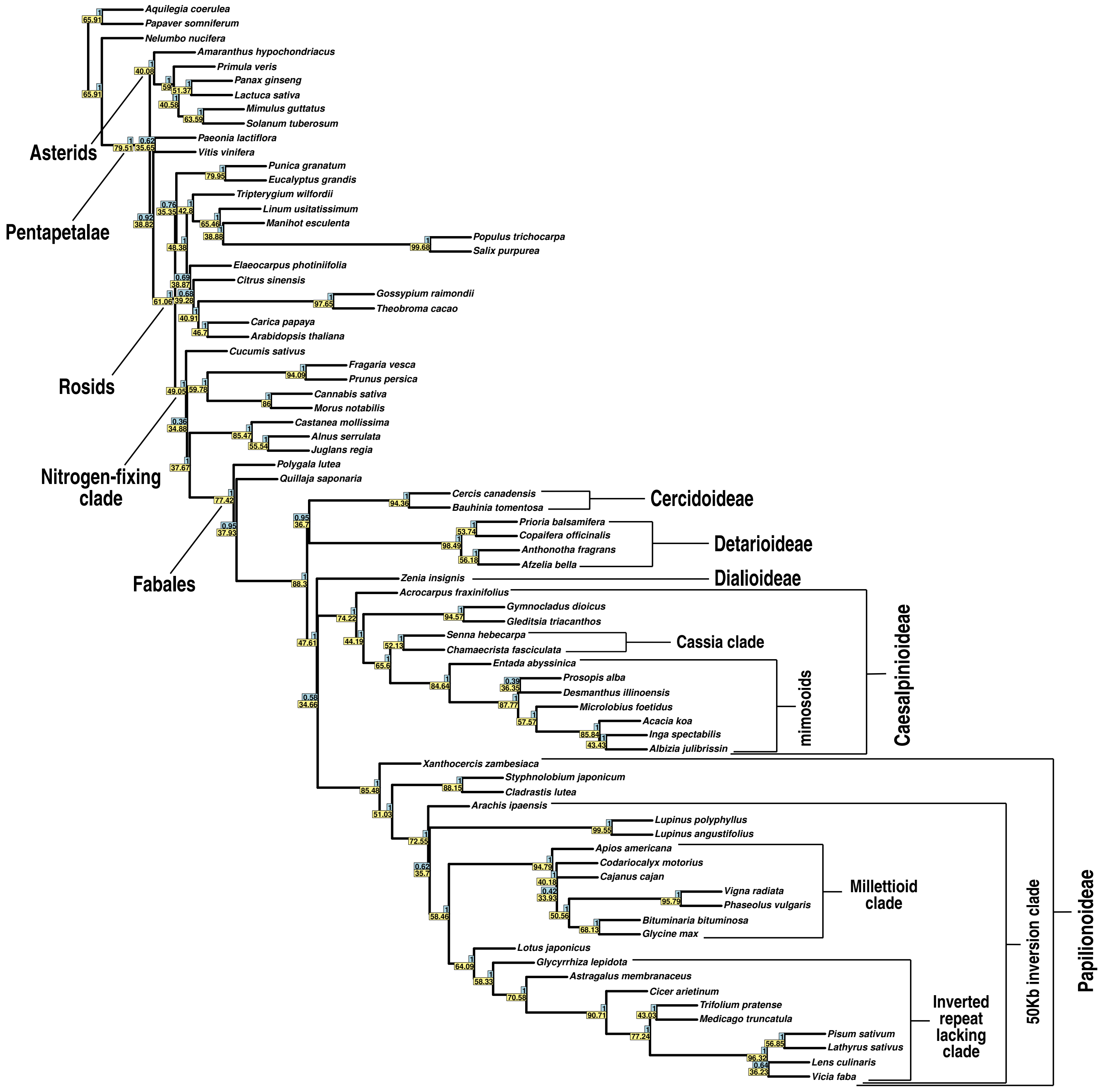


**Figure S7. Phylogeny estimated under the multi-species coalescent with ASTRAL.** Support values indicated represent local posterior probability (blue rectangles) and quartet support (yellow rectangles).

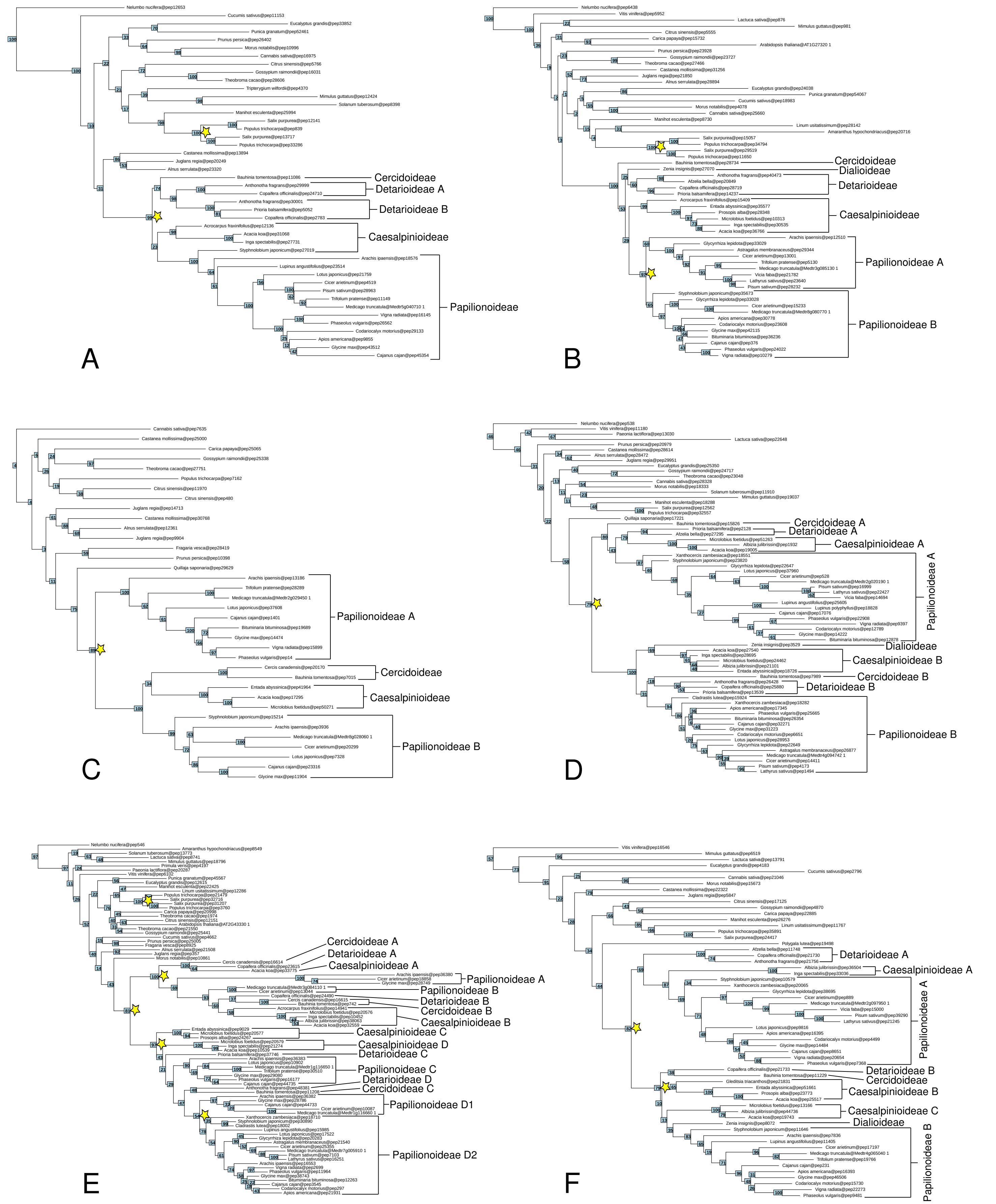


**Figure S8. Examples of homolog clusters with gene duplications in legumes that passed the bootstrap filter.** Yellow stars behind nodes indicate locations of gene duplications, numbers on nodes indicate bootstrap support. The plotted gene trees are extracted from (A) cluster3675_1rr_1rr, showing a duplication subtending Detarioideae, (B) cluster1032_1rr_1rr, showing a duplication subtending Papilionoideae, (C) cluster1248_1rr_1rr and (D) cluster2941_1rr_1rr, both with a duplication subtending the legume family. Trees for (E) cluster51_7rr_1rr and (F) cluster544_1rr_1rr show evidence of more than one duplication, including one specific to Papilionoideae in the former.

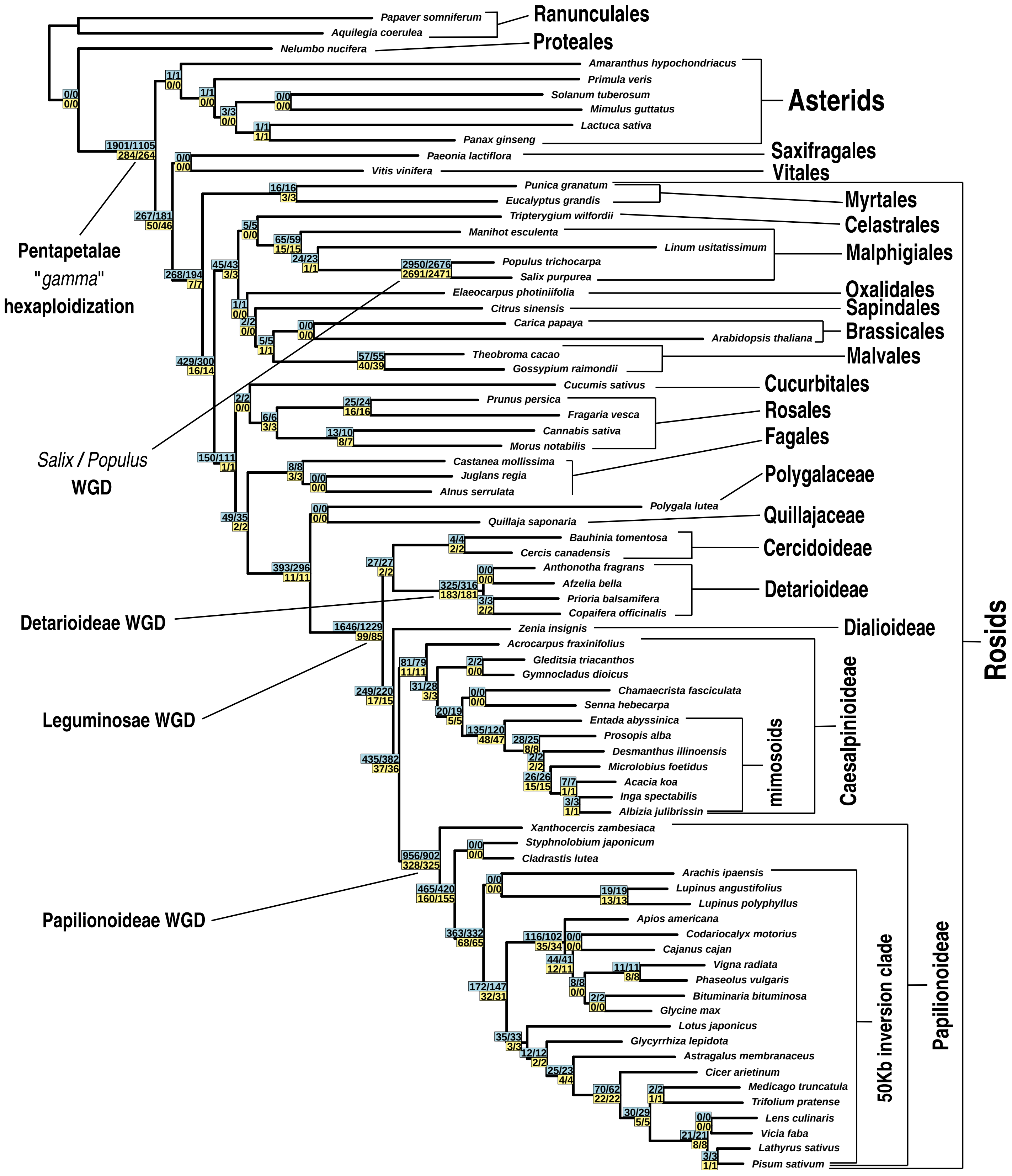


**Figure S9. Numbers of gene duplications mapped across the phylogeny.** The topology used is the ML topology of the nuclear concatenated alignment of 1,103 genes, duplications were counted from 8,038 homolog clusters. Numbers above branches (with blue background) and below branches (with yellow background) represent numbers of duplications and numbers of homolog trees with duplications, without or with a bootstrap filter of 50%, respectively.

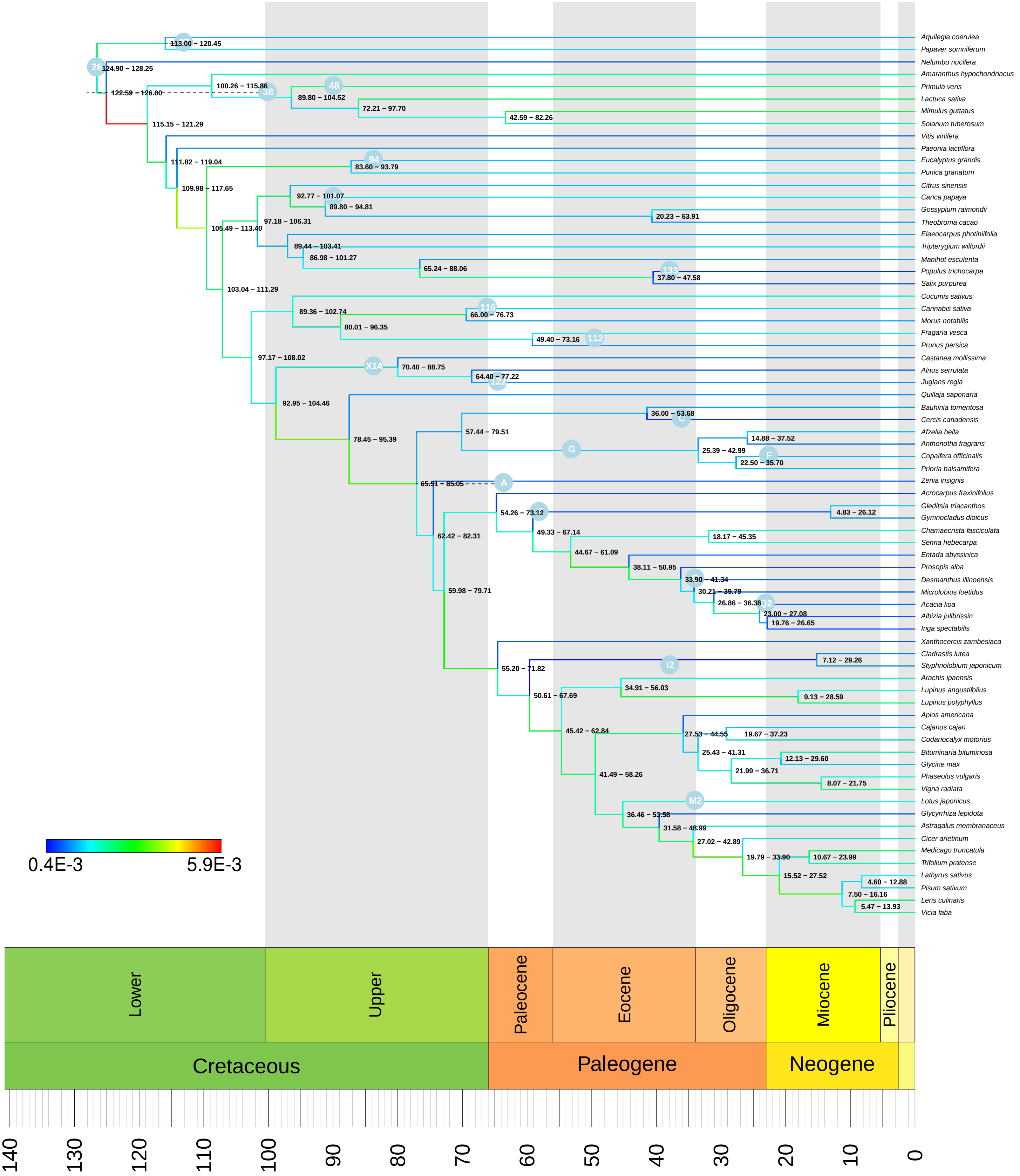

**Figure S10. Chronogram estimated under the UCLN clock model.** Numbers behind nodes indicate 95% HPD intervals. Substitution rate is indicated by colored branches, as indicated by the color legend, in substitutions per site per million years. Fossil calibrations as listed in Table 1 are indicated by blue labeled circles.

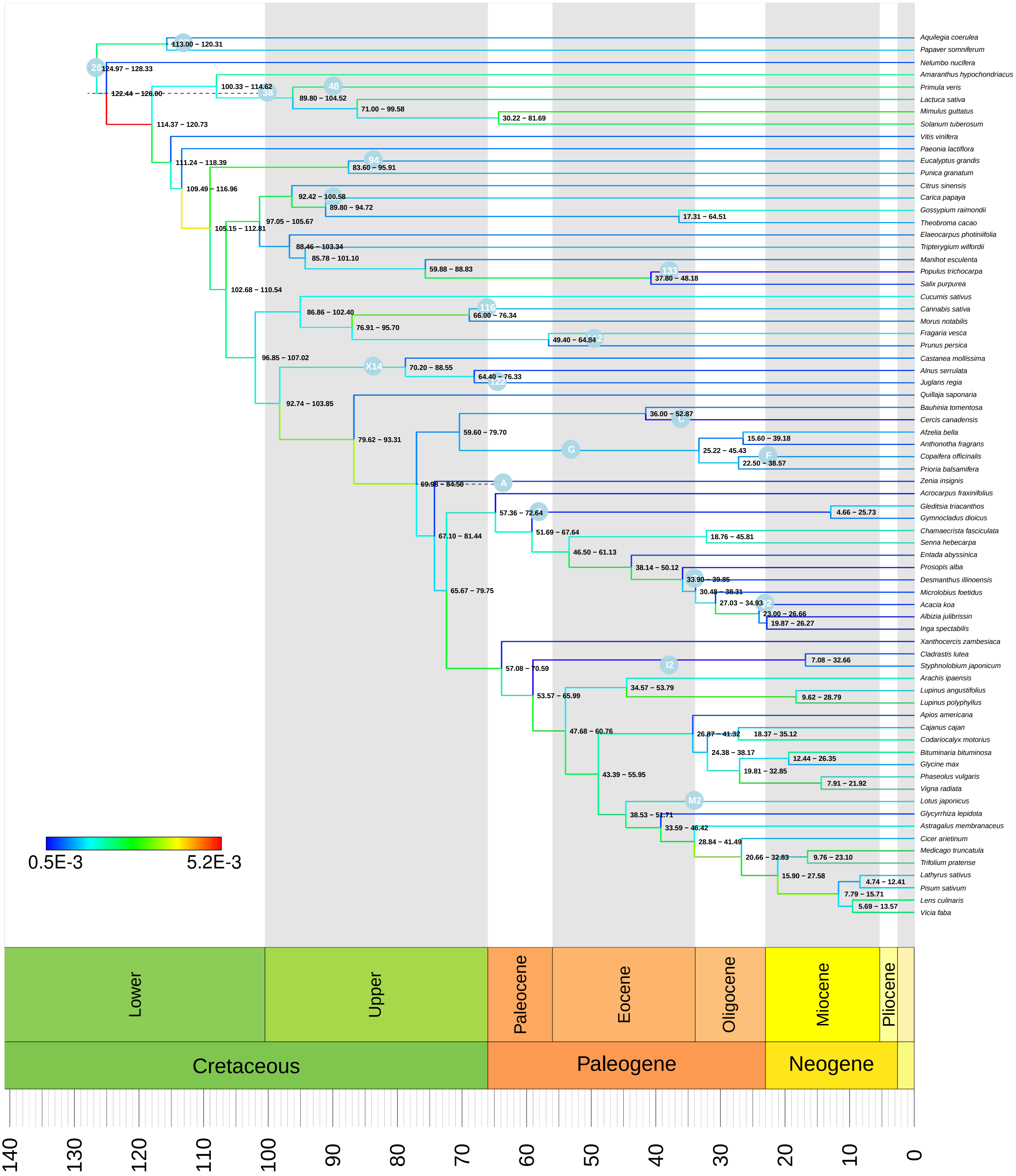


**Figure S11. Chronogram estimated under the UCLN clock model, with alternative prior 2.** Numbers behind nodes indicate 95% HPD intervals. Substitution rate is indicated by colored branches, as indicated by the color legend, in substitutions per site per million years. Fossil calibrations as listed in Table 1 are indicated by blue labeled circles.

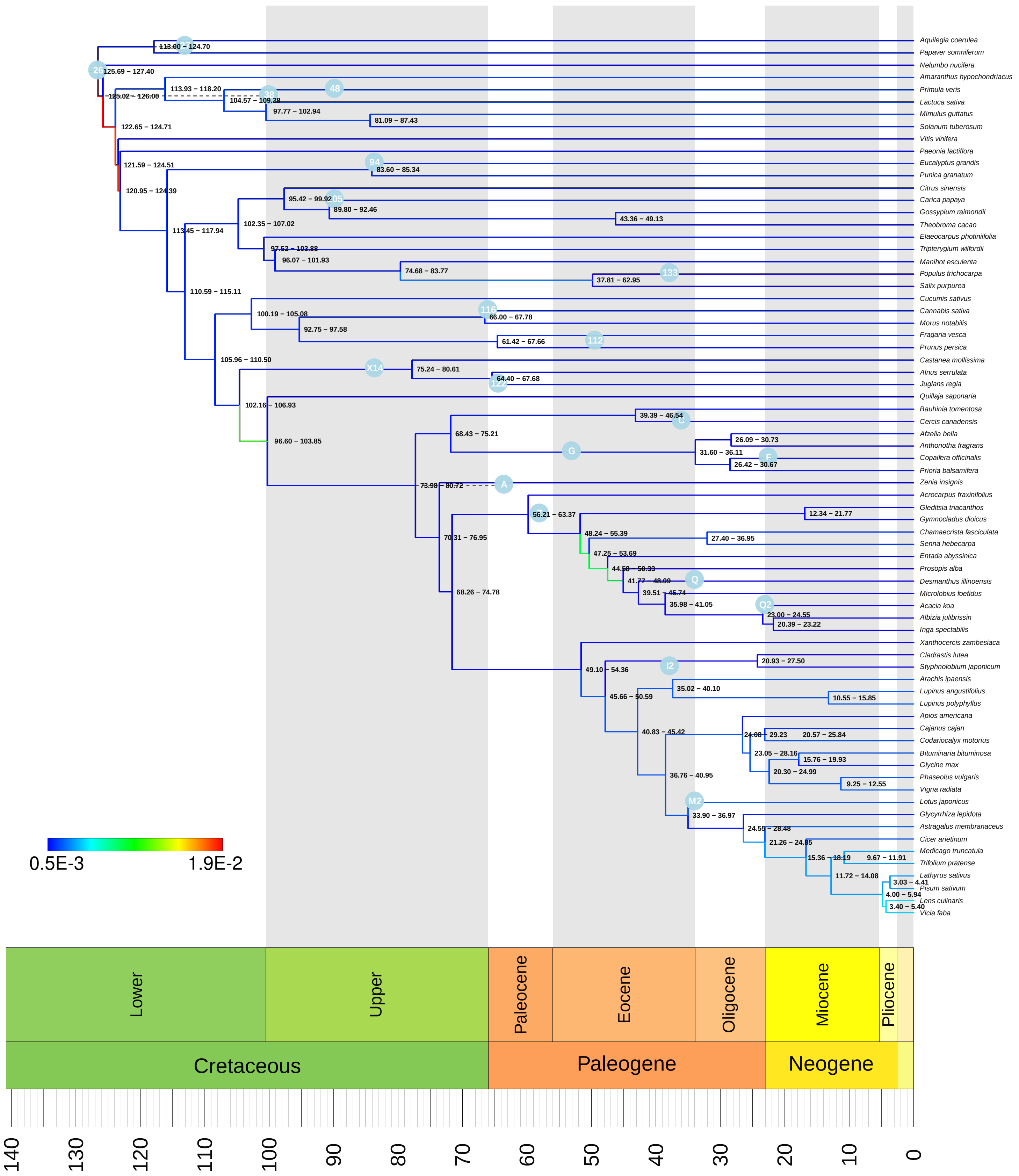


**Figure S12. Chronogram estimated under the RLC model.** Numbers behind nodes indicate 95% HPD intervals. Substitution rate is indicated by colored branches, as indicated by the color legend, in substitutions per site per million years. Fossil calibrations as listed in Table 1 are indicated by blue labeled circles.

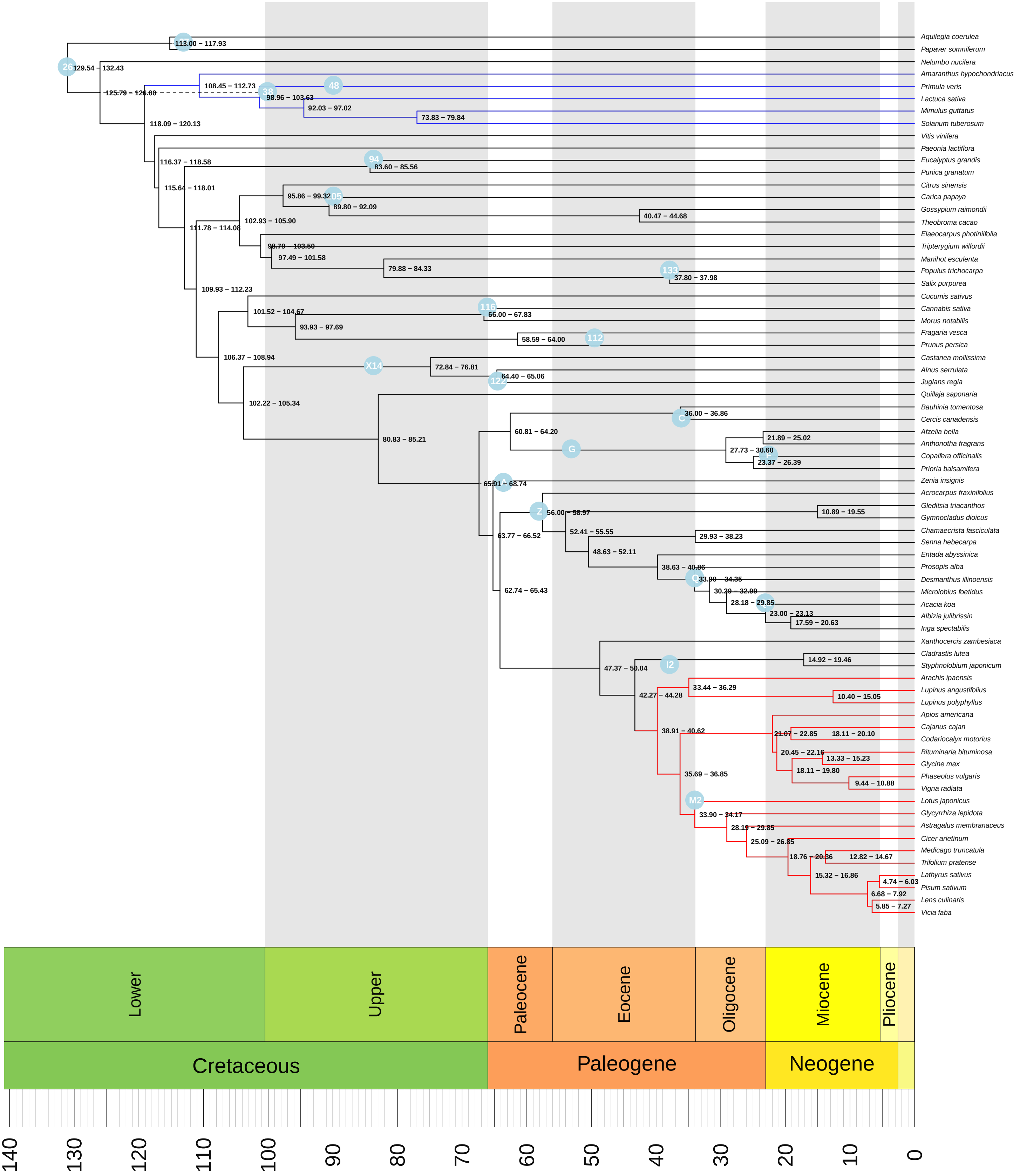


**Figure S13. Chronogram estimated under the FLC3 model.** Numbers behind nodes indicate 95% HPD intervals. Clock partitions are indicated by colored branches. Fossil calibrations as listed in Table 1 are indicated by blue labeled circles.

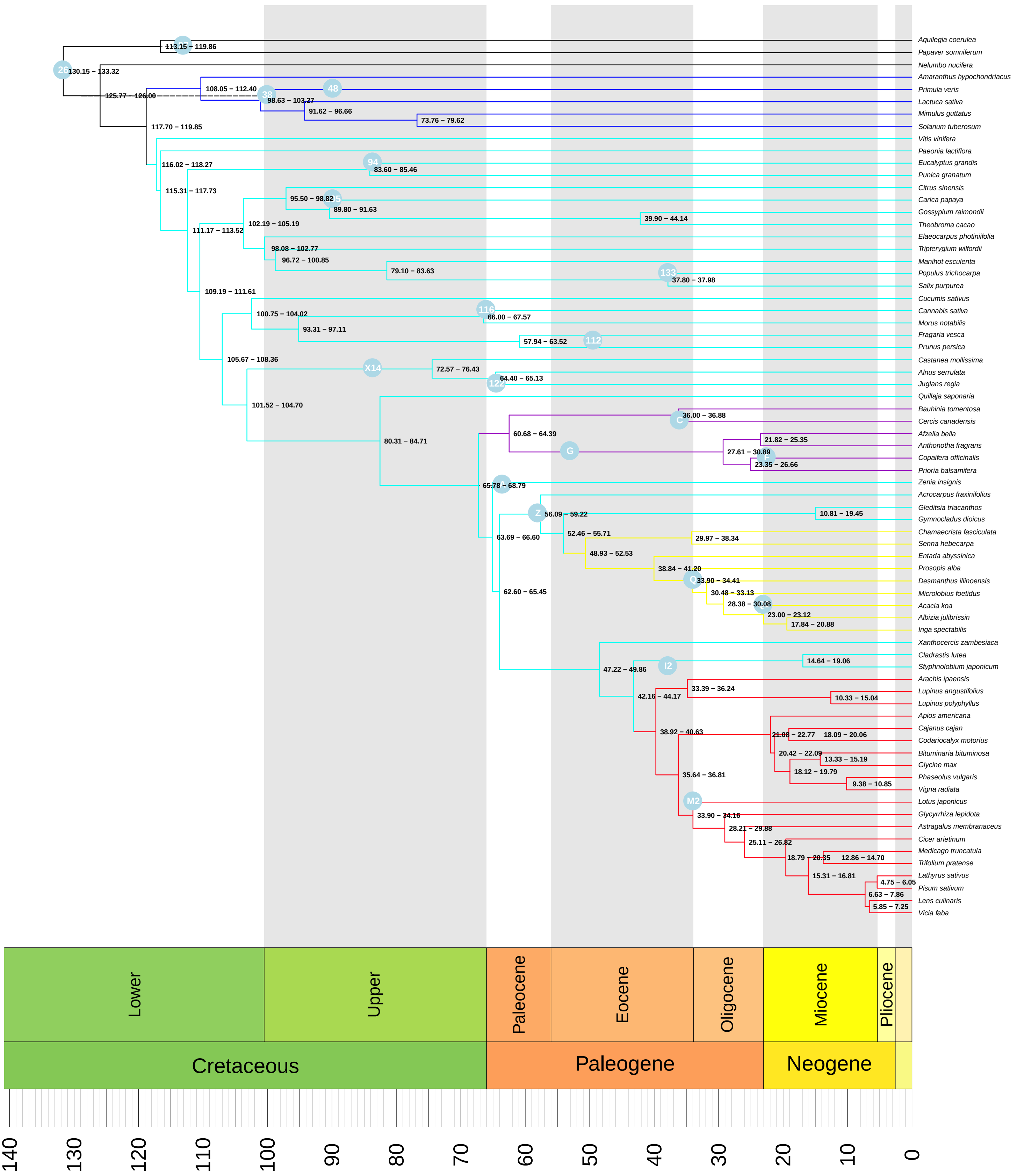


**Figure S14. Chronogram estimated under the FLC6 model.** Numbers behind nodes indicate 95% HPD intervals. Cock partitions are indicated by colored branches. Fossil calibrations as listed in Table 1 are indicated by blue labeled circles.

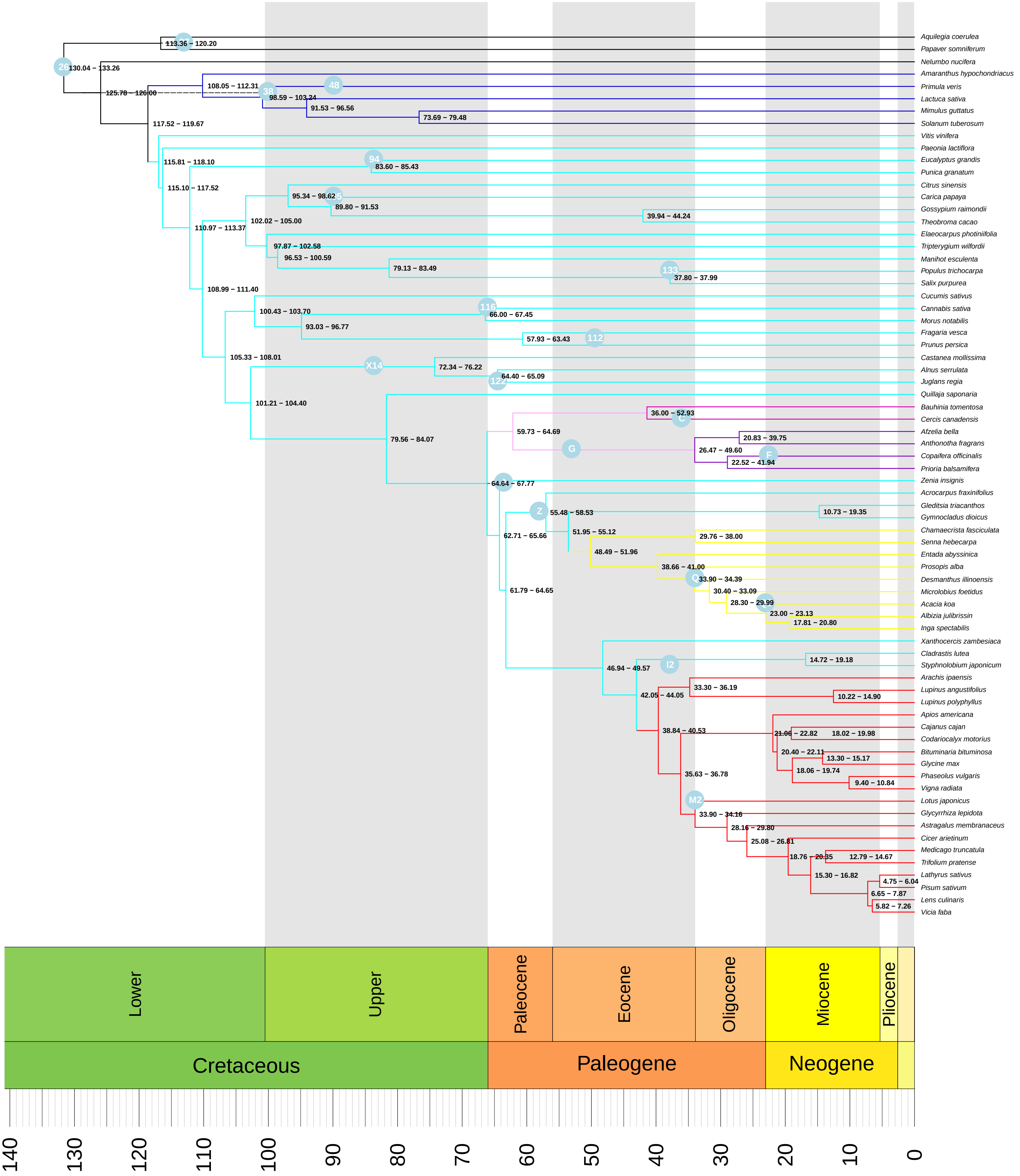


**Figure S15. Chronogram estimated under the FLC8 model.** Numbers behind nodes indicate 95% HPD intervals. Cock partitions are indicated by colored branches. Fossil calibrations as listed in Table 1 are indicated by blue labeled circles.

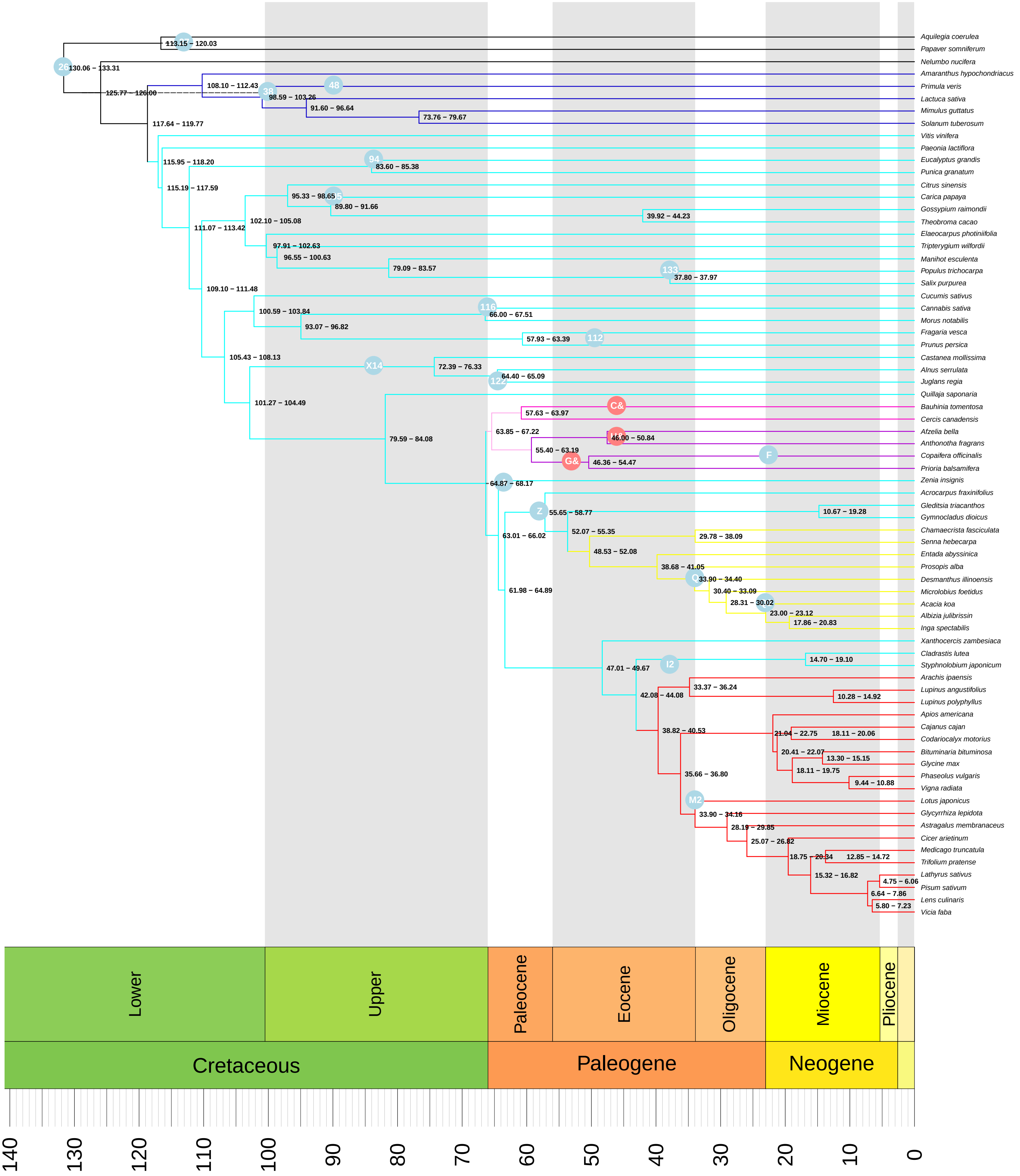


**Figure S16. Chronogram estimated under the FLC8 model, with alternative prior 1.** Numbers behind nodes indicate 95% HPD intervals. Clock partitions are indicated by colored branches. Fossil calibrations as listed in Table 1 are indicated by blue labeled circles, with alternative calibrations as red circles.

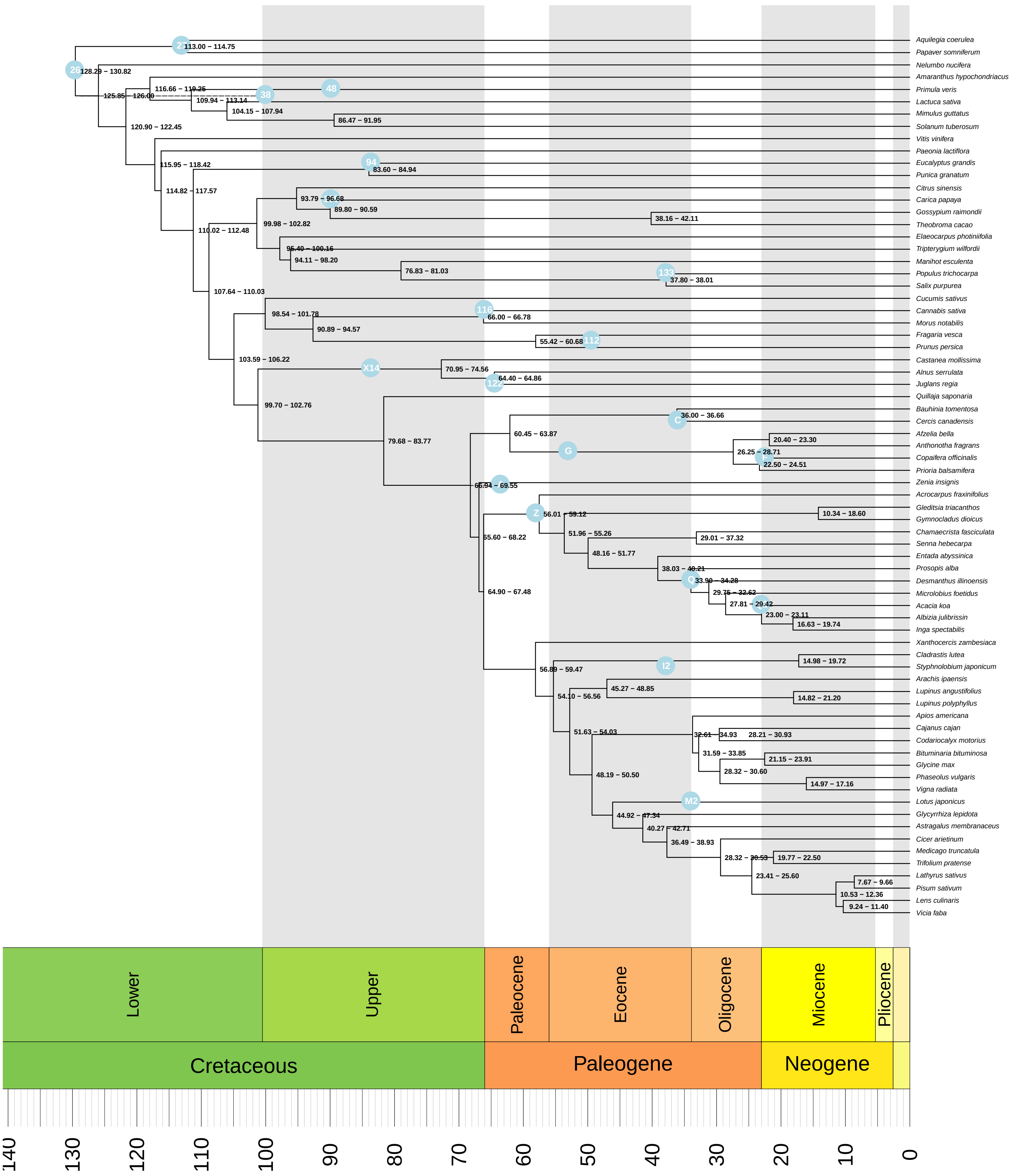


**Figure S17. Chronogram estimated under the STRC model.** Numbers behind nodes indicate 95% HPD intervals. Fossil calibrations as listed in Table 1 are indicated by blue labeled circles.

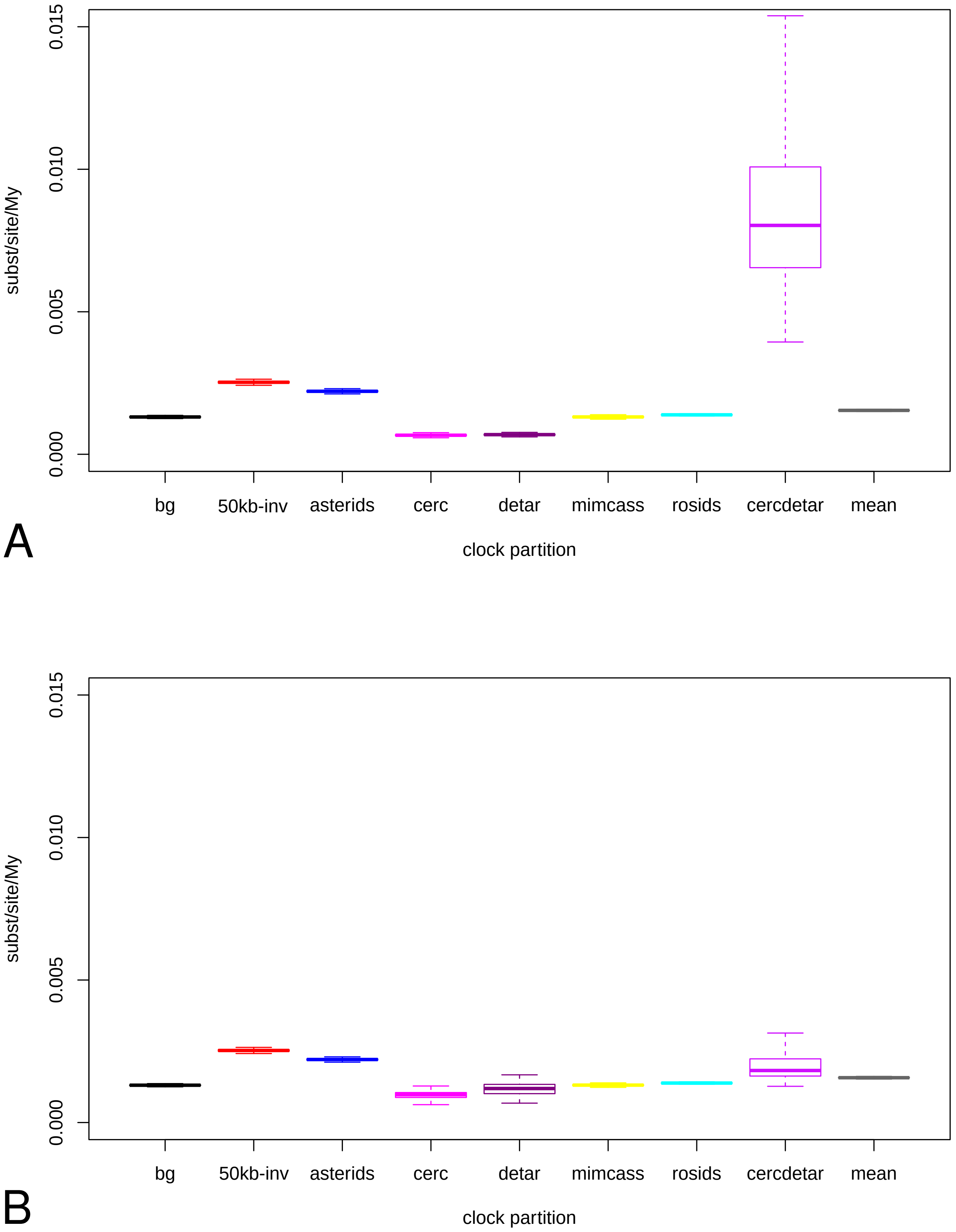


**Figure S18. Substitution rates as estimated in FLC8 analyses for the different clock partitions.** Boxplots for each partition for (A) alternative prior 1 and (B) the “normal” prior setting. Colors correspond to the partitions as shown in Figs 5, S14, S15 and S18.

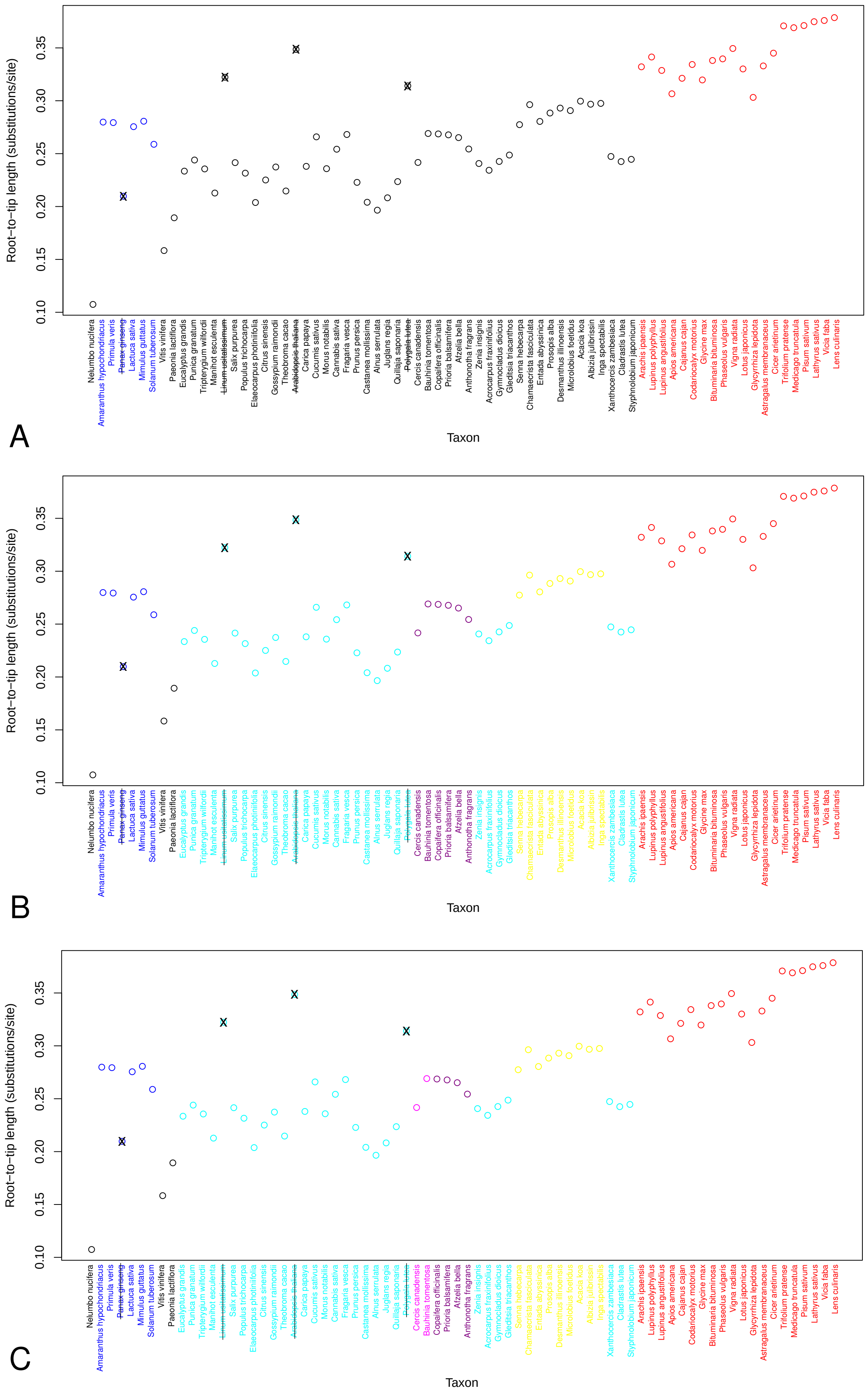

**Figure S19. Root-to-tip lengths per taxon with partitions of fixed local clocks indicated.** Pruned taxa with outlier root-to-tip lengths are indicated with an X, partitions are indicated with colors. (A) FLC3, (B) FLC6, (C) FLC8.
